# Disruption of the CSF-1-CSF-1R axis alters cerebellar microglia and is associated with motor and social interaction defects

**DOI:** 10.1101/639526

**Authors:** Veronika Kana, Fiona A. Desland, Maria Casanova-Acebes, Pinar Ayata, Ana Badimon, Elisa Nabel, Kazuhiko Yamamuro, Marjolein Sneeboer, I-Li Tan, Meghan E. Flanigan, Samuel A. Rose, Christie Chang, Andrew Leader, Hortense Le Bouris, Eric Sweet, Navpreet Tung, Aleksandra Wroblewska, Yonit Lavin, Peter See, Alessia Baccarini, Florent Ginhoux, Violeta Chitu, E. Richard Stanley, Scott J. Russo, Zhenyu Yue, Brian D. Brown, Alexandra L. Joyner, Lotje De Witte, Hirofumi Morishita, Anne Schaefer, Miriam Merad

## Abstract

Microglia, the brain resident macrophages, critically shape forebrain neuronal circuits. However, their precise function in the cerebellum is unknown. Here we show that human and mouse cerebellar microglia express a unique molecular program distinct from forebrain microglia. Cerebellar microglial identity was driven by the CSF-1R ligand CSF-1, independently of the alternate CSF-1R ligand, IL-34. Accordingly, CSF-1 depletion from Nestin^+^ cells led to severe depletion and transcriptional alterations of cerebellar microglia, while microglia in the forebrain remained intact. Strikingly, CSF-1 deficiency and alteration of cerebellar microglia were associated with reduced Purkinje cells, altered neuronal function, and defects in motor learning and social novelty interactions. These findings reveal a novel CSF-1-CSF-1R signaling-mediated mechanism that contributes to motor function and social behavior.

**Summary:** Microglia are a heterogeneous population whose identity and function are dictated by signals from their microenvironment. Kana et al. show that CSF-1 signaling is critical for maintaining cerebellar microglial transcriptional identity and homeostasis, and that altering the CSF-1 – CSF-1R axis leads to motor and behavioral defects.

## Introduction

Microglia, the resident macrophages of the central nervous system (CNS), form a continuous cellular network throughout the CNS, that controls brain immune and inflammatory responses and contributes to the regulation of neuronal homeostasis (Li and Barres, 2018). Recent studies have revealed that the brain microenvironment is heterogeneous throughout the various brain regions, and that differences in neuronal subtypes, metabolism and function could influence or be influenced by the microglia that reside in these regions (Ayata et al., 2018; Grabert et al., 2016; Tay et al., 2017), hence the need to study microglia in their regional context.

The cerebellum is an ancient part of the CNS known to control motor coordination and function (Manto et al., 2012), and is increasingly appreciated as an important regulator of higher cognitive functions (Wagner et al., 2017). However, most studies on neuronal–microglial interactions have focused on the forebrain (Parkhurst et al., 2013a; Schafer and Stevens, 2015; Squarzoni et al., 2014), and very little is known about the regulation of the microglial network in the cerebellum, prompting us in this study to probe the mechanisms that regulate microglia homeostasis in the cerebellum.

Signaling through the colony stimulating factor-1 receptor (CSF-1R) is required for microglia homeostasis (Stanley and Chitu, 2014), as suggested by the profound microglia depletion observed in *Csf1r* deficient mice (Dai et al., 2002; Ginhoux et al., 2010), but surprisingly, mice that lack the main CSF-1R ligand, CSF-1 (Ginhoux et al., 2010; Yoshida et al., 1990), show only a moderate reduction of microglia (Wegiel et al., 1998). This discrepancy was explained in subsequent reports by the identification of an alternate CSF-1R ligand, named interleukin-34 (IL-34) (Lin et al., 2008), which is produced by CNS populations in the forebrain and shown to control microglial maintenance in the forebrain, but not in the cerebellum (Greter et al., 2012; Stanley and Chitu, 2014). These results prompted us to revisit the distribution of CSF-1R ligands in the cerebellum and their contribution to cerebellar microglial maintenance.

Using a mouse model in which CSF-1 was specifically depleted from Nestin^+^ cells, along with extensive profiling of microglia in different brain regions of the mouse and human brain, we found that the CSF-1R ligand, CSF-1, shaped cerebellar microglia identity and was uniquely required for microglial homeostasis in the cerebellum. Strikingly, disruption of cerebellar microglia in CSF-1-deficient brains was associated with profound alterations of Purkinje cell (PC) morphology and function, and with motor learning and social interaction defects.

## Results

### IL-34 and CSF-1 have non-overlapping distribution in the forebrain and cerebellum, where they shape microglia numbers and phenotype

NextGen sequencing of microglia isolated from different human brain regions revealed that human cerebellar microglia expressed a transcriptional program distinct from that of forebrain microglia (Table S1 and Fig. 1, A and B), and were particularly enriched in metabolism and energy genes (*COX7B, ATP5A1, PSMD9*), a conserved functional network previously described in mouse cerebellar microglia (Ayata et al., 2018; Grabert et al., 2016). Many orthologous genes more highly expressed in human cerebellar compared to forebrain microglia were also more highly expressed in mouse cerebellar microglia (*NUPR1, EMP3, CDR1, LGALS3, APOE*) (Fig. 1 B), suggesting that similar tissue factors might imprint the cerebellar microglial program in human and mouse brains.

**Figure 1.**
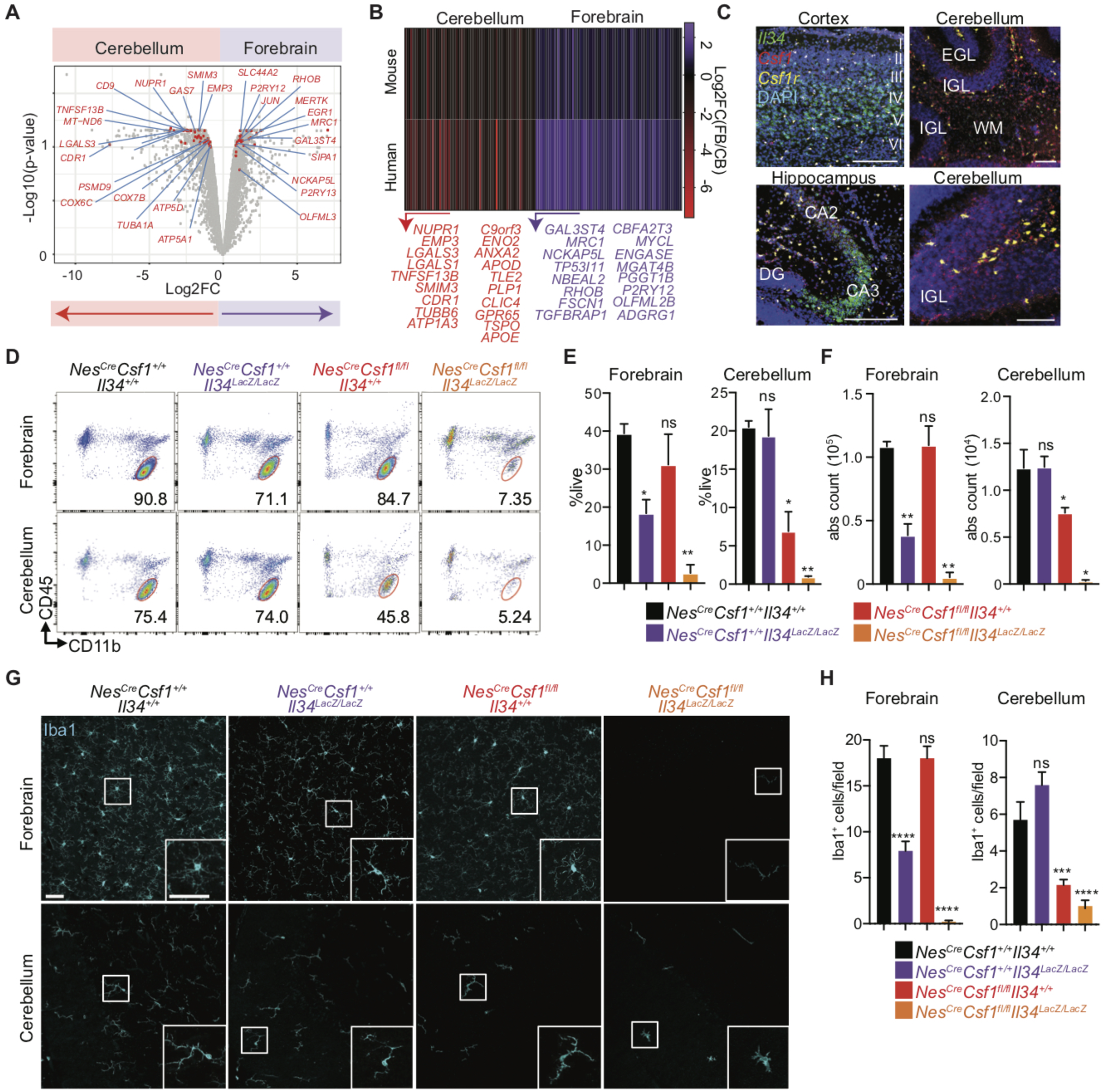
Cerebellar microglia depend on CSF-1 growth factor for their maintenance in tissue. (A) Volcano plot showing select differentially expressed genes (DEGs) between human cerebellar and forebrain (superior temporal gyrus) microglia. *n*=2 matched patient samples/brain region. (B) Heat map and representative genes of adult human and mouse orthologous genes with conserved fold change in cerebellar and forebrain cortical microglia. (C) Representative images from single-molecule RNA *in situ* hybridization for *Il34, Csf1*, and *Csf1r* in wild type P8 cerebral cortex, hippocampus, and cerebellum. (I-VI): cortical layers. (CA): cornu ammonis, (DG): dentate gyrus, (EGL): external granular layer, (IGL): internal granular layer, (WM): white matter. Scale bars: 100 µm. (D to F) Representative flow cytometric plot (D) and quantification of the percent of live (E) and absolute count (F) of forebrain and cerebellar microglia (defined as doublet^−^DAPI^−^CD11b^+^CD45^int^). *n=3-7* adult mice/group. Data are pooled from five independent experiments. (G) Representative confocal images of Iba1 (cyan) immunofluorescence stainings. Images representative of *n=3* adult mice/group. Scale bar: 20 µm. (H) Quantification of Iba1^+^ microglia/field in cortex and cerebellum. *n=3* adult mice/group, *n=3-4* fields/region/mouse. Data are a representative of two independent experiments. DEGs in (A) defined as Benjamini-Hochberg adjusted p-value<0.1. Conserved genes in (B) defined as: one to one orthologues expressing > 5 TPM in all replicates of at least one brain region in both species, and having an unadjusted p-value < 0.05 and a log_2_FC > 0.5 in the same direction in both species (log_2_FC > 0.75 in the same direction for listed genes). Graphs show mean ± SEM. *\*P≤0.05, **P≤0.01, ***P≤0.001, ****P<0.0001* using multiple student t tests (E and F) or ANOVA with Tukey’s post-hoc test (H).

Although IL-34 and CSF-1 signal through the same CSF-1R and can both promote macrophage survival, they have distinct distributions in the brain (Stanley and Chitu, 2014). Using RNA *in situ* hybridization (Fig. 1 C) and cell type-specific translating ribosome affinity purification (TRAP) (Fig. S1 A), we found that *Il34* expression dominated in forebrain regions, such as the cerebral cortex and the CA3 region of the hippocampus, while *Csf1* was most strongly expressed in the cerebellum (Fig. 1 C). The cellular source of *Il34* and *Csf1* also differed, as glial cells highly expressed *Csf1*, while *Il34* was mainly produced by neurons (Fig. S1 A). *Csf1r* expression, however, was prominent in microglia throughout the brain (Fig. 1 C). As in mouse brains, human forebrain regions mainly expressed *IL34*, whereas *CSF1* predominated in the cerebellum (Fig. S1 B).

Given the distinct spatial distribution of *Csf1* and *Il34* in the brain parenchyma, we tested how the specific depletion of each ligand affected microglia in different brain regions. Because *Csf1*^*op/op*^ mice, which bear a spontaneous CSF-1 null mutation, have bone and musculoskeletal developmental defects that may indirectly affect brain development (Yoshida et al., 1990), we crossed *Nestin*^*Cre/+*^ mice to *Csf1*^*fl/fl*^ mice (*Nes*^*Cre*^*Csf1*^*fl/fl*^) and significantly reduced *Csf1* specifically in the cerebellum, while *Il34* expression remained unaffected (Fig. S1 C). In the brain, Nestin^+^ cells mostly consist of CD45^−^ cells, including neuronal and glial progenitor cells, but not microglial cells (Fig. S1 D). Generation of *Nes*^*Cre*^*Csf1*^*fl/fl*^ mice allowed for *Csf1* depletion from all cell types deriving from neuroectodermal stem cells, including astrocytes, one of the main sources of *Csf1* (Fig. S1 A), as well as oligodendrocytes, and neurons, thus circumventing the development defects observed in *Csf1*^*op/op*^ mice.

In contrast to *Il34*^*LacZ/LacZ*^ deficient mice, which have reduced microglia in forebrain regions, including the cerebral cortex and hippocampus (Greter et al., 2012; Wang et al., 2012), *Nes*^*Cre*^*Csf1*^*fl/fl*^ mice had normal forebrain microglia frequencies and numbers, but significantly reduced microglia in the cerebellum (Fig. 1, D to H), thus mirroring the expression pattern of *Il34* and *Csf1* in their respective brain regions. Importantly, CSF-1 depletion from microglia in *Cx3cr1*^*Cre*^*Csf1*^*fl/fl*^ mice had no defects in the cerebellar microglial compartment (Fig. S1, E and F), establishing that a microglial CSF-1 source is not required for microglial homeostasis in the cerebellum, and excluding a relevant impact of transient *Cx3cr1* expression during neuronal development (Haimon et al., 2018) in *Cx3cr1*^*Cre*^*Csf1*^*fl/fl*^ mice. We also generated knock-out mice that lacked both CSF-1 and IL-34 (*Nes*^*Cre*^*Csf1*^*fl/fl*^*Il34*^*Lacz/Lacz*^) and found almost a complete depletion of microglia within all regions of the brain (Fig. 1, D to H) establishing the critical and non-redundant roles for CSF-1 and IL-34 in microglial homeostasis, as well as the significant contribution of CSF-1 to the maintenance of cerebellar microglia.

### Disruption of CSF-1 –CSF-1R signaling predominantly affects cerebellum homeostasis

NextGen profiling of the few cerebellar microglia that persisted in *Nes*^*Cre*^*Csf1*^*fl/fl*^ mice at post-natal day 8 (P8), the peak of cerebellar foliation and lengthening of cerebellar folia (Sudarov and Joyner, 2007), (Fig. 2 A and Fig. S2, A to C) revealed a significantly altered transcriptional program (Table S2 and Fig. 2, A to C), including downregulation of homeostatic and sensome genes (*Mertk, Gpr34* and *P2ry12)* (Matcovitch-Natan et al., 2016), growth and development genes (*Egr2, Spp1, Cd9*), and metabolism genes (*Hif1a, Slc37a2*) (Fig. 2, A and C). Many downregulated genes (*Apoe, Lpl, Spp1, Itgax, Gpnmb, Clec7a*) are also known to be developmentally regulated in microglia (Matcovitch-Natan et al., 2016). There was also a striking upregulation of immune response genes (*Axl, Lamp1, Clec4a1*) (Fig. 2, A to C). Downregulation of growth, development, and metabolism genes (*Klf2, Notch1, Tgf1, Dagla, Dotl1, Capn2*) also persisted in adult CSF-1 deficient mice (Fig. 2 D). In contrast, adult forebrain microglia in IL-34 deficient mice acquired a gene signature found in aging and neurodegenerative diseases, also known as a disease associated microglial (DAM) signature (Keren-Shaul et al., 2017) (Fig. 2 D). The DAM signature was more prevalent in adult IL-34 deficient brain compared to adult CSF-1 deficient brain (Fig. 2 E), consistent with prior reports that human Alzheimer’s post-mortem brains have reduced IL-34 levels particularly in the forebrain, and not the cerebellum (Walker et al., 2017). Accordingly, we found that *ex*-*vivo* exposure of neonatal microglia (Fig. 2 F) to CSF-1 or IL-34 drives a common transcriptional program, whether the microglia were isolated from the cerebellum or forebrain (Table S3, Fig. S2 D, and Fig. 2 G), particularly with CSF-1 inducing development and metabolism genes (*Klf2, Nes, Jag1*) and IL-34 inducing immune genes (*Clec4a2 Cd86, Tnfrsf1b*) (Fig. 2 G), indicating that IL-34 and CSF-1 are among the key drivers of regional microglial transcriptional profiles.

**Figure 2.**
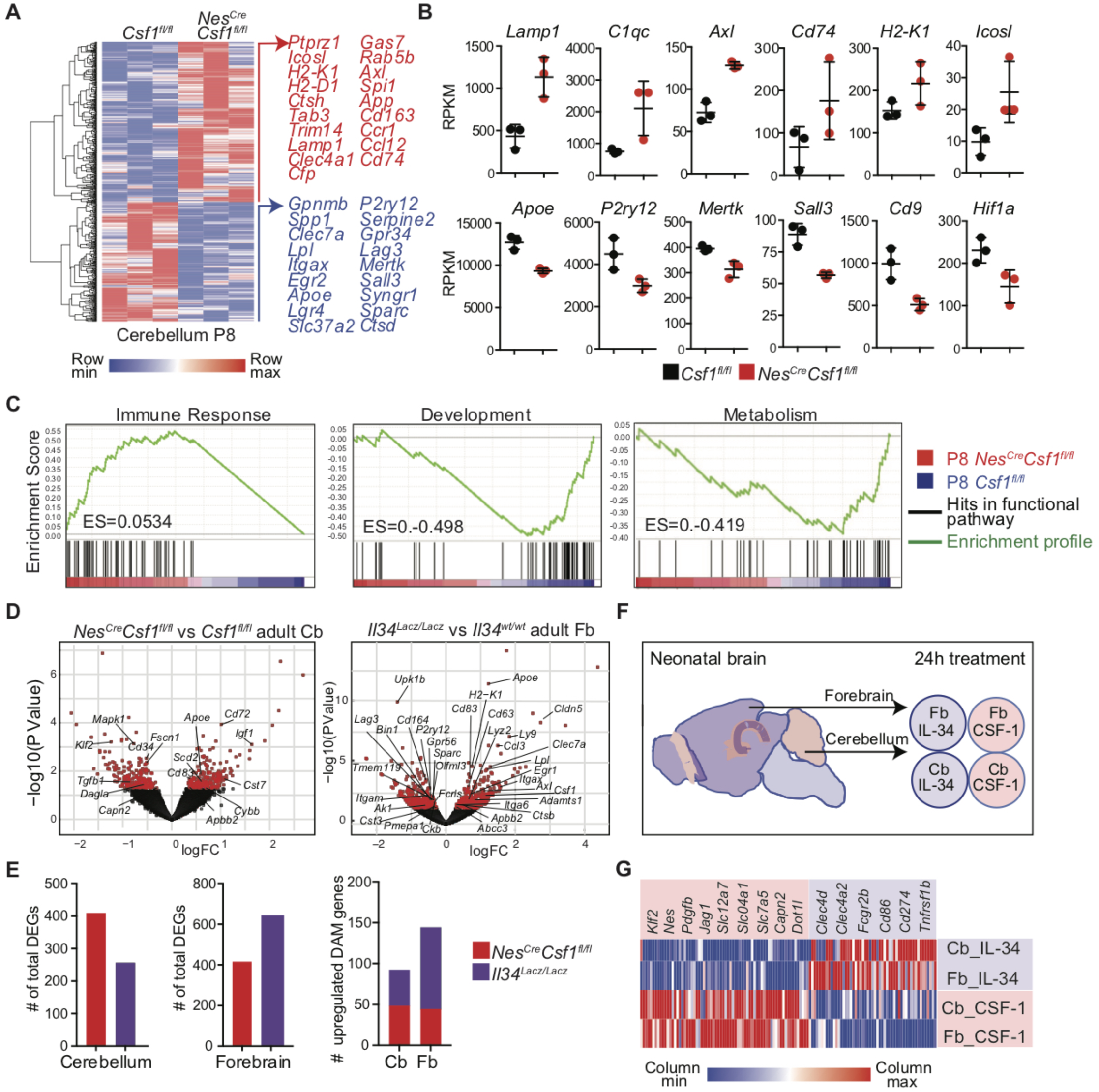
CSF-1 and IL-34 drive distinct microglial programs in the brain tissue. (A) Hierarchical clustering (left) and select representative genes (right) from 860 DEGs between P8 *Csf1*^*fl/fl*^ and *Nes*^*Cre*^*Csf1*^*fl/fl*^ cerebellar microglia. (B) Normalized expression (RPKM) of select immune response, homeostatic and metabolism genes in *Csf1*^*fl/fl*^ and *Nes*^*Cre*^*Csf1*^*fl/fl*^ cerebellar microglia. *n=3* mice/group. (C) GSEA analysis of P8 *Nes*^*Cre*^*Csf1*^*fl/fl*^ cerebellar microglia showing positive enrichment of genes involved in immune response pathways, and negative enrichment of genes involved in developmental and metabolic pathways. (D) Volcano plots showing select development, metabolism, and DAM genes in adult *Nes*^*Cre*^*Csf1*^*fl/fl*^ cerebellar microglia (left) and select homeostatic and DAM genes in adult *Il34*^*Lacz/Lacz*^ forebrain microglia (right). (E) Quantification of total DEGs in adult *Nes*^*Cre*^*Csf1*^*fl/fl*^ and *Il34*^*Lacz/Lacz*^ microglia from cerebellum (left) and forebrain (middle) and total number of upregulated intersecting genes from DAM and CSF-1R ligand deficient microglia (right). (F and G) Experimental design (F) used to generate data for heat map (G) of select representative DEGs involved in growth, differentiation, metabolism (red), and immune response (purple) pathways induced in wild type neonatal cortical and cerebellar microglia stimulated with either 100 ng/ml IL-34 or 20 ng/ml CSF-1 for 24 hours. DEGs defined as: read cut-off>10, p-value<0.05 (A and E) and LogFC(TPM)>0.25, Benjamini-Hochberg adjusted p-value<0.05 (G). Graphs show means ± SEM.

Using congenic parabiotic mice we confirmed that cerebellar microglia were maintained locally (Fig. S3 A), independent of adult hematopoiesis, as previously described for whole brain microglia (Ajami et al., 2007; Ginhoux et al., 2010). This was confirmed with flow cytometry showing no differences in *Nes*^*Cre*^*Csf1*^*fl/fl*^ cerebellar microglia expression of Tmem119, a microglial marker absent on peripherally derived monocytes and macrophages (Fig. S3 B) (Bennett et al., 2016), therefore excluding that peripherally derived monocytes accounted for the remaining microglia in *Nes*^*Cre*^*Csf1*^*fl/fl*^ cerebella. We also confirmed these cells were not replaced by perivascular macrophages, as there were no changes in key genes reported to be specifically upregulated in these border associated cells (*Mrc1, Cd36, Lyve1, Slc40a1, Hpgd*) (Goldmann et al., 2016) (Fig. S3 C). Reduction of microglia was already observed at embryonic age 13.5 (E13.5) in *Nes*^*Cre*^*Csf1*^*fl/fl*^ brain rudiments (Hoeffel et al., 2015) (Fig. S3, D and E, Fig. S5), remained low at E17.5 (Fig. S3 F) and was more prominent in the cerebellum than in the forebrain (Fig. 3 A). However, macrophages were not affected in the yolk sac, fetal liver or fetal limbs of *Nes*^*Cre*^*Csf1*^*fl/fl*^ mice (Fig. S3, G to H). In contrast, *Il34*^*LacZ/LacZ*^ mice did not exhibit reduced microglia during embryonic development as previously reported (Greter et al., 2012) (Fig. S3 I). Cerebellar microglia remained reduced throughout life in *Nes*^*Cre*^*Csf1*^*fl/fl*^ mice (Fig. 3 B). Interestingly, forebrain microglia in *Nes*^*Cre*^*Csf1*^*fl/fl*^ mice while reduced in embryos recovered by P23 (Fig. 3 B), suggesting a developmental switch in growth factor demand and an acquired dependence on the *Il34* ligand (Fig. S3, I and J).

**Figure 3.**
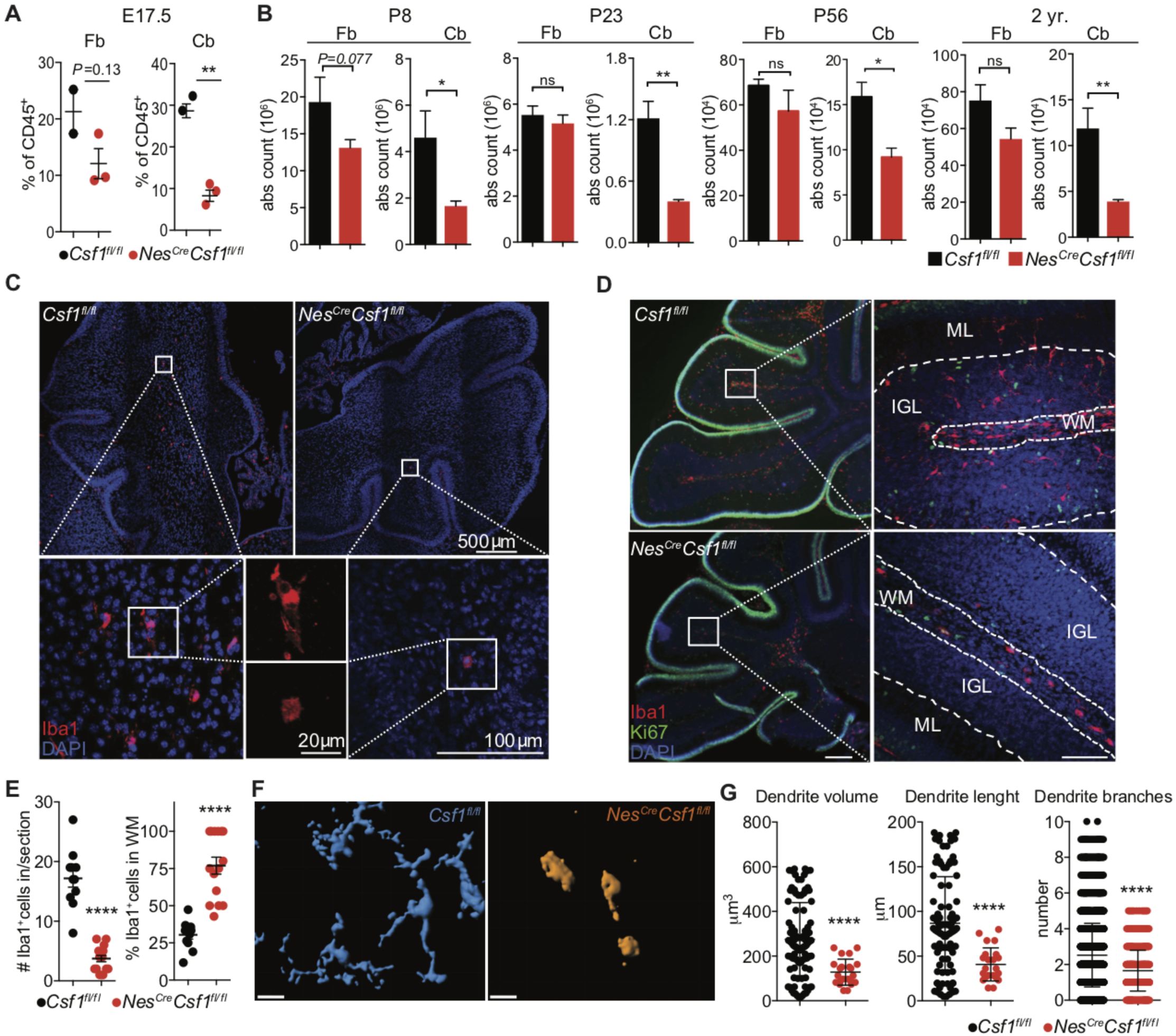
CSF-1 controls microglia morphology, spatial distribution and development during embryonic and adult life. (A) Flow cytometric quantification of doublet^−^DAPI^−^CD45^+^Ly6G^−^CD11b^lo^F4/80^hi^ microglia from E17.5 *Csf1*^*fl/fl*^ and *Nes*^*Cre*^*Csf1*^*fl/fl*^ forebrain and cerebella. *n*=3-5 mice/group. Data are pooled from two independent experiments. (B) Quantification of doublet^−^DAPI^−^CD11b^+^CD45^int^ microglia from P8, P23, P56, and 2 yr. old *Csf1*^*fl/fl*^ and *Nes*^*Cre*^*Csf1*^*fl/fl*^ forebrain and cerebellum. *n*=3-5 mice/group. Data are pooled from at least two independent experiments. (C) Representative immunofluorescence stainings of Iba1^+^ microglia in newborn (P0.5) pups, in control and *Nes*^*Cre*^*Csf1*^*fl/fl*^ cerebella. Image representative of at least n=3 mice/group from two independent litters. (D) Representative immunofluorescence stainings showing morphological and spatial distribution of Iba1^+^ microglia in P8 *Csf1*^*fl/fl*^ and *Nes*^*Cre*^*Csf1*^*fl/fl*^ cerebella. (Ki67^+^ cells): proliferating external granule cell layer, (ML): molecular layer. Scale bars: 200µm (left), 100µm (right). Image representative of n=3 mice/group of at least 3 independent experiments. (E) Quantification of Iba1^+^ microglia within P7 *Csf1*^*fl/fl*^ and *Nes*^*Cre*^*Csf1*^*fl/fl*^ cerebellar white matter and cortex (left), and percentage of microglia located only in the white matter (right). *n*=3 mice/group, 4 sections/mouse, one representative experiment of two independent experiments. (F and G) Imaris automated 3D reconstruction (F) and process quantification (G) of Iba1^+^ microglia in P8 *Csf1*^*fl/fl*^ (blue) and *Nes*^*Cre*^*Csf1*^*fl/fl*^ (orange) cerebella. Scale bar: 10µm. *n*=3 mice/group, *n*=12-34 cells/genotype. (Graphs show mean ± SEM. *\*P≤0.05, **P≤0.01, ****P<0.0001* using student’s t-test.)

Imaging of the cerebellum of *Nes*^*Cre*^*Csf1*^*fl/fl*^ mice at birth (P0.5) (Fig. 3 C) and at P6-8 (Fig. 3 D) revealed scarce microglia with a rounded, amoeboid morphology confined to the cerebellar white matter (Fig. 3 E), with significant alterations in microglial process morphology, including volume, length and number of branches (Fig. 3, F and G). Control microglia, however, were ramified and distributed throughout the white matter, the Purkinje cell (PC) layer, the molecular layer, and the developing granule cell layers (Fig. 3, C to G).

### CSF-1-deficient mice have cerebellar alterations and behavioral defects

MRI imaging of *Nes*^*Cre*^*Csf1*^*fl/fl*^ brains revealed increased brain mass, reduced brain size and ventricle volume, (Fig. S4, A and B), and increased cerebellar volume (Fig. S4 B), consistent with prior reports in *Csf1*^*op/op*^ and *Csf1r*^*-/-*^ mice (Nandi et al., 2012). Strikingly, *Nes*^*Cre*^*Csf1*^*fl/fl*^ mice had a significant reduction of PC numbers (Fig. 4 A, Fig. S4 C), which was also confirmed in CSF-1 deficient *Csf1*^*op/op*^ mice (Fig. S4 D). In contrast to previous reports (Murase and Hayashi, 2002; Nandi et al., 2012), *Csf1r* was not expressed by Calbindin^+^ (Calb^+^) PCs (Fig. S4, E and F). We confirmed that *Csf1r* was mainly expressed in microglial cells using TRAP (Fig. S4 G), as well as with RT-PCR (Fig. S4 H). As in mouse brains, human PC also lacked *CSF1R* expression (Fig. S4 I).

PCs form a continuous cellular layer within the cerebellar cortex, and their dendritic trees expand into the cerebellar molecular layer. While the main excitatory inputs of the cerebellar cortex originate from climbing and parallel fibers, the only cerebellar output is from PC axons, and their synaptic plasticity is required for successful motor learning (Nguyen-Vu et al., 2013). Imaging of *Nes*^*Cre*^*Csf1*^*fl/fl*^ PCs revealed an increase in aberrant dendrites emerging from somas (Fig. 4 B and Fig. S4, J and K), overall increased branching assessed by dendrite branching complexity quantification (Fig. 4 C), no change in dendritic spine density (Fig. 4, D and E), but decreased mean spine diameter, spine head diameter, and mean spine head volume to length ratio (Fig. 4 E), suggesting a defect in PC neural circuit formation. Climbing fiber elimination is a neurodevelopmental process ensuring that each PC ultimately receives inputs from a single climbing fiber; a process that starts in the first postnatal week (Kano et al., 2018). Developing microglia cluster below the PC layer at P7, and participate in climbing fiber elimination (Nakayama et al., 2018). At P7, *Nes*^*Cre*^*Csf1*^*fl/fl*^, PC dendrites were shorter resulting in a thinner molecular cell layer (Fig. 4 F and G), suggesting an altered and delayed PC maturation. PC soma were devoid of microglia interactions (Fig. 4 F, Fig 3 D and E) and contained more VGluT2^+^ climbing fiber puncta (Fig. 4 H and I), suggesting decreased climbing fiber elimination.

**Figure 4.**
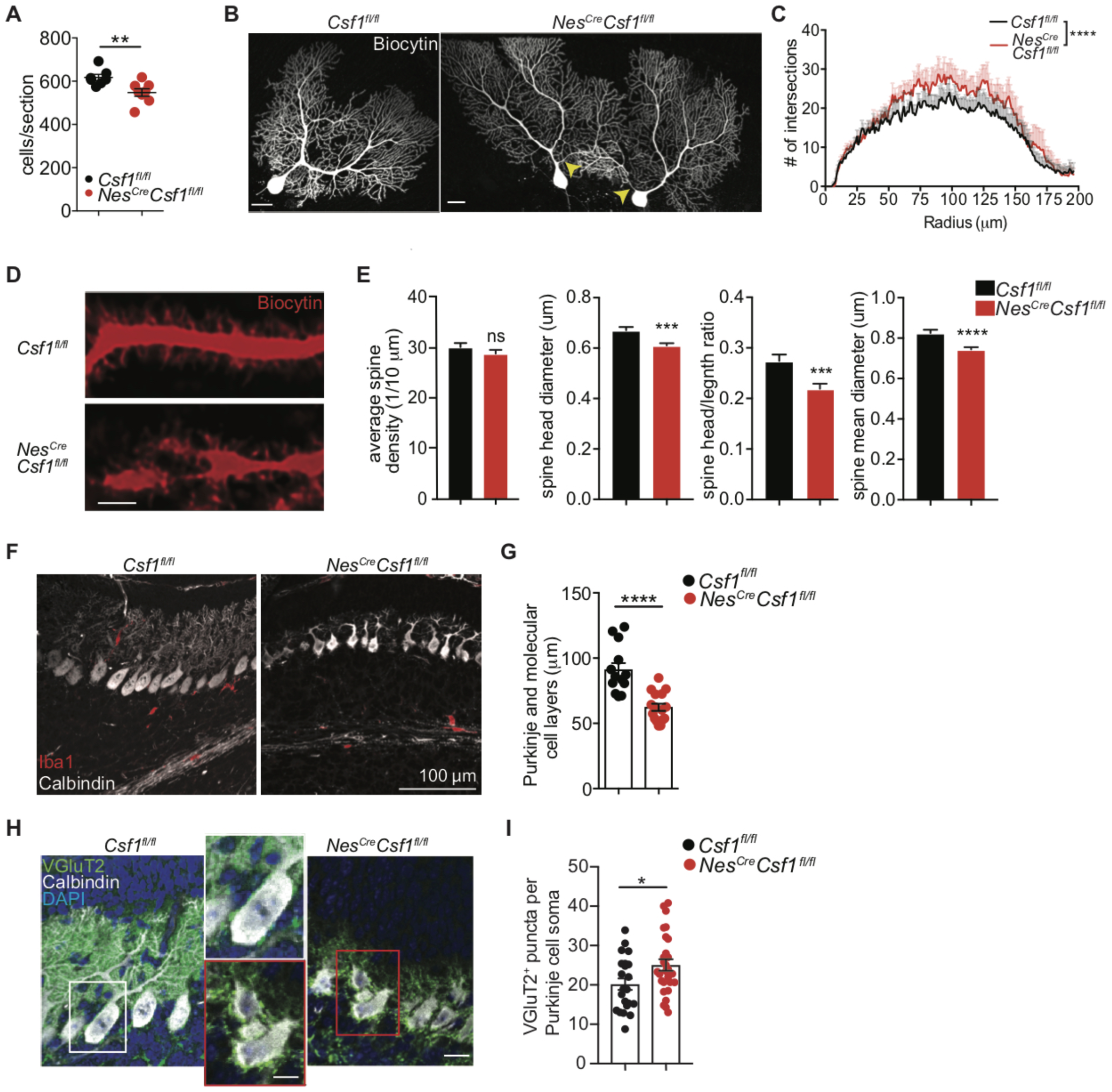
*Nes*^*Cre*^*Csf1*^*fl/fl*^ mice have morphological Purkinje cell alterations. (A) Quantification of PC numbers per sagittal section in 6 week old *Csf1*^*fl/fl*^ and *Nes*^*Cre*^*Csf1*^*fl/fl*^ cerebella, stained with H&E. *n*=4 mice/group, 1-2 sections/mouse, pooled from two independent experiments. (B-C) Representative whole cell imaging (B) and dendrite branching complexity quantification (C) of biocytin-filled and fluorophore conjugated-streptavidin stained PC. Arrows point to aberrant dendrites emanating from *Nes*^*Cre*^*Csf1*^*fl/fl*^ PC soma. Scale bar: 20µm. *n*=3 mice/genotype, *n*=7-8 cells/group, pooled from three independent patch experiments. Only dendrite branching up to 200µm radii from PC soma center was quantified, as dendrites beyond this distance may not have been reliably filled with biocytin. (D) Representative confocal images of biocytin-labeled PC dendrites and spines from *Csf1*^*fl/fl*^ and *Nes*^*Cre*^*Csf1*^*fl/fl*^ cerebella. Scale bars: 2µm. (E) Imaris quantification of PC dendritic spines in *Csf1*^*fl/fl*^ and *Nes*^*Cre*^*Csf1*^*fl/fl*^ cerebella. D and E: *n*=3 mice/genotype, *n=*3 neurons/mouse, *n=*5 dendrites/neuron, pooled from three independent patch experiments. (F and G) Representative confocal images of Calb^+^ PC and Iba1^+^ microglia (F) and quantification of PC and molecular cell layer thickness (G) in P7 *Csf1*^*fl/fl*^ and *Nes*^*Cre*^*Csf1*^*fl/fl*^ cerebella. *n*=3 mice/group, 4-6 sections/mouse. (H and I) Representative confocal images of VgluT2^+^ puncta on Calb^+^ PC soma (H) and quantification of VGluT2^+^ puncta per PC soma (I) in P7 *Csf1*^*fl/fl*^ and *Nes*^*Cre*^*Csf1*^*fl/fl*^ cerebella. *n*=3-4 mice/group, 4-6 sections/mouse. (Graphs show means ± SEM. *\*P ≤ 0.05, **P ≤ 0.01, ***P ≤ 0.001, ****P≤0.0001* using student’s t-test (A, E, G and I), and two-way ANOVA (C)).

Electrophysiological analysis of PCs in 6 week old mice showed increased spontaneous miniature excitatory post-synaptic currents (mEPSCs), with no differences in amplitude (Fig. 5, A and B), consistent with excess dendrite branching observed in the remaining PCs in these mice. Strikingly, *Nes*^*Cre*^*Csf1*^*fl/fl*^ mice showed mild signs of ataxia (Fig. S4 L), and when tested for motor learning capacities during repeated trials on an accelerating rotarod, 5-7 week old *Nes*^*Cre*^*Csf1*^*fl/fl*^ mice displayed impairment on the fourth training day (Fig. 5 C), suggesting a motor learning deficit consistent with cerebellar dysfunction. Besides its well-known role in motor coordination and function, the cerebellum is increasingly appreciated as an important regulator of higher cognitive functions (Wagner et al., 2017). We therefore performed two behavioral tests, social preference and social novelty, in *Nes*^*Cre*^*Csf1*^*fl/fl*^ mice using the 3-chamber sociability paradigm (Fig. 5 D). Both *Csf1*^*fl/fl*^ and *Nes*^*Cre*^*Csf1*^*fl/fl*^ mice showed no defects in the social preference test, both spending significantly more time interacting with the stimulus mouse over the inanimate cup (Fig. 5 E). However, *Nes*^*Cre*^*Csf1*^*fl/fl*^ mice failed to preferentially interact with a novel stimulus mouse (Fig. 5 F), indicating a deficit in social memory, a behavioral defect commonly found in mouse models of neurodevelopmental and neuropsychiatric disorders, such as autism spectrum disorders (ASD).

**Figure 5.**
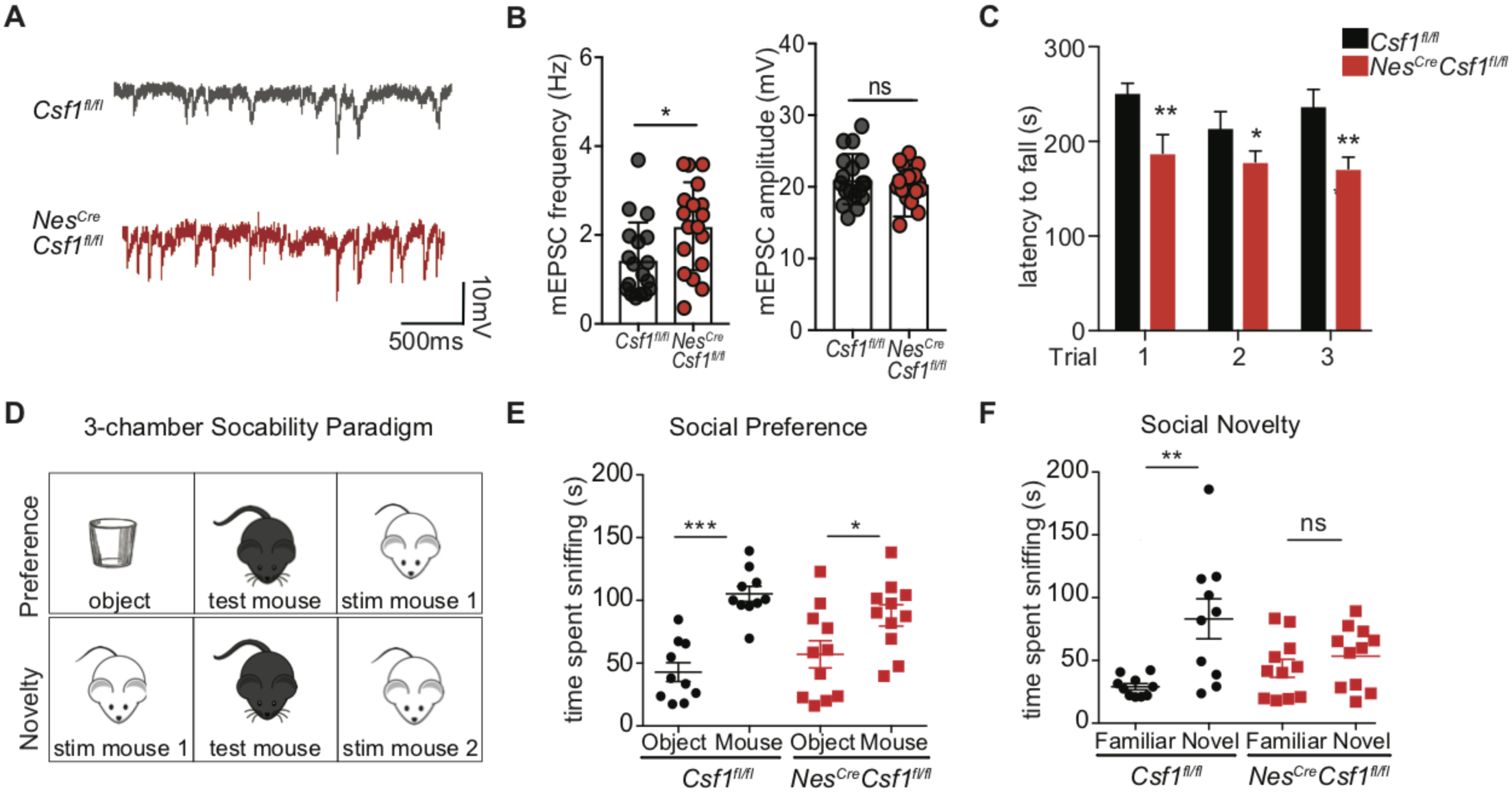
*Nes*^*Cre*^*Csf1*^*fl/fl*^ mice show electrophysiological Purkinje cell alterations and develop motor and behavioral defects. (A and B) Representative traces (A) and quantification (B) of PC mEPSC frequencies and amplitudes in *Nes*^*Cre*^*Csf1*^*fl/fl*^ PCs compared to controls. *n*=3 mice/genotype, *n*=6 cells/mouse. Data are a pool of three independent patch experiments. (C) Quantification of the latency for *Csf1*^*fl/fl*^ and *Nes*^*Cre*^*Csf1*^*fl/fl*^ mice to fall from accelerating rotating beam. *n*=12-15 mice/group, pooled from two independent experiments. (D to F) Schematic (D) and quantification (E and F) of the 3-chamber sociability paradigm used to assess defects in social preference (E) and social novelty (F) in *Csf1*^*fl/fl*^ and *Nes*^*Cre*^*Csf1*^*fl/fl*^ mice. *n*=10-12 mice/group, pooled from two independent experiments. (Graphs show means ± SEM. *\*P ≤ 0.05, **P ≤ 0.01, ***P ≤ 0.001, ****P≤0.0001* using multiple student’s t-tests (A), one-way ANOVA (C and D).

## Discussion

Cerebellar microglia display a unique morphological (Ashwell, 1990; Lawson et al., 1990) and transcriptional (Ayata et al., 2018; Grabert et al., 2016) profile, but little is known about their role in cerebellar development, homeostasis and function. Here, we explored the role of the macrophage growth factor CSF-1 for microglia development and maintenance in the cerebellum, and the consequences of nervous system CSF-1 depletion for cerebellar development and function. We show that despite both signaling through the same receptor, CSF-1R, the distinct expression patterns of IL-34 and CSF-1 shape microglia numbers and phenotypes in the forebrain and cerebellum. Genetic depletion of *Il34* or *Csf1* expression resulted in mutually exclusive reductions of postnatal microglia numbers, with IL-34 as the main forebrain cytokine, and CSF-1 as the main cerebellum cytokine. CSF-1 deficient animals, which lack cerebellar microglia, displayed alterations in cerebellar morphology, Purkinje cell loss with increased branching complexity, and motor learning and sociability defects.

Microglia form a grid-like pattern throughout the brain where each microglia controls its own territory that is not shared with others (Kettenmann et al., 2011), facilitating a fine-tuned microglia-neuron interaction. Strikingly, *Nes*^*Cre*^*Csf1*^*fl/fl*^ microglia were largely confined to central white matter regions of the cerebellum, instead of being evenly distributed throughout the brain, suggesting a migration defect during cerebellar development. The progressive radial distribution of microglia during cerebellar development and growth has been studied in the quail brain (Cuadros et al., 1997), but the mechanism of distribution of microglia throughout the brain during development is largely unknown. Cerebellar astrocytes express *Csf1* (Fig. S1 A), a known chemotactic factor for macrophages (Calvo et al., 1998; Pixley and Stanley, 2004). Because cerebellar astrocytes extend their long and ordered cellular processes throughout several cerebellar layers, they might thus serve as guiding structures for developing microglia. Our data, indeed, suggests that microglia may use CSF-1 secreting cells as a scaffold for radial migration into the gray matter, since the absence of neuroectodermal CSF-1 production left remaining microglia stuck in the central white matter of the cerebellum.

Intriguingly, functional pathways induced in postnatal forebrain and cerebellar microglia stimulated with either IL-34 or CSF-1 mirrored the altered biological pathways of microglia from IL-34 or CSF-1 deficient mice, revealing that CSF-1 and IL-34 are among the key drivers of regional microglial transcriptional signatures. Even more interestingly, microglia isolated from mice that underwent maternal immune activation (MIA) *in utero*, were also shown to upregulate gene networks involved in embryonic development (Mattei et al., 2017), similar to those described here by CSF-1 treatment, suggesting that increased CSF-1 expressed during MIA could potentially contribute to this transcriptional phenotype.

We also show that CSF-1 deficient animals, which lack cerebellar microglia, display alterations in cerebellar morphology, Purkinje cell number and function, and motor learning and sociability defects. Previous studies have shown that microglia serve a key role in proper cerebellar climbing fiber elimination, both shifting the excitatory drive and decreasing the inhibitory drive onto PCs, starting early in cerebellar development and lasting into adulthood (Nakayama et al., 2018). Indeed, *Nes*^*Cre*^*Csf1*^*fl/fl*^ cerebella have increased VGluT2 excitatory inputs onto developing (P7) PC soma compared to controls, suggesting a defect in early stage climbing fiber elimination (Ichikawa et al., 2016). Moreover, reduced pruning due to microglia depletion in CSF-1 deficient cerebella could be a potential mechanism by which adult *Nes*^*Cre*^*Csf1*^*fl/fl*^ PC branching is increased, and thereby also contributing to the increased firing potential (mEPSC) seen in their PCs. Not only can increased excitatory drive result in neuronal, and particularly PC death (Johnston, 2005; Slemmer et al., 2005), but altered climbing fiber elimination can also lead to cerebellar motor dysfunction and ataxia (Johnson et al., 2007). Lastly, increasing evidence suggests that cerebellar dysfunction may also contribute to a subset of ASD (Fatemi et al., 2012), a heterogeneous group of disorders, which presents a substantial challenge to diagnosis and treatment. Accordingly, post-mortem brain studies of patients with ASD have consistently reported reduced PC counts in the cerebellar cortex (Bauman and Kemper, 2005; Fatemi et al., 2012) and cerebellar injury is linked to increased incidence of ASD (Limperopoulos et al., 2007). Moreover, studies in mouse models have shown that genetic disruption of cerebellar PC function can lead to similar social interaction defects as shown in CSF-1 deficient cerebella (Tsai et al., 2012).

Previous reports suggested that PC express *Csf1r* (Murase and Hayashi, 2002; Nandi et al., 2012). However, we found no *Csf1r* or *CSF1R* expression in mouse and human PC, respectively, using multiple techniques and different ages, providing evidence for microglial deregulation as the single driver for the neuronal and behavioral perturbations observed in our CSF-1 deficiency model. In addition, *Iba1*^*Cre*^*Csf1r*^*fl/fl*^ mice showed similar PC perturbations (Nakayama et al., 2018), again supporting a microglial role for PC development and regulation. Further studies must be performed to exclude a transient, but relevant PC *Csf1r* expression during embryonic or early postnatal development.

The distinct distribution of IL-34 and CSF-1, and the distinct regional microglial transcriptional signatures described here, are conserved in both the mouse and human brain, highlighting the importance of this discrete expression and signaling for microglial biology. Altogether these results reveal the CSF-1-CSF-1R axis as a critical regulator of cerebellar development and integrity, and identify the CSF-1-CSF1-R axis as a potential therapeutic target to help restore cerebellar microglial and PC function, and to improve developmental motor and sociability disorders.

## Supplementary Figure Legends

**Figure S1.**
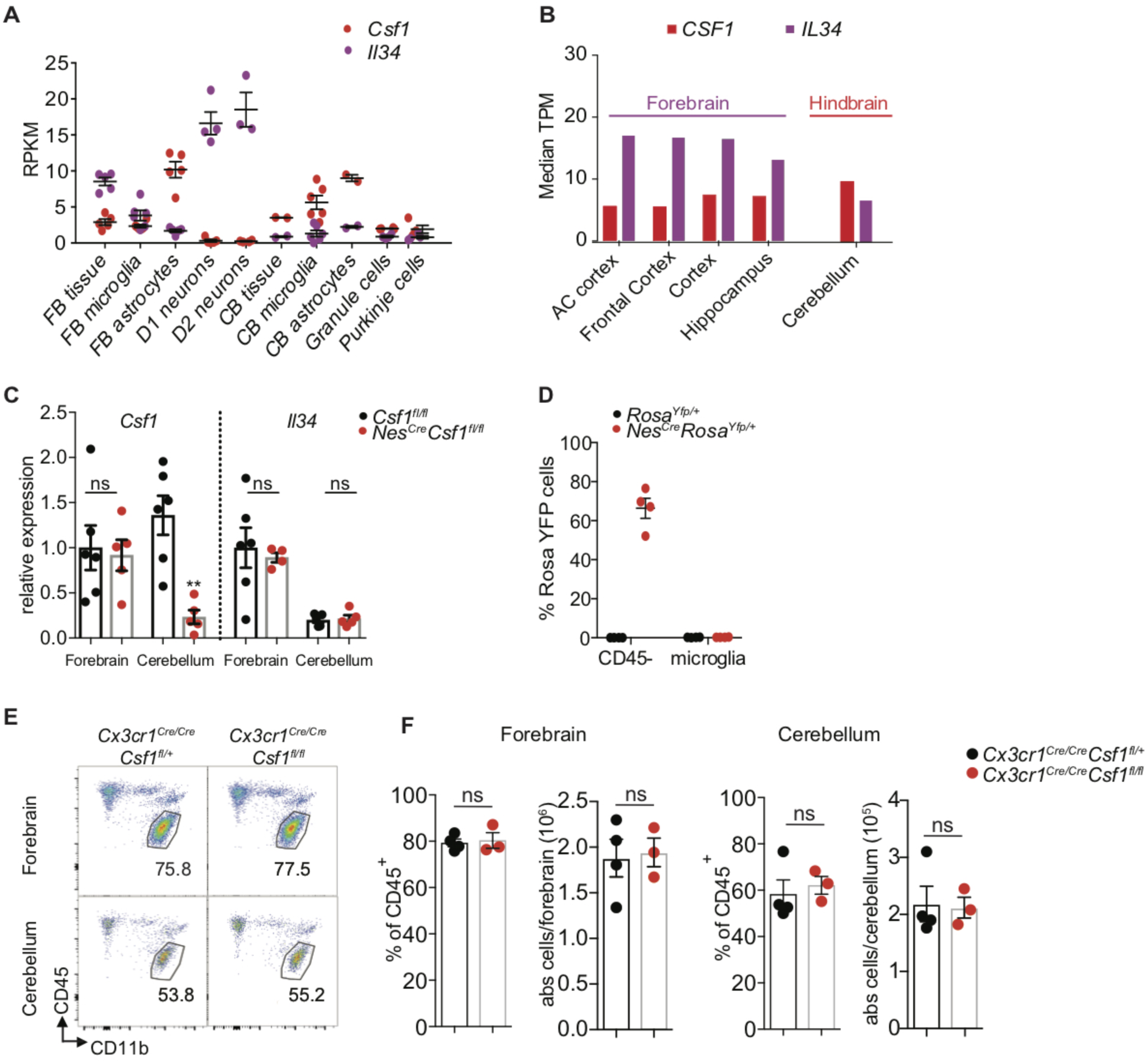
Distinct expression patterns of CSF-1 and IL-34 RNA in mice and human. (A) Normalized (RPKM) *Csf1 and Il34* mRNA expression levels in different brain tissues, glial cells, and neuronal cell populations from the forebrain and cerebellum. Expression data generated by TRAP sequencing. *n*=4-5/tissue or cell-type. (B) Normalized (median TPM) gene expression of *IL34* and *CSF1* mRNA transcripts from different human forebrain regions and cerebellum. Expression data generated by RNA seq and downloaded from the Human Protein Atlas database. (C) Relative qPCR expression of *Csf1* and *Il34* mRNA in forebrain and cerebellum from 6-7 week old *Csf1*^*fl/fl*^ and *Nes*^*Cre*^*Csf1*^*fl/fl*^ mice. All groups normalized to *Csf1*^*fl/fl*^ forebrain expression. *n*=3-4 mice/group. Data are a pool of two independent experiments. (D) Flow cytometric analysis showing *Yfp* (Nestin) expression in DAPI^−^CD45^−^CX3CR1^−^ parenchymal cells and DAPI^−^CD45^int^CX3CR1^hi^ microglia from adult *Rosa*^*Yfp/+*^ and *Nes*^*Cre*^*Rosa*^*Yfp/+*^ brains. *n*=3-4 mice/group. (E and F) Representative pseudocolor plots (E) and quantification (F) of flow cytometric analysis of forebrain and cerebellar doublet^−^DAPI^−^ CD11b^+^CD45^int^ microglia from *Cx3cr1*^*Cre*^*Csf1*^*fl/wt*^ and *Cx3cr1*^*Cre*^*Csf1*^*fl/fl*^ mice. Numbers adjacent to gate show percentages of CD45^+^ cells. *n*=3-4 mice/group. (Graphs shows mean ± SEM. *n.s.*=not significant).

**Figure S2.**
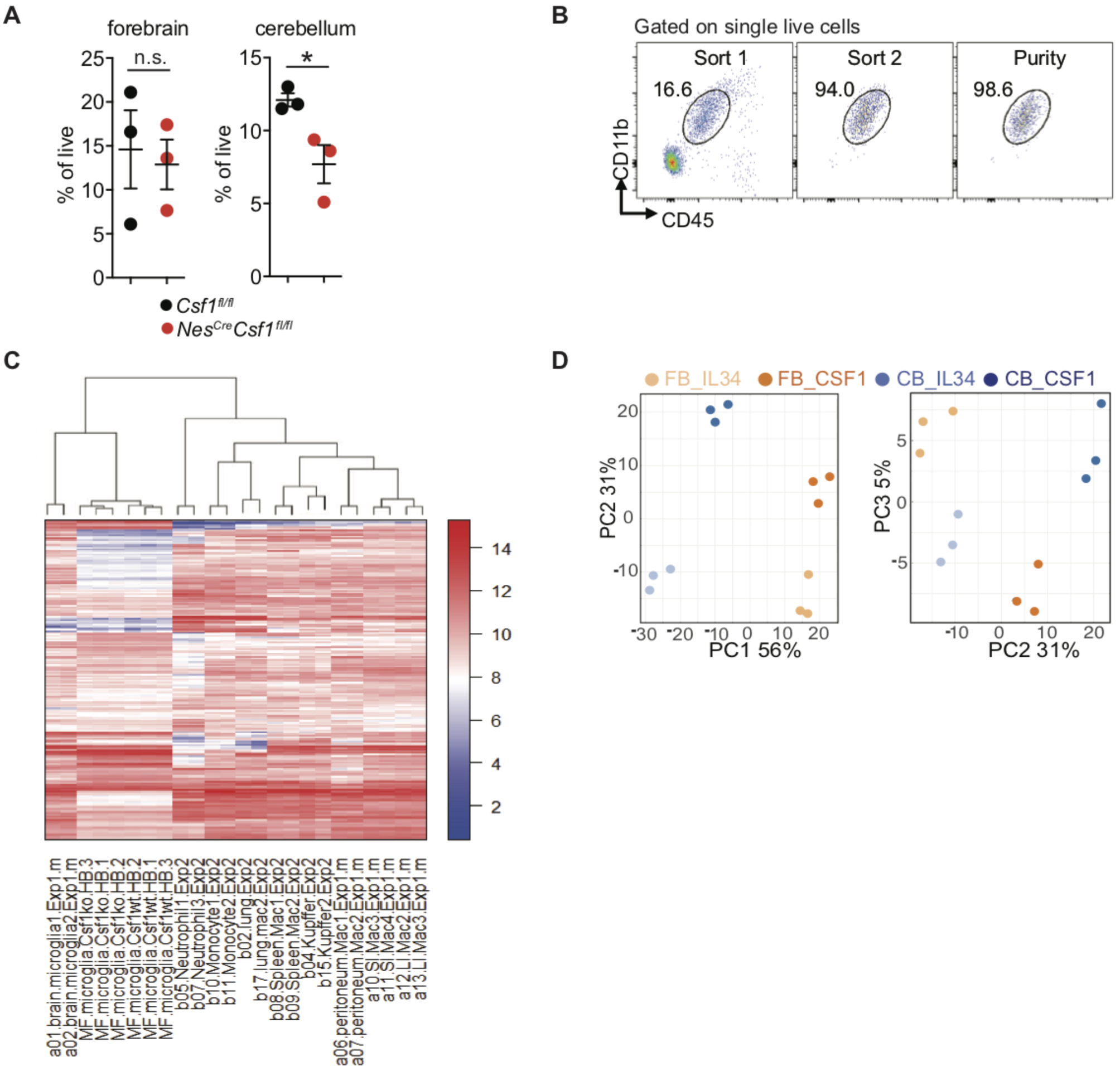
*Nes*^*Cre*^*Csf1*^*fl/fl*^ microglia profile clusters with published microglia profile and distinct clustering of CSF-1 or IL-34 stimulated forebrain and cerebellar neonatal microglia. (A) Flow cytometric quantification of doublet^−^DAPI^−^CD11b^+^CD45^int^ microglia from P8 forebrain (left) and cerebellum (right) of *Csf1*^*fl/fl*^ and *Nes*^*Cre*^*Csf1*^*fl/fl*^ littermates, used for NexGen sequencing. *n=3* mice/group. (B) Flow cytometry sorting strategy for microglia, used for NexGen sequencing. Cells were double sorted to reach purity >98%. *n=3* mice/group. (C) Heat map of hierarchically clustered transcriptomes of P8 *Csf1*^*fl/fl*^ and *Nes*^*Cre*^*Csf1*^*fl/fl*^ cerebellar microglia alongside previously published expression profiles of monocytes, tissue-resident macrophages, and neutrophils (Lavin et al., 2014). *Csf1*^*fl/fl*^ and *Nes*^*Cre*^*Csf1*^*fl/fl*^ microglia cluster with the other microglial transcriptomes. Scale represents row Z-score. (D) PCA of stimulated neonatal cortical and cerebellar microglia showing discrete clustering of samples. PC1 on the left plot shows 56% of variability in the expression data can be explained by brain region of microglia, while PC2 on the left and right plots shows 31% of variability can be explained by cytokine stimulation, regardless of brain region origin of the microglia.

**Figure S3.**
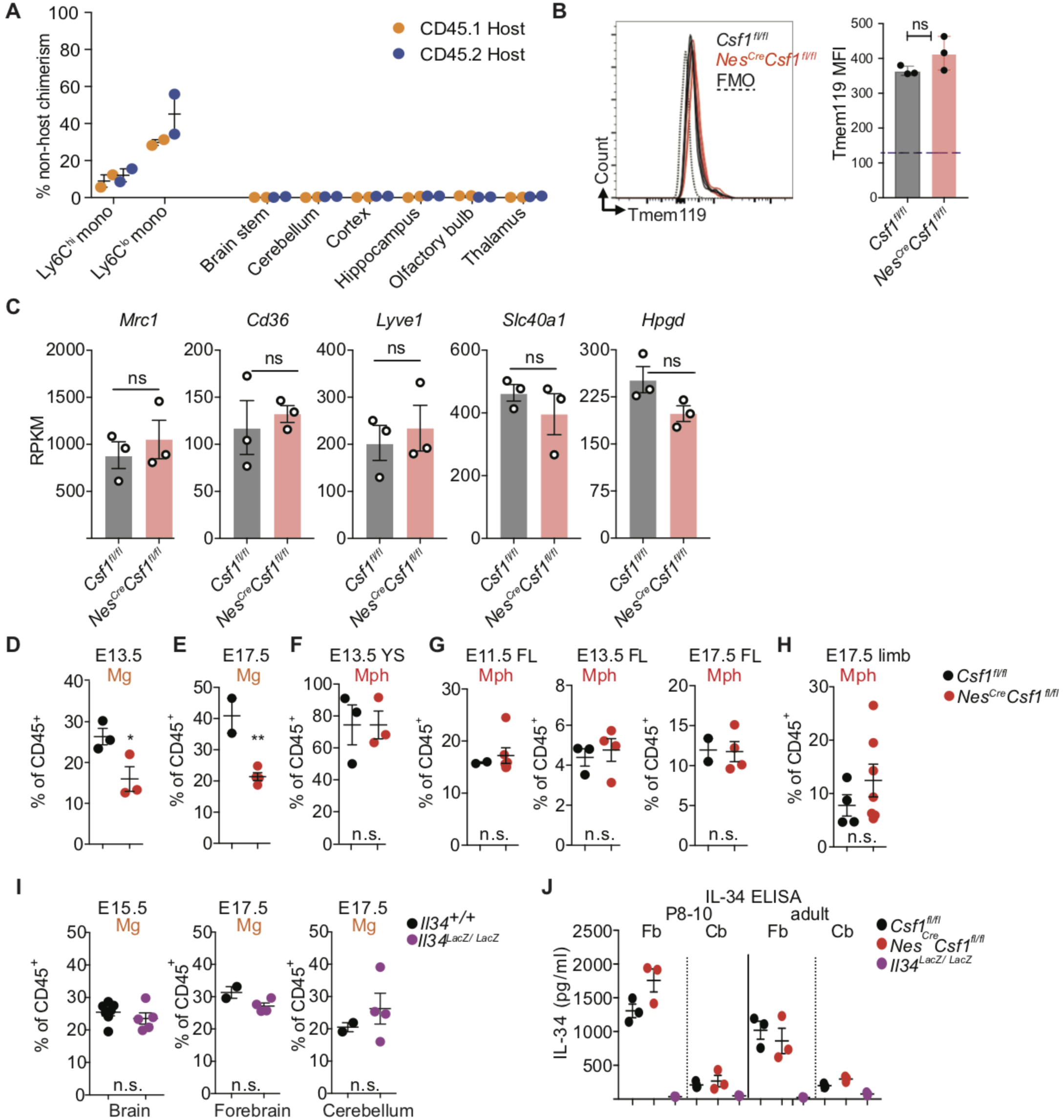
Characterization of cerebellar microglia and embryonic macrophages in control and CSF-1 deficient mice. (A) Percent chimerism of monocytes and total CD45^+^ cells in tissue of various brain regions from CD45.1 and CD45.2 parabionts anastomosed for 3 months. *n*=2 mice/group. (B) Flow cytometric analysis of Tmem119 expression in cerebellar microglia from control and CSF-1 deficient mice including fluorescence minus one (FMO) control. *n=*3 mice/genotype. Data are representative of two independent experiments. (C) Normalized (RPKM) mRNA expression of select perivascular macrophage genes in microglia from *Csf1*^*fl/fl*^ and *Nes*^*Cre*^*Csf1*^*fl/fl*^ P8 cerebella. *n=*3 mice/group. (D and E) Quantification of the percentage of microglia in whole brain rudiments from embryos at E13.5 (D) and E17.5 (E) in *Csf1*^*fl/fl*^ and *Nes*^*Cre*^*Csf1*^*fl/fl*^ mice. *n*=2-3 mice/group at each age. Data are representative from at least two independent experiments. (F to H) Quantification of the percentage of macrophages (Mph) in yolk sac (F), fetal liver (G), and limbs (H) of embryos at E13.5, E11.5, and E17.5 in *Csf1*^*fl/fl*^ and *Nes*^*Cre*^*Csf1*^*fl/fl*^ mice. *n*=2-7 mice/group at each age. Data are pooled from at least two independent experiments. (I) Quantification of the percentage of microglia (Mg) in whole brains from E15.5, and isolated forebrain and cerebellum from E17.5 *Il34*^*wt/wt*^ and *Il34*^*LacZ/LacZ*^ mice. *n*=2-7 mice/group. Data are pooled from at least two independent experiments. (J) ELISA for IL-34 protein (pg/ml) from P8-10 and adult forebrain and cerebellar brain tissue from *Csf1*^*fl/fl*^, *Nes*^*Cre*^*Csf1*^*fl/fl*^, and *Il34*^*LacZ/LacZ*^ mice. *n*=3 mice/group. (Graphs show mean ± SEM. *n.s.*=not significant, *\*\*P≤0.01* using student’s t-tests.)

**Figure S4.**
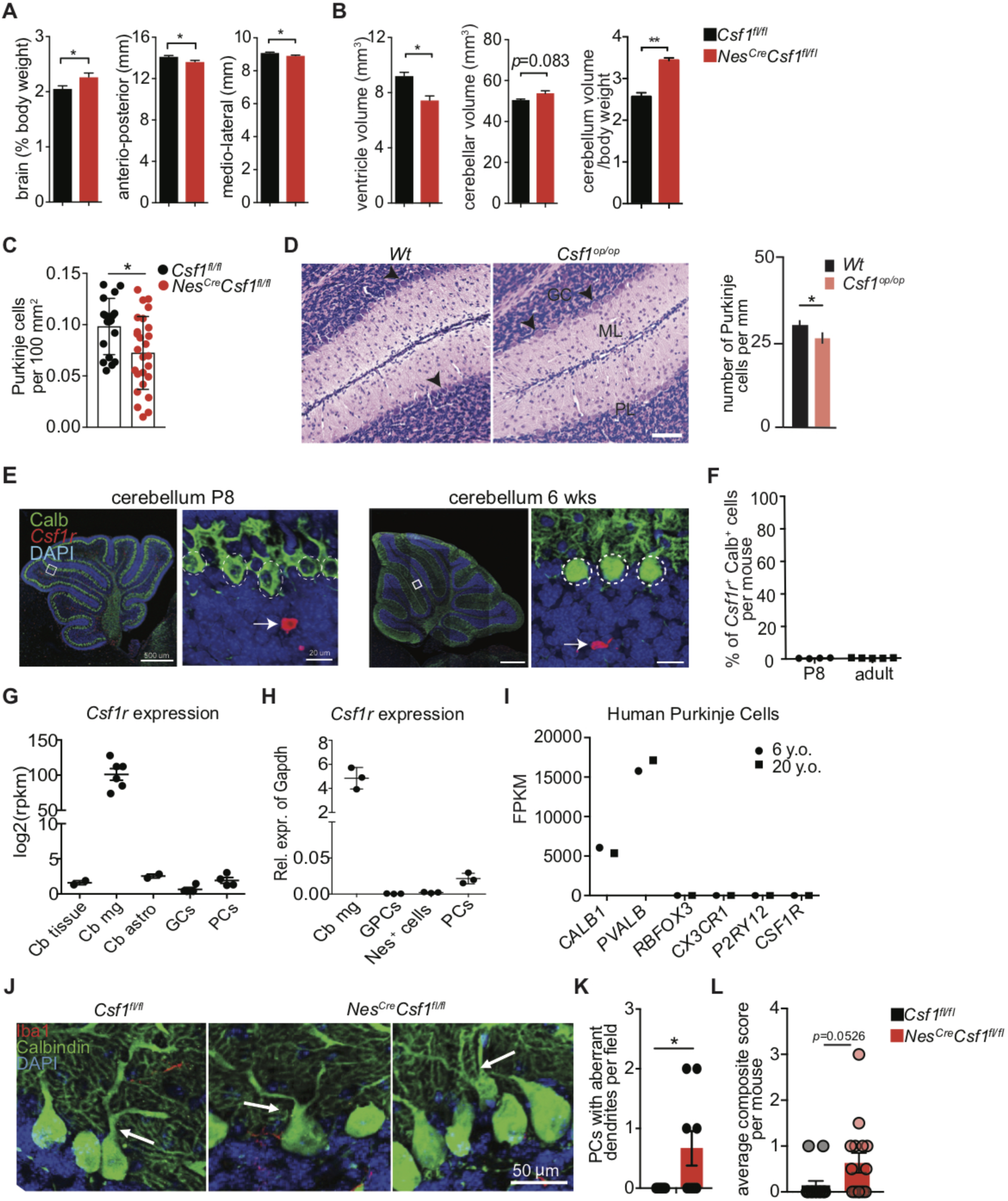
Morphological and behavioral alterations in CSF-1 deficient mice. (A and B) Quantification of brain mass and brain size (A), cerebellar ventricular volume, total cerebellar volume, and total cerebellar volume normalized to body weight (B) as measured by MRI in control and *Nes*^*Cre*^*Csf1*^*fl/fl*^ mice. *n*=3 mice/group, one experiment. (C) Quantification of Calbindin^+^ PC per mm^2^ of PC and molecular cell layers in P21 *Csf1*^*fl/fl*^ and *Nes*^*Cre*^*Csf1*^*fl/fl*^ cerebella. *n*=4 mice/group, 4-6 sections/mouse. (D) Representative H&E stainings (left) and quantification (right) of PCs from 2 week old wild type and *Csf1*^*op/op*^ cerebella. Arrows point to PCs. (GC): granule cell layer, (ML): molecular layer, (PL): PC layer. *n*=15 sections from 3 mice/genotype. Scale bar: 75 µm. Representative experiment from two independent experiments. (E and F) Representative immunofluorescence images (E) and quantification (F) of *Csf1r* mRNA expression in Calb^+^ PCs from P8 and 6 week old wild type cerebella. Scale bars: 500 µm (overview) and 20µm (inserts). *n=*4-5 mice/group. (G) Normalized (RPKM) *Csf1r* mRNA expression in different cell-types from adult wild type cerebella. Expression data generated by TRAP sequencing. (Cb): cerebellum, (mg): microglia, (astro): astrocyte, (GCs): granule cells. *n=*2-6 mice/cell type. (H) Relative expression of *Csf1r* in different cell-types from P6 wild type cerebella. Expression data generated by RT-PCR. (GPCs): granule precursor cells, (Nes^+^): Nestin^+^. *n=*3 mice/cell type. (I) Normalized expression (FPKM) of neuronal and microglial specific genes in laser capture micro-dissected PCs from cerebella of a 6 y.o. and 20 y.o. patient. Expression data generated by RNA seq and downloaded from ENCODE consortium. 6 y.o.: *n=*30 cells, 20 y.o.: *n=*20 cells. (J and K) Representative immunofluorescence stainings (J) and quantification (K) of Calb^+^ PCs with multiple dendrites emerging from their soma, in adult *Csf1*^*fl/fl*^ and *Nes*^*Cre*^*Csf1*^*fl/fl*^ cerebella. *n*=3 mice/group, *n=*7-11 high power fields/genotype. (L) Quantification of the percentage of mice positive for ataxia across multiple models (composite score of kyphosis presence, gait analysis, hanging wire test, ledge test, and hind limb clasping presence) in adult *Csf1*^*fl/fl*^ and *Nes*^*Cre*^*Csf1*^*fl/fl*^ mice. *n*=14-15 mice/group. (Graphs show means ± SEM. *\*P≤0.05,**P≤0.01* using student’s t-test.)

**Figure S5.**
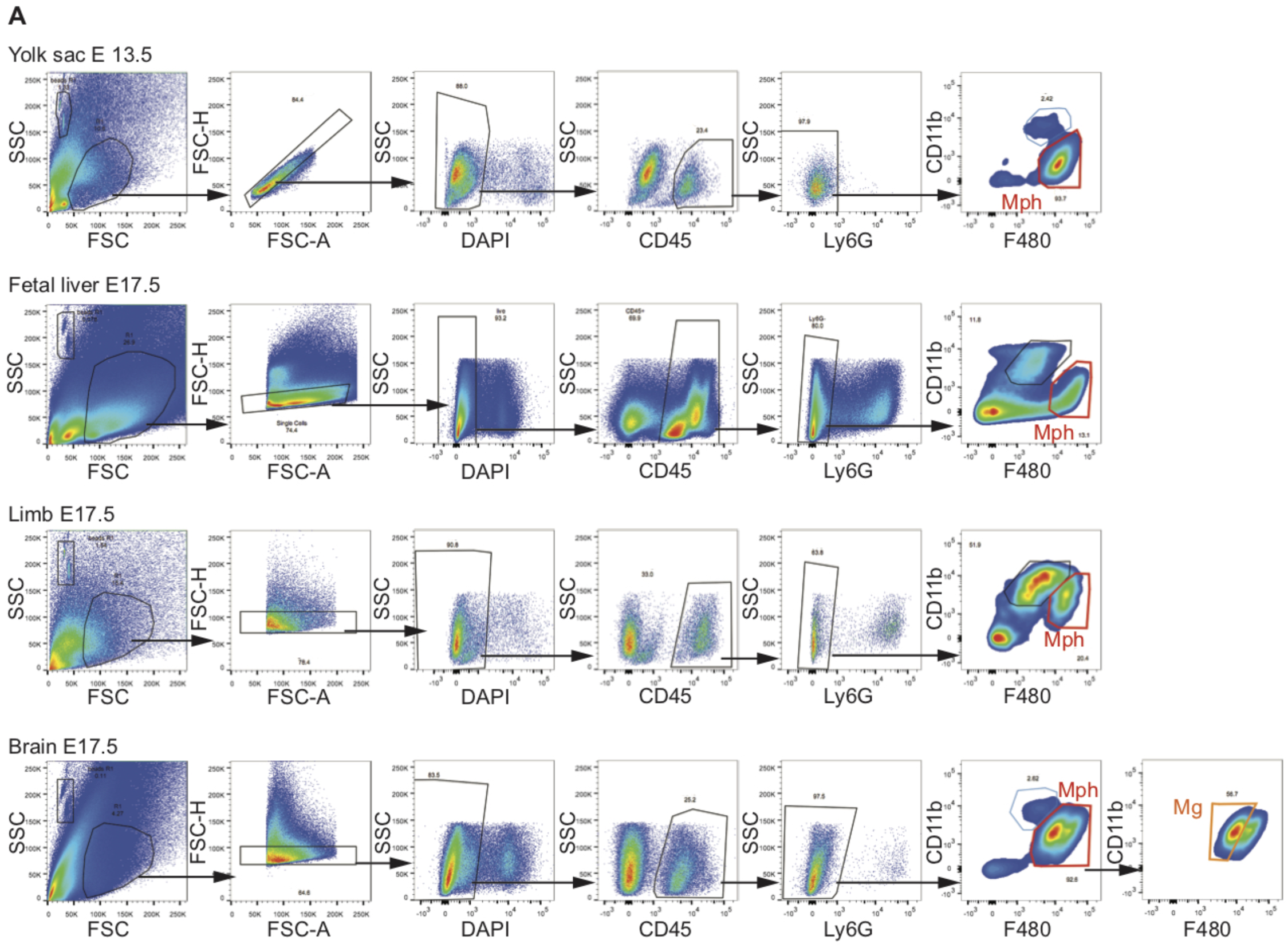
Gating strategies for flow cytometry analysis of embryonic tissues in CSF-1 deficient mice. (A) Representative gating strategies of yolk sac, fetal liver, limbs and brains (shown is forebrain) of *Csf1*^*fl/fl*^ and *Nes*^*Cre*^*Csf1*^*fl/fl*^ embryos at the specified ages of development. These gating strategies were used in Figure 3 A and Supplementary Figure 3 D to J. Mph: macrophages, Mg: microglia.

## Materials and Methods

### Animals

*Nes*^*Cre*^, *Cx3cr1*^*Cre*^, C57BL/6 CD45.2 and CD45.1, Ubiquitin-GFP, and R26-stop-EYFP were purchased from Jackson Laboratory. *Csf1*^*fl/fl*^ mice were generously provided by Sherry Abboud Werner (Harris et al., 2012). *Il34*^*Lacz/Lacz*^ mice were generated by EUCOMM (IKMC Project – ID: 33127) (generously provided by Marco Colonna) (Wang et al., 2012). *Nes*^*Cre*^*Csf1*^*fl/fl*^ and *Nes*^*Cre*^*Csf1*^*fl/fl*^*Il34*^*LacZ/LacZ*^ mice were generated in house by crossing *Nes*^*Cre*^, *Csf1*^fl/fl^, and *Il34*^*Lacz/Lacz*^ mouse lines. *Csf1*^*op/op*^ were originally obtained from Jackson Laboratory on an outbred C57/BL/J C3Heb/FeJ-a/a CD1 background and backcrossed for at least 10 generations onto the FVB/NJ background. *Tg(Aldh1l1-eGFPL10a)JD130, Tg(Pcp2-eGFPL10a)DR168, Tg(NeuroD1-eGFPL10a)JP241, Tg(Drd1-eGFPL10a)CP73*, and *Tg(Drd2-eGFPL10a)CP101* mice were used for astrocyte-, Purkinje cell-, granule cell-, D1 neuron-, and D2 neuron-TRAP, respectively (Doyle et al., 2008a; Doyle et al., 2008b; Heiman et al., 2008b). *Eef1a1*^*LSL.eGFPL10a/+*^ mice, which carry a loxP-flanked STOP cassette (LSL) upstream of the *eGFP-L10a* gene under the control of eukaryotic translation elongation factor 1 alpha 1, *Eef1a1*, (generously provided by Ana Domingos) (Stanley et al., 2013) were crossed with *Cx3cr1*^*CreErt2/+(Litt)*^ (generously provided by Dan Littman) (Parkhurst et al., 2013b) to generate a microglia-specific TRAP mouse. Microglia-specific mRNA enrichment was validated (Ayata et al., 2018). To activate Tamoxifen-inducible Cre (CreErt2 or Cre/Esr1*), mice were gavaged at 4-6 weeks of age with five doses of 100 mg/kg of Tamoxifen (T5648, Sigma, St. Louis, MO) in corn oil (C8267, Sigma) with a separation of at least 48 hours between doses. For timed embryo studies, male and female mice were bred overnight and vaginal plugs looked for the following morning, with this stage considered E0. Mice were housed at two to five animals per cage with a 12-hour light/dark cycle (lights on from 0700 to 1900 hours) at constant temperature (23°C) with ad libitum access to food and water. All animal protocols were approved by IACUC at Icahn School of Medicine at Mount Sinai and were performed in accordance with NIH guidelines.

### RNAscope *in situ* hybridization

Expression and localization of *Csf1r, Csf1 and Il34* was assessed using RNAscope *in situ* hybridization technology (ACD) following the manufacturer’s instructions. The multiplexed assay was performed on fixed frozen brain, cut in the sagittal plane at 14 micron thickness. Proprietary probes to *Csf1r, Il34*, and *Csf1* were hybridized to their respective target RNA for 2hrs at 40°C and their signal amplified for 15-30 minutes at 40 degrees. The last amplification step contained the fluorophores used for visualization, Alexa488 for *Il34*, Atto550 for *Csf1*, and Atto647 for *Csf1r*. Slides were then counter-stained with DAPI for 2 minutes, mounted with ProLong Gold (Invitrogen), and cover-slipped. Hybridized brain slices were visualized using confocal microscopy (Zeiss) and NanoZoomer whole slide imaging software (Hamamatsu).

### Single molecule RNA in situ hybridization with immunofluorescence

6 week old wild type mice (n=5) were anesthetized with ketamine (120 mg/kg) and xylazine (24 mg/kg) and perfused transcardially with 10 ml PBS followed by 40mL 10% formalin in PBS. The brains were then removed and fixed in 10% neutral-buffered formalin for another 48 hours. Fixed brains were embedded in paraffin, sectioned at 5 µm, mounted on Superfrost® Plus slides (Fisher Scientific, Hampton, NH), baked in a dry oven for 1 hour at 60°C and stored at room temperature until further processing. *In situ* hybridization was carried out using RNAScope custom-designed probes for *Csf1r* in combination with the RNAScope 2.0 Red kit following the manufacturer’s recommendation (Advanced Cell Diagnostics, Newark, CA). After completing *in situ* hybridization before colorimetric reaction, sections were rinsed with ddH_2_O, 0.1X PBS, 0.25X PBS, 0.5X PBS, and 1X PBS, blocked in 2% normal goat serum in PBS for 1 hour at room temperature and incubated with Calbindin D-28 antibody (dilution 1:5000, Swant) in 2% normal goat serum in PBS overnight at 4°C. Calbindin signal was amplified with anti-rabbit horseradish-peroxidase-conjugated secondary antibody and Alexa Fluor® 488 tyramide from Tyramide Signal Amplification Kit (Life Technologies) following manufacturer’s instructions and nuclei were stained with DAPI (0.2 mg/ml). Sections were then rinsed 1X PBS, 0.5X PBS, 0.25X PBS, 0.1X PBS, and ddH_2_O; subjected to colorimetric reaction; dried 15 min at 60°C, mounted using EcoMount (EM897L, Biocare Medical, Concord, CA) and dried overnight. Single-plane tile scans were taken under LSM780 Confocal Microscope (Zeiss) with 10× objective for quantification and representation. *Csf1r*^+^ Calbindin^+^ Purkinje cells were manually counted using Zen 2011 software and represented as dot plots using GraphPad Prism v5.01.

### Immunofluorescence staining

Mice were transcardially perfused with cold 1X PBS and their brains dissected and put into 4% PFA overnight. Following fixation, brains were placed into 30% sucrose for 2 nights, before embedding into OCT. 20-micron sections were cut from each brain using a cryostat. Sections were then thawed at room temperature, washed in 1X PBS, and incubated with blocking buffer (1X PBS, 0.5% Triton-X, 2% BSA, 5% goat serum) for 1hr at 4 degrees. The following primary antibodies were used overnight at 4 degrees: Iba1 (1:500, WAKO), Ki-67 (1:100, eBiosciences), CD68 (1:250, Serotec), NeuN (1:100, Abcam), Calbindin (1:500, CB-955, Abcam), VGluT2 (1:1000, AB2251-I, EMD Millipore). Secondary antibodies (all from eBioscience) conjugated to their respective fluorophores were incubated at 1:200 for 2hrs at room temperature. Slides were then counterstained with DAPI and mounted with Prolong Gold (Invitrogen) media. Images were acquired using widefield and confocal microscopy (Zeiss), and analyzed using Fiji (ImageJ) and Imaris (Bitplane). Manual quantifications were performed by an observer blinded to genotypes.

### Cell suspension preparation and microglia isolation

Perinatal and adult brains, yolk sac, embryonic limbs, and fetal livers were dissected from mice at specified ages, as previously described (Greter et al., 2012; Hoeffel et al., 2015; Lavin et al., 2014). Brains of P21 and older were perfused with ice cold PBS prior to tissue isolation, apart from brains used for microglia sorting and subsequent ultra low input (ULI) RNA sequencing, according to Immgen standardized protocol listed on immgen.org. For the perinatal brain, yolk sac, and limbs the tissue was enzymatically digested with 0.4 mg/ul collagenase IV (Sigma C5138) for 10-20 minutes at 37°C, followed by mechanical trituration with an 18G blunt tipped syringe and filtration through a 70 micron filter. Adult brain was digested with 0.4mg/ul collagenase IV for 30 minutes. Fetal liver tissue was only mechanically triturated to obtain a homogeneous cell suspension. Cell pellets from all adult brain were further processed by centrifugation in 5ml of 40% Percoll (Sigma-Aldrich), at 2300 rpm for 25min with no brake. Myelin was aspirated and the cell pellet was washed twice with FACS buffer (PBS w/o Ca^2+^ and Mg^2+^ supplemented with 2% heat inactivated FBS and 5mM EDTA) prior to staining. For parabionts, brain regions specified were dissected, minced, and digested in RPMI supplemented with 10% FCS, 0.2 mg/ml collagenase IV and 0.06 mg/ml DNaseI, for 1 hour at 37°C. Cells were pelleted at 1350rpm for 5 minutes, and resuspended with 7ml of 30% Percoll. The cell suspension was spun at 3000 rpm for 10 mins with no brake, washed twice with FACS buffer prior to staining.

### Flow cytometry and microglia sorting

Following single cell suspension preparation, cells were pelleted and subsequently stained with fluorophore-conjugated antibodies against CD11b (clone M1/70), CD45 (clone 30-F11), CD45.1 (clone A20), CD45.2 (clone 104), CX3CR1 (clone SA011F11, BioLegend), F4/80 (clone BM8, BioLegend), Ly6C (clone HK1.4), Ly6G (clone 1A8, BioLegend), Gr-1 (clone RB6-8C5), CD115 (clone AFS98), MHC II (clone M5/114.15.2), CD11c (clone N418), CD64 (clone X54-5/7.1, BioLegend), CD86 (clone GL1, BioLegend), Tmem119 (clone 106-6, Abcam), and either DAPI or Propium Iodide viability dyes (all from eBioscience if not indicated otherwise). Flow cytometry was performed using a Fortessa analyzer (BD) and FACS sorting was performed using a FACS Aria II (BD) or LSRII (BD). The gating strategies used for flow cytometry analysis of embryonic tissues is adapted from (Hoeffel et al., 2015) and shown in Supplementary Figure 5. Resident microglia were sorted as doublet^−^DAPI^−^CD11b^+^CD45^int^ (Fig. S2 B). Flow cytometry data analysis was performed using FlowJo (TreeStar) software. For ultra-low-input (ULI) RNA seq, microglia were double sorted to reach a purity of >98% and 1000 cells were sequenced.

### Parabiosis

Parabiotic mice were generated by surgically linking age and size matched congenic CD45.2 (C57BL/6) and CD45.1 (C57BL/6) mice as previously described (Ginhoux et al., 2009).

### Primary neonatal microglia culture

Preparation of primary neonatal microglia culture was adapted from (Bronstein et al., 2013). The cerebellum and cortex from 10-20 newborn pups (P2-3) were dissected and mechanically triturated with a 1000p pipette in ice-cold 1X HBSS. Following incubation for 15 minutes at 37°C in 0.25% Trypsin-/EDTA (Fisher), cells were spun down and washed twice in 1X HBSS. Cells were then re-suspended in complete medium (DMEM, 10% FBS, 100 U/ml penicillin 100 mg/ml streptomycin) and split evenly into 3 poly-D-lysine (Sigma) coated T-75 flasks. Half of the medium was changed 3 days later, and subsequently every other day. Between days 10-14 post plating into flasks, the media was removed from each flask and the cells were dissociated by incubating in Accutase (ThermoFisher) for 10 minutes at 37°C. Cells were then pelleted, washed, and stained on ice for 30 minutes before FACS sorting. Sorted DAPI^−^CD11b^+^CD45^int^ microglia from the mixed culture were seeded at density of 2 × 10^5^ cells onto poly-D-lysine coated 12-well plates, in complete medium containing either 100 ng/ml IL-34 or 20 ng/ml CSF-1 for 24 hours.

### Human samples

Brain tissue was provided by the Netherlands Brain Bank (NBB; www.brainbank.nl). Informed consent for research purposes was obtained for brain autopsy and the usage of tissue and clinical information. For this study, brain tissues were collected from 2 female donors with bipolar disorder (aged 45) and multisystem atrophy (aged 60) and processed according to standardized protocols for next generation sequencing of primary microglia.

### Next generation sequencing

#### Human microglia

Primary microglia (pMG) obtained from the Netherlands Brain Bank were isolated as described before (Melief et al., 2016) with minor modifications. In short, 2-10 grams of superior temporal gyrus (STG) and cerebellum were collected from 2 female donors within 8 hours after death, stored in Hibernate medium on ice and the isolation procedure was started within 2 to 24 hours after autopsy. A single cell suspension was generated by dissociating the tissue mechanically through a metal sieve, followed by an enzymatic digestion using collagenase Type I (3700 units/mL, Worthington) and DNase I (200 µg/mL, Roche). Microglia were further purified using a Percoll gradient and positive selection using CD11b-conjugated magnetic microbeads (Miltenyi Biotec GmbH), which resulted in a 99% pure microglia population as analyzed by flow cytometry. Microglia were lysed in RLT buffer and RNA was extracted according to the protocol provided by the RNeasy Mini Kit (Qiagen, USA). cDNA synthesis and library preparation was performed using the SMART-Seq v4 Ultra Low Input RNA Kit and Low Input Library Prep Kit v2 (Clontech), respectively. Sequencing was performed using the Illumina NextSeq-500 system. Transcript abundances were quantified with the Ensembl GRCh38 cDNA reference using Kallisto version 0.43.0. Transcript abundances were summarized to gene level using tximport. The expression matrix was filtered for only transcripts with greater than 5 RPKM in both replicates of at least one brain region, leaving 9,927 genes. Differential expression statistics between different brain regions were generated using DEseq2. Genes with a Benjamini-Hochberg adjusted p-value < 0.1 were called differentially expressed. GO term enrichment was performed using the topGO package in R using genes upregulated in either brain region as input.

#### Primary adult mouse microglia

1 × 10^3^ forebrain and cerebellar microglia (DAPI^−^CD11b^+^CD45^int^) were FACS sorted (FACS Aria II, BD) according to the Immgen standardized protocol for ULI RNA sequencing (Tan et al., 2016). RNA extraction, library preparation, and sequencing were performed at the Broad Technology Labs Boston using the NextSeq-500 system (Illumina). Differentially expressed genes for ULI RNAseq were determined as having at least 10 reads and a p-value < 0.05. Differential expression statistics were calculated with the edgeR package in R. Comparison of CSF-1 deficient microglia RNA seq data to published data of sorted myeloid populations from adult mice (*40*) were downloaded from NCBI (GEO accession GSE63340). The dataset was re-normalized with the dataset from the present study using the TMM normalization as implemented in the calcNormFactors() function in the edgeR package. Datasets were hierarchically clustered using spearman correlation distance of the top 1000 differentially expressed genes among sample types as determined using the ebayes() function in the Limma package. For GO term enrichment analysis, DEGs were inputted to the Panther Classification System (patherdb.org) using the statistical enrichment test on the “GO biological process complete” annotation data set. Gene lists for each GO term were generated in R using the intersection of the DEGs and Bioconductor’s Genome wide annotation for Mouse (org.Mm.egGO2ALLEGS). Panther Classification System and Bioconductor’s Genome wide annotation for Mouse are both sourced by the Gene Ontology database. Gene set enrichment analysis plots were generated using GSEA software (Subramanian et al., 2005) using our DEG values for P8 *Csf1*^*fl/fl*^ and *Nes*^*Cre*^*Csf1*^*fl/fl*^ cerebellar microglia and pathway gene set lists curated from our GO term analysis, with 1000 gene permutations. To compare human and mouse microglia, DEseq2 differential expression tables for human and mouse brain region comparisons were filtered for one to one orthologues (determined via Ensembl) that were expressed greater than 5 RPKM in all replicates of at least one brain region in both species, leaving 7,071 genes. To determine conserved brain region expression differences, genes that had an adjusted p-value < 0.1 and a log_2_FC > 0.5 in the same direction in both species were selected.

#### Cultured neonatal mouse microglia

Following 24hr incubation with either IL-34 or CSF-1, the supernatant was removed and the attached microglia were harvested directly in Qiazol (Qiagen). Total RNA was extracted using the miRNeasy Kit (Qiagen) following the manufacturer’s protocol. cDNA synthesis and library preparation was performed using the SMART-Seq v4 Ultra Low Input RNA Kit and Low Input Library Prep Kit v2 (Clontech), respectively. Sequencing was performed using the Illumina NextSeq-500 system. Transcript abundances were quantified with the Ensembl GRCm38 Mouse cDNA and ncRNA reference using Kallisto version 0.43.0. Transcript abundances were summarized to gene level using tximport. The expression matrix was filtered for only transcripts with greater than 2 TPM in all replicates of at least one condition, leaving 10,751 genes. Differentially expressed genes for neonatal microglial cultures were determined using a one-way ANOVA and a Benjamini-Hochberg adjusted p-value threshold < 0.01. Differential expression statistics between specific conditions were calculated using the Limma package in R (Ritchie et al., 2015). Hierarchical clustering of genes was done using Pearson correlation distance and average linkage. PCA for the neonatal cultures was done using Log2(TPM + 1) values for genes determined to be differentially expressed by one-way ANOVA, 675 genes in total. Genes specifically upregulated for either cytokine were identified by selecting those that had at least a log2FC > 0.25 and an adjusted p-value < 0.05 in one condition compared to all of the other 3.

### Translating ribosome affinity purification (TRAP)

6-8 week old transgenic *Tg(Aldh1l1-eGFPL10a)JD130, Tg(Pcp2-eGFPL10a)DR168, Tg(NeuroD1-eGFPL10a)JP241, Tg(Drd1-eGFPL10a)CP73, Tg(Drd2-eGFPL10a)CP101*, and *Cx3cr1*^*CreErt2/+(Litt)*^; *Eef1a1*^*LSL.eGFPL10a/+*^ mice (n=2-6 per genotype) were euthanized with CO_2_ and brain regions of interest were dissected. Ribosome-associated mRNA from neurons, microglia or astrocytes was isolated from each region as previously described (Heiman et al., 2014; Heiman et al., 2008a) where each sample corresponds to a single mouse. RNA clean-up from TRAP samples and 5% of the unbound fractions from TRAP samples was performed using RNeasy Mini Kit (Qiagen) following manufacturer’s instructions. RNA integrity was assayed using an RNA Pico chip on a Bioanalyzer 2100 (Agilent, Santa Clara, CA), and only samples with RIN>9 were considered for subsequent analysis. Double-stranded cDNA was generated from 1-5 ng of RNA using Nugen Ovation V2 kit (NuGEN, San Carlos, CA) following manufacturer’s instructions. Fragments of 200 bp were obtained by sonicating 500 ng of cDNA per sample using the Covaris-S2 system (Duty cycle: 10%, Intensity: 5.0, Bursts per second: 200, Duration: 120 seconds, Mode: Frequency sweeping, Power: 23W, Temperature: 5.5°C - 6°C, Covaris Inc., Woburn, MA). Subsequently, these fragments were used to produce libraries for sequencing by TruSeq DNA Sample kit (Ilumina, San Diego, CA, USA) following manufacturer’s instructions. The quality of the libraries was assessed by 2200 TapeStation (Agilent). Multiplexed libraries were directly loaded on NextSeq 500 (Ilumina) with High Output single read sequencing for 75 cycles. Raw sequencing data was processed by using Illumina bcl2fastq2 Conversion Software v2.17. Raw sequencing reads were mapped to the mouse genome (mm9) using the TopHat2 package (v2.1.0) (Kim and Salzberg, 2011). Reads were counted using HTSeq-count (v0.6.0) (Anders et al., 2014) against the Ensembl v67 annotation. The read alignment, read counting as well as quality assessment using metrics such as total mapping rate, mitochondrial and ribosomal mapping rates were done in parallel using an in-house workflow pipeline called SPEctRA (Purushothaman and Shen, 2016). Dow plots representing Reads Per Kilobase of transcript per Million mapped reads (RPKM) of indicated genes were made on GraphPad Prism v5.01 (https://www.graphpad.com).

### IL-34 ELISA

Brains were isolated, indicated regions dissected, flash frozen in liquid nitrogen and stored at −80°C until further use. Proteins were extracted by homogenizing the tissue in ice-cold RIPA buffer (Sigma) and lysates were centrifuged at 16,000 × g at 4°C. Supernatants containing 70 µg of total protein were used to determine IL-34 concentration using an IL-34 ELISA kit (BioLegend, San Diego, CA).

### Quantitative RT-PCR

RNA was extracted from FACS-isolated GFP^+^ cells from P6 *Atoh1-GFP* mice (Chen et al., 2002), FACS-isolated CFP^+^ cells from P6 *Nestin-CFP* mice (Mignone et al., 2004), FACS-isolated TdTomato^+^ cells from P6 *Pcp2*^*Cre/+*^;*R26*^*LSL-TdTomato/+*^ mice (Madisen et al., 2010; Zhang et al., 2004), and 6 week old *Csf1*^*fl/fl*^ and *Nes*^*Cre*^*Csf1*^*fl/fl*^ mice using a miRNeasy Micro Kit (Qiagen), according to the manufacturer’s protocol. cDNA was prepared using the iScript cDNA synthesis kit (Bio-Rad). qRT-PCR was performed using PowerUp Sybr Green Master Mix (Applied Biosystems). Fold-changes in expression were calculated using the ΔΔCt method. The *Gapdh* gene was used to normalize the results. The following primer pairs were used: *Csf1*: F 5’ CATCCAGGCAGAGACTGACA and R 5’ CTTCGTGATCCTCCTTCCAG (Harris et al., 2012); *Csf1r*: F 5’ TGTCATCGAGCCTAGTGGC3’ and R 5’ CGGGAGATTCAGGGTCCAAG3’; *Il34* F 5’-CTTTGGGAAACGAGAATTTGGAGA and R 5’-GCAATCCTGTAGTTGATGGGGAAG; *Gapdh*: F 5’ CCAAGGTGTCCGTCGTGGATCT3’ and R 5’ GTTGAAGTCGCAGGAGACAACC3’.

### Human Purkinje cell and brain tissue RNA seq data

Single cell RNA seq data from human Purkinje cells were extracted from the ENCODE database (encodeproject.org). 6 yr old patient ENCODE listing: ENCSR900JSG, GEO:GSE78474. 20 yr old patient ENCODE listing: ENCSR8882YA, GEO:GSE78473. *IL34* and *CSF1* RNA seq data from human forebrain and cerebellar tissue were extracted from the Human Protein Atlas database (www.proteinatlas.org) (*52*) using the GTEx RNA seq dataset (https://gtexportal.org/home/).

### Patch-clamp recording

Animals were decapitated under isofluorane anesthesia. Brains were quickly removed and transferred into ice-cold (0 – 4°C) artificial cerebrospinal fluid (ACSF) of the following composition (in mM): 210.3 sucrose, 11 glucose, 2.5 KCl, 1 NaH2PO4, 26.2 NaHCO3, 0.5 CaCl2, and 4 MgCl2. Acute sagittal slices (350 µm) were prepared from the cerebellum of 4 week old mutant and control littermates. Slices were allowed to recover for 40 min at room temperature in the same solution, but with reduced sucrose (105.2 mM) and addition of NaCl (109.5 mM). Following recovery, slices were maintained at room temperature in standard ACSF composed of the following (in mM): 119 NaCl, 2.5 KCl, 1 NaH2PO4, 26.2 NaHCO3, 11 glucose, 2 CaCl2, and 2 MgCl2. Whole-cell patch-clamp recordings (Edwards et al., 1989; Tsai et al., 2012) were taken using borosilicate glass electrodes (3-5 MΩ), from the soma of visually identified Purkinje cells located at the ganglionic layer of the cerebellum, which are clearly visible even under 10x objective. Whole-cell voltage-clamp recordings were obtained with the internal solution containing (in mM): 120 Cs-methanesulfonate, 10 HEPES, 0.5 EGTA, 8 NaCl, 4 Mg-ATP, 1 QX-314, 10 Na-phosphocreatine, and 0.4 Na-GTP. Miniature excitatory EPSCs were passively recorded in the presence of TTX (1 µM; Abcam) in the standard ACSF (as above). Miniature EPSCs were recorded at −60 mV. Data were low-pass filtered at 10 kHz and acquired at 10 kHz using Multiclamp 700B (Axon Instruments) and pClamp 10 (Molecular Devices). Purkinje cells for recording were selected at random. Series and membrane resistance was continuously monitored, and recordings were discarded when these measurements changed by > 20%. Recordings in which series resistance exceeded 25MΩ were rejected. Detection and analysis of EPSCs were performed using MiniAnalysis (Synaptosoft). Series resistance was monitored and canceled using a bridge circuit, and pipette capacitance was compensated. Voltage signals were low-pass filtered at 10 kHz.

### VGluT2 and Purkinje cell quantification and structural analysis

#### Purkinje cell quantification

Male P7, P21 and 6 week old *Csf1*^*fl/fl*^ and *Nes*^*Cre*^*Csf1*^*fl/fl*^ littermates, and 2 week old FVB wild type and *Csf1*^*op/op*^ mice on FVB background, were perfused transcardially with 4% PFA, pH 7.4 and brains incubated overnight in the same. For P7 and P21 *Csf1*^*fl/fl*^ and *Nes*^*Cre*^*Csf1*^*fl/fl*^ mice, brains were cut in half sagittally, sectioned at 15 µm, stained with anti-Calbindin and anti-VGluT2 antibody and DAPI. Purkinje cells were counted in the manually determined Purkinje cell and molecular cell layers using Imaris. VGluT2 puncta on manually outlined Purkinje cell soma, were automatically determined using Imaris Spot Function. For 6 week old *Csf1*^*fl/fl*^ and *Nes*^*Cre*^*Csf1*^*fl/fl*^ mice, brains were cut in half sagittally, and sectioned at 5 µm. Slides were stained with hematoxylin and eosin and scanned for digital imaging (Pannoramic 250 Flash III whole-slide scanner, 3DHISTECH). All Purkinje cells of all whole cerebellar sections were manually counted by an observer blinded to genotypes. For wild type and *Csf1*^*op/op*^ mice, 5 µm thick paraffin embedded brain sections were stained with H&E and pictures obtained by a Leica microscope. PCs were quantitated from hematoxylin and eosin stained sections from cerebellar lobes IX-X by manual counting.

#### Intracellular labeling of Purkinje cells

Randomly selected Purkinje cells that were being passively recorded were filled with 0.4% biocytin (Tocris) using patch-clamp recording pipettes, once the recordings were completed. Neurons in deeper portions of the Purkinje-cell layer were targeted and filled for 15 min, and then the pipette was slowly withdrawn so that the cell membrane could reseal. Slices (350-mm thick) were then fixed in 4% paraformaldehyde in 0.1M phosphate buffer for 24 h, rinsed thoroughly in phosphate buffer solution (PBS) and incubated for 90 min in a PBS solution containing 0.5% Triton-X, 10% normal goat serum and streptavidin Alexa Fluor 488 conjugate (1:500, Life Technologies). Slices were then rinsed in PBS, mounted on Superfrost Plus slides (VWR International), air-dried, and cover slipped in Vectashield mounting media (Vector Labs) for the subsequent imaging.

#### Dendritic spine imaging and characterization

Images were acquired using an Upright LSM 780 Confocal microscope (Carl Zeiss). A non-biased approach was taken to characterize dendritic spines on secondary dendrites within 200 mm of the soma. To assure sufficient sampling, five dendritic spine segments were taken from each neuron, and three neurons were assessed per animal (Dumitriu et al., 2011). Dendritic segments were imaged using a 100X lens (numerical aperture 1.4; Carl Zeiss) and a zoom of 4.0. Images were taken with a resolution of 300 × 100 pixels, dwell time was 1.58 um/s, and the line average was set to 8. Pixel size was 0.03 mm in the x–y plane and 0.05 mm in the z plane. To assure consistency of imaging across different confocal sessions and at different depths of the tissue, power was consistently set to 3.0%, and gain was adjusted within the range of 600-800 units to achieve consistent light intensity values within the same set of neurons. Images were de-convolved using a 3D resolution enhancement with AutoDeblur software (Media Cybernetics) (Christoffel et al., 2011; Rodriguez et al., 2008; Rodriguez et al., 2006) and then run through the spot detector in Imaris (Bitplane). Purkinje cell dendritic spine reconstruction and analysis was performed using Imaris FilamentTracer. Start and end points of dendrites to be segmented were manually selected and traced using the AutoPath method, and automatic thresholds used to determine dendrite diameter and surface rendering. Spines were considered to have a minimum seed point diameter of 0.2 mm and a maximum spine length of 2.0 mm (including branched spines). The seed point threshold was manually decreased to detect the maximum seed points for each dendrite. Automatic threshold were used for spine diameter rendering. The average values for dendritic spine length, density, diameter, and volume measurements were calculated by the Imaris software and exported from the Statistics tab. All analyses were performed blinded to the group conditions until the analyses were completed.

#### Whole cell imaging and dendrite complexity quantification

Images were acquired using an Upright LSM 780 Confocal microscope (Carl Zeiss) using a 63X lens (numerical aperture 1.4; Carl Zeiss) and a zoom of 0.6. Images were taken with a resolution of 1024 x1024 pixels, dwell time was 1.58 um/s, and the line average was set to 4. Pixel size was 0.22um in the x–y plane and 1.00mm in the z plane. Gain was set to 600-800 units. Dendrite complexity quantification was performed using the Sholl analysis plugin in Fiji (ImageJ).

### Behavioral tests

#### Ataxia assessment

Cerebellar ataxia in *Csf1*^*fl/fl*^ and *Nes*^*Cre*^*Csf1*^*fl/fl*^ mice was assessed using a composite score of hind limb clasping, ledge test, gait and kyphosis, as previously described (Ginhoux et al., 2009).

#### Rotarod

Motor effects in *Il34*^*LacZ/LacZ*^ and *Nes*^*Cre*^*Csf1*^*fl/fl*^ animals, and their respective littermate controls, were tested using the accelerating rotarod paradigm (Brooks and Dunnett, 2009), with additional video recording. Mice were placed on a motorized beam that was accelerated from 4-40 rpm. The latency of the mice to fall was recorded. If an animal clung to the rod and completed a full passive rotation, the timer was stopped for the animal and the time was noted. Prior to the final recorded rotarod session on day 4, with 3 trials at 5 min intervals each, animals were habituated to the paradigm during 3 training days with 3 trials of 5 min each. Differences between groups were statistically evaluated by Student’s t tests.

#### Sociability assessments

Sociability deficits were assessed in 6-8 week old male *Nes*^Cre^*Csf1*^fl/fl^ mice and their littermate controls. The premise for these behavioral tests lies in (1) the preference of mice spending time around other mice over inanimate objects, and (2) with novel over familiar subjects (Ellegood and Crawley, 2015; File and Hyde, 1978). To begin, test mice were habituated to an empty 3-chambered Plexiglass box for 10min. Mice were then restricted to the middle chamber by dividers, while a novel inanimate object and a novel stimulus mouse were placed under wire cups in either of the 2 flanking chambers (social preference). The dividers were then removed and the test mouse was allowed to freely explore the entire box for 10min. Following this, test mice were again restricted to the middle chamber, while the inanimate object was replaced with a second novel stimulus mouse (social novelty). Test mice were again released from the middle chamber and allowed to explore for 10min the chambers with the familiar and new stimulus mice. The behavior sessions were recorded using EthoVision (Noldus), and the time each test mouse spent near and/or sniffing the object or stimulus mice was manually recorded. Stimulus mice were age and sex matched for these assessments.

### Magnetic Resonance Imaging

All imaging were performed at the Translational and Molecular Imaging Institute at Mount Sinai Hospital on a Bruker Biospec 70/30 7Tesla scanner with a B-GA12S gradient insert (gradient strength 440 mT/m and slew rate 3444 T/m/s). A Bruker four-Channel mice brain phased array was used for all data acquisition in conjunction with a Bruker volume transmit 86mm coil. All mice were imaged under isoflurane anesthesia (3% induction and 1.5% maintenance). All mice were imaged on a heated bed and respiration was monitored continuously. The following protocols were used. After a three-plane localizer. A DTI protocol was acquired with a Pulsed Gradient Spin Echo - EPI sequence with the following parameters: TR=5500ms, TE=22.672ms, 4 segments, 30 gradient directions with b-value = 1000 s/mm2 and 5 B0’s, FOV=25mm, Matrix=128×128, slice thickness=0.5mm, skip=0, 3 averages, total acquisition time=38mins. A T2 anatomical scan was obtained with a 3D-RARE (Rapid Acquisition with Relaxation Enhancement) sequence with a RARE factor of 8, TR=2500ms, TEeff=62.5ms, FOV=25.6mmx25.6mm, Slice Thickness=0.4mm, skip=0, 18 slices, matrix size 256×256, 30 averages, total acquisition time=31mins. Scans were transferred to offline workstation for processing. In house software TMII-ROI developed under Matlab (R2015b, The Mathworks Inc., Natick, MA 2000) was used to analyze volume.

### Statistical analysis

For all graphical analyses, mean values, S.E.M values, Student’s t-test (two-tailed, unpaired), and one-way ANOVA were calculated using Prism 7 (GraphPad).

## Supplemental Materials

*Supplementary Figure 1:* Distinct expression patterns of CSF-1 and IL-34 mRNA in mice and human.

*Supplementary Figure 2: Nes*^*Cre*^*Csf1*^*fl/fl*^ microglia profile clusters with published microglia profile and distinct clustering of CSF-1 or IL-34 stimulated forebrain and cerebellar neonatal microglia.

*Supplementary Figure 3:* Characterization of cerebellar microglia and embryonic macrophages in control and CSF-1 deficient mice.

*Supplementary Figure 4:* Morphological and behavioral alterations in CSF-1 deficient mice.

*Supplementary Figure 5:* Gating strategies for flow cytometry analysis of embryonic tissues in CSF-1 deficient mice.

*Table S1.* Gene ontology term pathways for RNA seq expression data of purified human cerebellar and forebrain microglia.

*Table S2.* Gene ontology term pathways for RNA seq expression data of sorted P8 and adult *Csf1*^*fl/fl*^ and *Nes*^*Cre*^*Csf1*^*fl/fl*^, and adult *Il34*^*wt/wt*^ and *Il34*^*LacZ/LacZ*^ cerebellar and forebrain microglia.

*Table S3.* Gene ontology term pathways for RNA seq expression data of cultured and sorted primary neonatal cerebellar and forebrain microglia stimulated with CSF-1 or IL-34.

All tables are provided in Excel format.

## Acknowledgments

We thank the members of the M.M. and B.D.B. laboratory and Soyon Hong for extensive discussions and critiques of the manuscript, the Immgen Consortium for sequencing, the ENCODE Consortium and the ENCODE production laboratories for generating the particular datasets, Patrick Hof for advice on Purkinje cell quantification, Sayan Nandi for help with Purkinje cell quantification and the Mount Sinai Flow Cytometry Core and Translational Imaging Core.

## Author contributions

V.K., F.A.D, M.C.-A., P.A., A.B., E.N., K.Y, M.S, I.-L.T., M.E.F., H.L.B., E.S., N.T., A.W., Y.L., P.S., and A.B. performed experiments. V.K., F.A.D., P.A., A.B., E.N., K.Y, I.-L.T., S.A.R., C.C, A.L., H.L.B., and P.S. analyzed data. V.K., F.A.D., P.A., A.B., E.N., K.Y., I.-L.T., P.S., F.G., S.N., V. C., H.M., A.S., and M.M. designed experiments. V.K., F.A.D., M.C.-A., P.A., A.B., E.N., K.Y., P.S., F.G., A.N., V.C., E.R.S., S.J.R., Z.Y., B.D.B., A.L.J., L.D.W., H.M., A.S. and M.M. provided intellectual contribution to the project. V.K., F.A.D. and M.M. wrote the paper;

## Data and materials availability

All data is available in the main text or the supplementary materials. RNA Seq and ULI RNA Seq data will be deposited in GEO; accession number pending.

