## Supplemental Table 1-3 for "Disruption of the CSF-1-CSF-1R axis alters cerebellar microglia and is associated with motor and social interaction defects"

### Supplementary Table S1

| GO Term_ID | GO Term Pathway: Human_Cerebellum | Fisher pvalue | GO Term_ID | GO Term Pathway: Human_Superior Temporal Gyrus | Fisher pvlaue |
|---|---|---|---|---|---|
| GO:0006614 | SRP-dependent cotranslational protein ta... | < 1e-30 | GO:0007165 | signal transduction | 3.70E-05 |
| GO:0000184 | nuclear-transcribed mRNA catabolic proce... | < 1e-30 | GO:0007264 | small GTPase mediated signal transductio... | 4.10E-05 |
| GO:0019083 | viral transcription | < 1e-30 | GO:0023052 | signaling | 5.50E-05 |
| GO:0006413 | translational initiation | < 1e-30 | GO:0007154 | cell communication | 8.90E-05 |
| GO:0006364 | rRNA processing | < 1e-30 | GO:0071495 | cellular response to endogenous stimulus | 0.00015 |
| GO:0000956 | nuclear-transcribed mRNA catabolic proce... | < 1e-30 | GO:0035556 | intracellular signal transduction | 0.00021 |
| GO:0006402 | mRNA catabolic process | < 1e-30 | GO:1901701 | cellular response to oxygen-containing c... | 0.00035 |
| GO:0090150 | establishment of protein localization to... | < 1e-30 | GO:0001568 | blood vessel development | 0.00036 |
| GO:0006612 | protein targeting to membrane | < 1e-30 | GO:1901699 | cellular response to nitrogen compound | 0.00037 |
| GO:0019080 | viral gene expression | < 1e-30 | GO:0051056 | regulation of small GTPase mediated sign... | 0.0004 |
| GO:0006613 | cotranslational protein targeting to mem... | < 1e-30 | GO:0051091 | positive regulation of DNA binding trans... | 0.00041 |
| GO:0045047 | protein targeting to ER | < 1e-30 | GO:0007186 | G-protein coupled receptor signaling pat... | 0.0006 |
| GO:0072599 | establishment of protein localization to... | < 1e-30 | GO:0071417 | cellular response to organonitrogen comp... | 0.00064 |
| GO:0022613 | ribonucleoprotein complex biogenesis | < 1e-30 | GO:0032495 | response to muramyl dipeptide | 0.00065 |
| GO:0016072 | rRNA metabolic process | < 1e-30 | GO:0045943 | positive regulation of transcription fro... | 0.00065 |
| GO:0070972 | protein localization to endoplasmic reti... | < 1e-30 | GO:0051345 | positive regulation of hydrolase activit... | 0.00104 |
| GO:0042254 | ribosome biogenesis | < 1e-30 | GO:0001944 | vasculature development | 0.00112 |
| GO:0006412 | translation | 8.20E-30 | GO:0071407 | cellular response to organic cyclic comp... | 0.0013 |
| GO:0072657 | protein localization to membrane | 2.40E-29 | GO:0006356 | regulation of transcription from RNA pol... | 0.00134 |
| GO:0043043 | peptide biosynthetic process | 3.60E-29 | GO:0048514 | blood vessel morphogenesis | 0.00148 |
| GO:0006401 | RNA catabolic process | 5.50E-29 | GO:1901700 | response to oxygen-containing compound | 0.00159 |
| GO:0006605 | protein targeting | 2.20E-28 | GO:0022412 | cellular process involved in reproductio... | 0.00161 |
| GO:0034470 | ncRNA processing | 3.80E-27 | GO:1902531 | regulation of intracellular signal trans... | 0.00161 |
| GO:0043604 | amide biosynthetic process | 3.40E-26 | GO:0072358 | cardiovascular system development | 0.00173 |
| GO:0019439 | aromatic compound catabolic process | 5.10E-26 | GO:1902895 | positive regulation of pri-miRNA transcr... | 0.00178 |
| GO:0006518 | peptide metabolic process | 1.90E-25 | GO:0009719 | response to endogenous stimulus | 0.00202 |
| GO:1901361 | organic cyclic compound catabolic proces... | 4.60E-25 | GO:0051090 | regulation of DNA binding transcription ... | 0.00203 |
| GO:0034655 | nucleobase-containing compound catabolic... | 4.80E-25 | GO:0050896 | response to stimulus | 0.00206 |
| GO:0046700 | heterocycle catabolic process | 7.00E-25 | GO:0042089 | cytokine biosynthetic process | 0.00232 |
| GO:0044270 | cellular nitrogen compound catabolic pro... | 1.40E-24 | GO:0030522 | intracellular receptor signaling pathway | 0.00247 |
| GO:0034660 | ncRNA metabolic process | 5.40E-24 | GO:0051261 | protein depolymerization | 0.0028 |
| GO:0043603 | cellular amide metabolic process | 4.40E-21 | GO:0051716 | cellular response to stimulus | 0.00291 |
| GO:0016071 | mRNA metabolic process | 6.80E-21 | GO:0008015 | blood circulation | 0.00309 |
| GO:0016032 | viral process | 1.40E-18 | GO:1901652 | response to peptide | 0.00321 |
| GO:0044403 | symbiosis, encompassing mutualism throug... | 4.70E-18 | GO:0046058 | cAMP metabolic process | 0.00352 |
| GO:0044419 | interspecies interaction between organis... | 8.20E-18 | GO:0042107 | cytokine metabolic process | 0.00352 |
| GO:0072594 | establishment of protein localization to... | 1.30E-17 | GO:0003013 | circulatory system process | 0.0037 |
| GO:0006396 | RNA processing | 5.30E-17 | GO:0001659 | temperature homeostasis | 0.0038 |
| GO:1901566 | organonitrogen compound biosynthetic pro... | 1.20E-15 | GO:0070266 | necroptotic process | 0.0038 |
| GO:0044085 | cellular component biogenesis | 3.90E-14 | GO:0043551 | regulation of phosphatidylinositol 3-kin... | 0.0038 |
| GO:0033365 | protein localization to organelle | 4.60E-14 | GO:0097201 | negative regulation of transcription fro... | 0.0038 |
| GO:0006119 | oxidative phosphorylation | 1.20E-13 | GO:0061614 | pri-miRNA transcription from RNA polymer... | 0.0038 |
| GO:0044265 | cellular macromolecule catabolic process | 6.70E-13 | GO:1902893 | regulation of pri-miRNA transcription fr... | 0.0038 |
| GO:0042773 | ATP synthesis coupled electron transport | 7.20E-13 | GO:0051336 | regulation of hydrolase activity | 0.00386 |
| GO:0042775 | mitochondrial ATP synthesis coupled elec... | 2.60E-12 | GO:0001578 | microtubule bundle formation | 0.00407 |
| GO:0002181 | cytoplasmic translation | 7.20E-12 | GO:0043547 | positive regulation of GTPase activity | 0.00429 |
| GO:0022904 | respiratory electron transport chain | 1.80E-11 | GO:0014070 | response to organic cyclic compound | 0.00493 |
| GO:0006886 | intracellular protein transport | 6.10E-11 | GO:2001237 | negative regulation of extrinsic apoptot... | 0.00512 |
| GO:0042255 | ribosome assembly | 6.80E-11 | GO:0007010 | cytoskeleton organization | 0.00521 |
| GO:0051704 | multi-organism process | 1.20E-10 | GO:0043122 | regulation of I-kappaB kinase/NF-kappaB ... | 0.0053 |
| GO:0046907 | intracellular transport | 1.60E-10 | GO:0008625 | extrinsic apoptotic signaling pathway vi... | 0.00533 |
| GO:0042273 | ribosomal large subunit biogenesis | 2.20E-10 | GO:1901653 | cellular response to peptide | 0.00556 |
| GO:0046034 | ATP metabolic process | 2.20E-10 | GO:2001236 | regulation of extrinsic apoptotic signal... | 0.00582 |
| GO:0009057 | macromolecule catabolic process | 2.60E-10 | GO:0051254 | positive regulation of RNA metabolic pro... | 0.00583 |
| GO:0045333 | cellular respiration | 3.30E-10 | GO:0001525 | angiogenesis | 0.00588 |
| GO:0009205 | purine ribonucleoside triphosphate metab... | 3.60E-10 | GO:0044093 | positive regulation of molecular functio... | 0.00596 |
| GO:0071840 | cellular component organization or bioge... | 4.20E-10 | GO:0009966 | regulation of signal transduction | 0.00617 |
| GO:0009167 | purine ribonucleoside monophosphate meta... | 4.70E-10 | GO:0007178 | transmembrane receptor protein serine/th... | 0.00622 |
| GO:0009126 | purine nucleoside monophosphate metaboli... | 4.70E-10 | GO:1901342 | regulation of vasculature development | 0.00622 |
| GO:0009144 | purine nucleoside triphosphate metabolic... | 4.70E-10 | GO:0010522 | regulation of calcium ion transport into... | 0.00636 |
| GO:0009199 | ribonucleoside triphosphate metabolic pr... | 6.30E-10 | GO:0097191 | extrinsic apoptotic signaling pathway | 0.00636 |
| GO:0051649 | establishment of localization in cell | 6.90E-10 | GO:0032870 | cellular response to hormone stimulus | 0.00689 |
| GO:0009141 | nucleoside triphosphate metabolic proces... | 7.60E-10 | GO:1901698 | response to nitrogen compound | 0.00692 |
| GO:0009161 | ribonucleoside monophosphate metabolic p... | 1.40E-09 | GO:0071800 | podosome assembly | 0.00696 |
| GO:0022618 | ribonucleoprotein complex assembly | 2.40E-09 | GO:0002753 | cytoplasmic pattern recognition receptor... | 0.00696 |
| GO:0009123 | nucleoside monophosphate metabolic proce... | 2.80E-09 | GO:0097300 | programmed necrotic cell death | 0.00696 |
| GO:0006163 | purine nucleotide metabolic process | 3.20E-09 | GO:0010646 | regulation of cell communication | 0.00714 |
| GO:0009150 | purine ribonucleotide metabolic process | 3.60E-09 | GO:0007265 | Ras protein signal transduction | 0.00718 |
| GO:0072521 | purine-containing compound metabolic pro... | 4.60E-09 | GO:0009187 | cyclic nucleotide metabolic process | 0.0072 |
| GO:0022900 | electron transport chain | 5.10E-09 | GO:0051092 | positive regulation of NF-kappaB transcr... | 0.00725 |

| | | | | | |
|---|---|---|---|---|---|
| GO:0042274 | ribosomal small subunit biogenesis | 8.90E-09 | GO:0043087 | regulation of GTPase activity | 0.00738 |
| GO:0071826 | ribonucleoprotein complex subunit organi... | 9.90E-09 | GO:0071363 | cellular response to growth factor stimu... | 0.00743 |
| GO:0009259 | ribonucleotide metabolic process | 1.10E-08 | GO:0043123 | positive regulation of I-kappaB kinase/N... | 0.00757 |
| GO:0010629 | negative regulation of gene expression | 1.20E-08 | GO:0023051 | regulation of signaling | 0.00788 |
| GO:0015833 | peptide transport | 1.30E-08 | GO:0045765 | regulation of angiogenesis | 0.00894 |
| GO:1902600 | hydrogen ion transmembrane transport | 1.40E-08 | GO:0043244 | regulation of protein complex disassembl... | 0.00926 |
| GO:0015980 | energy derivation by oxidation of organi... | 1.40E-08 | GO:0009409 | response to cold | 0.00941 |
| GO:0015031 | protein transport | 1.40E-08 | GO:0043331 | response to dsRNA | 0.00941 |
| GO:0042886 | amide transport | 1.80E-08 | GO:1902041 | regulation of extrinsic apoptotic signal... | 0.00941 |
| GO:0019693 | ribose phosphate metabolic process | 3.00E-08 | GO:0042035 | regulation of cytokine biosynthetic proc... | 0.00941 |
| GO:0044248 | cellular catabolic process | 4.20E-08 | GO:0032675 | regulation of interleukin-6 production | 0.00941 |
| GO:0055086 | nucleobase-containing small molecule met... | 5.00E-08 | GO:0010243 | response to organonitrogen compound | 0.00949 |
| GO:0000028 | ribosomal small subunit assembly | 5.20E-08 | GO:0007015 | actin filament organization | 0.00971 |
| GO:0006753 | nucleoside phosphate metabolic process | 5.60E-08 | GO:0050794 | regulation of cellular process | 0.01035 |
| GO:0009117 | nucleotide metabolic process | 5.60E-08 | GO:0030029 | actin filament-based process | 0.01059 |
| GO:1901575 | organic substance catabolic process | 6.20E-08 | GO:0030036 | actin cytoskeleton organization | 0.01059 |
| GO:0033108 | mitochondrial respiratory chain complex ... | 1.20E-07 | GO:2001240 | negative regulation of extrinsic apoptot... | 0.01106 |
| GO:0010605 | negative regulation of macromolecule met... | 1.40E-07 | GO:0035924 | cellular response to vascular endothelia... | 0.01106 |
| GO:0045184 | establishment of protein localization | 1.60E-07 | GO:0071801 | regulation of podosome assembly | 0.01106 |
| GO:0006810 | transport | 1.70E-07 | GO:0010880 | regulation of release of sequestered cal... | 0.01106 |
| GO:0006414 | translational elongation | 2.50E-07 | GO:0070816 | phosphorylation of RNA polymerase II C-t... | 0.01106 |
| GO:0051234 | establishment of localization | 2.60E-07 | GO:1901099 | negative regulation of signal transducti... | 0.01106 |
| GO:0043933 | macromolecular complex subunit organizat... | 3.20E-07 | GO:0071902 | positive regulation of protein serine/th... | 0.0112 |
| GO:0071705 | nitrogen compound transport | 3.30E-07 | GO:0051646 | mitochondrion localization | 0.01145 |
| GO:0009892 | negative regulation of metabolic process | 5.30E-07 | GO:0051279 | regulation of release of sequestered cal... | 0.01145 |
| GO:0006120 | mitochondrial electron transport, NADH t... | 6.20E-07 | GO:0070848 | response to growth factor | 0.01173 |
| GO:0034622 | cellular macromolecular complex assembly | 6.70E-07 | GO:0050789 | regulation of biological process | 0.01193 |
| GO:0034613 | cellular protein localization | 7.20E-07 | GO:0034614 | cellular response to reactive oxygen spe... | 0.01288 |
| GO:0015992 | proton transport | 7.30E-07 | GO:0007286 | spermatid development | 0.01306 |
| GO:1901564 | organonitrogen compound metabolic proces... | 1.10E-06 | GO:0048771 | tissue remodeling | 0.01306 |
| GO:0070727 | cellular macromolecule localization | 1.20E-06 | GO:0007043 | cell-cell junction assembly | 0.01335 |
| GO:0009056 | catabolic process | 1.30E-06 | GO:0032635 | interleukin-6 production | 0.01335 |
| GO:0006818 | hydrogen transport | 1.40E-06 | GO:0007249 | I-kappaB kinase/NF-kappaB signaling | 0.01381 |
| GO:0071702 | organic substance transport | 1.50E-06 | GO:0045944 | positive regulation of transcription fro... | 0.01407 |
| GO:0007005 | mitochondrion organization | 1.70E-06 | GO:0097435 | supramolecular fiber organization | 0.0145 |
| GO:0051641 | cellular localization | 2.00E-06 | GO:1902680 | positive regulation of RNA biosynthetic ... | 0.0149 |
| GO:0006123 | mitochondrial electron transport, cytoch... | 5.30E-06 | GO:0045893 | positive regulation of transcription, DN... | 0.0149 |
| GO:0006091 | generation of precursor metabolites and ... | 5.70E-06 | GO:1903508 | positive regulation of nucleic acid-temp... | 0.0149 |
| GO:0032981 | mitochondrial respiratory chain complex ... | 9.00E-06 | GO:0043484 | regulation of RNA splicing | 0.01499 |
| GO:0097031 | mitochondrial respiratory chain complex ... | 9.00E-06 | GO:0072359 | circulatory system development | 0.0156 |
| GO:0010257 | NADH dehydrogenase complex assembly | 9.00E-06 | GO:0070588 | calcium ion transmembrane transport | 0.01576 |
| GO:0000027 | ribosomal large subunit assembly | 1.90E-05 | GO:0007204 | positive regulation of cytosolic calcium... | 0.01576 |
| GO:0008104 | protein localization | 2.30E-05 | GO:0071396 | cellular response to lipid | 0.01614 |
| GO:0070125 | mitochondrial translational elongation | 3.90E-05 | GO:0048545 | response to steroid hormone | 0.01679 |
| GO:0006725 | cellular aromatic compound metabolic pro... | 5.10E-05 | GO:0048646 | anatomical structure formation involved ... | 0.01696 |
| GO:0070126 | mitochondrial translational termination | 5.40E-05 | GO:0070301 | cellular response to hydrogen peroxide | 0.01698 |
| GO:0065003 | macromolecular complex assembly | 5.40E-05 | GO:0060402 | calcium ion transport into cytosol | 0.01698 |
| GO:0051179 | localization | 5.60E-05 | GO:0048146 | positive regulation of fibroblast prolif... | 0.01746 |
| GO:0033036 | macromolecule localization | 6.00E-05 | GO:0032755 | positive regulation of interleukin-6 pro... | 0.01746 |
| GO:0034641 | cellular nitrogen compound metabolic pro... | 8.90E-05 | GO:0044236 | multicellular organism metabolic process | 0.01746 |
| GO:0032543 | mitochondrial translation | 9.00E-05 | GO:0042108 | positive regulation of cytokine biosynth... | 0.01746 |
| GO:0055114 | oxidation-reduction process | 9.70E-05 | GO:0043550 | regulation of lipid kinase activity | 0.01746 |
| GO:0046483 | heterocycle metabolic process | 0.00014 | GO:0071375 | cellular response to peptide hormone sti... | 0.01822 |
| GO:1901360 | organic cyclic compound metabolic proces... | 0.00015 | GO:1903169 | regulation of calcium ion transmembrane ... | 0.01827 |
| GO:0006139 | nucleobase-containing compound metabolic... | 0.00017 | GO:0043242 | negative regulation of protein complex d... | 0.01827 |
| GO:0042407 | cristae formation | 0.00024 | GO:0051017 | actin filament bundle assembly | 0.01827 |
| GO:0007007 | inner mitochondrial membrane organizatio... | 0.00024 | GO:0006360 | transcription from RNA polymerase I prom... | 0.01827 |
| GO:0006415 | translational termination | 0.00024 | GO:0040013 | negative regulation of locomotion | 0.01876 |
| GO:0140053 | mitochondrial gene expression | 0.00026 | GO:0007281 | germ cell development | 0.01907 |
| GO:0017004 | cytochrome complex assembly | 0.00029 | GO:0032868 | response to insulin | 0.01965 |
| GO:0022607 | cellular component assembly | 0.00031 | GO:0071310 | cellular response to organic substance | 0.01978 |
| GO:0048519 | negative regulation of biological proces... | 0.00031 | GO:0002385 | mucosal immune response | 0.02034 |
| GO:0019637 | organophosphate metabolic process | 0.00033 | GO:0014808 | release of sequestered calcium ion into ... | 0.02034 |
| GO:0030490 | maturation of SSU-rRNA | 0.0004 | GO:0035082 | axoneme assembly | 0.02034 |
| GO:0045116 | protein neddylation | 0.00063 | GO:0007140 | male meiotic nuclear division | 0.02034 |
| GO:0051444 | negative regulation of ubiquitin-protein... | 0.00072 | GO:0051654 | establishment of mitochondrion localizat... | 0.02034 |
| GO:0043624 | cellular protein complex disassembly | 0.00075 | GO:0010761 | fibroblast migration | 0.02034 |
| GO:0071822 | protein complex subunit organization | 0.00086 | GO:0002756 | MyD88-independent toll-like receptor sig... | 0.02034 |
| GO:1904667 | negative regulation of ubiquitin protein... | 0.00103 | GO:0034643 | establishment of mitochondrion localizat... | 0.02034 |
| GO:0031397 | negative regulation of protein ubiquitin... | 0.00105 | GO:1903514 | release of sequestered calcium ion into ... | 0.02034 |
| GO:0000470 | maturation of LSU-rRNA | 0.00137 | GO:0070296 | sarcoplasmic reticulum calcium ion trans... | 0.02034 |

| | | | | | |
|---|---|---|---|---|---|
| GO:0007006 | mitochondrial membrane organization | 0.00139 | GO:0010591 | regulation of lamellipodium assembly | 0.02034 |
| GO:0001836 | release of cytochrome c from mitochondri... | 0.00149 | GO:0032570 | response to progesterone | 0.02034 |
| GO:0070925 | organelle assembly | 0.0015 | GO:0035272 | exocrine system development | 0.02034 |
| GO:1903321 | negative regulation of protein modificat... | 0.00185 | GO:0047497 | mitochondrion transport along microtubul... | 0.02034 |
| GO:0051438 | regulation of ubiquitin-protein transfer... | 0.00195 | GO:0007019 | microtubule depolymerization | 0.02034 |
| GO:2001244 | positive regulation of intrinsic apoptot... | 0.00209 | GO:0045912 | negative regulation of carbohydrate meta... | 0.02034 |
| GO:0050954 | sensory perception of mechanical stimulu... | 0.00209 | GO:0002920 | regulation of humoral immune response | 0.02034 |
| GO:2000059 | negative regulation of protein ubiquitin... | 0.00222 | GO:0035666 | TRIF-dependent toll-like receptor signal... | 0.02034 |
| GO:0002396 | MHC protein complex assembly | 0.00222 | GO:0009967 | positive regulation of signal transducti... | 0.0204 |
| GO:1901135 | carbohydrate derivative metabolic proces... | 0.0024 | GO:0065009 | regulation of molecular function | 0.02143 |
| GO:0043623 | cellular protein complex assembly | 0.00252 | GO:0048515 | spermatid differentiation | 0.02164 |
| GO:2000058 | regulation of protein ubiquitination inv... | 0.00255 | GO:0051494 | negative regulation of cytoskeleton orga... | 0.02164 |
| GO:1904666 | regulation of ubiquitin protein ligase a... | 0.00255 | GO:0045935 | positive regulation of nucleobase-contai... | 0.02171 |
| GO:0042776 | mitochondrial ATP synthesis coupled prot... | 0.00268 | GO:0048583 | regulation of response to stimulus | 0.02208 |
| GO:0009987 | cellular process | 0.00324 | GO:0019932 | second-messenger-mediated signaling | 0.02212 |
| GO:0015985 | energy coupled proton transport, down el... | 0.00403 | GO:0033993 | response to lipid | 0.02242 |
| GO:0015986 | ATP synthesis coupled proton transport | 0.00403 | GO:1901136 | carbohydrate derivative catabolic proces... | 0.02285 |
| GO:0007605 | sensory perception of sound | 0.00416 | GO:0045926 | negative regulation of growth | 0.02285 |
| GO:0032984 | macromolecular complex disassembly | 0.00454 | GO:0007266 | Rho protein signal transduction | 0.02287 |
| GO:0008535 | respiratory chain complex IV assembly | 0.00461 | GO:0032970 | regulation of actin filament-based proce... | 0.02362 |
| GO:0015672 | monovalent inorganic cation transport | 0.00501 | GO:0022612 | gland morphogenesis | 0.02425 |
| GO:0000462 | maturation of SSU-rRNA from tricistronic... | 0.00554 | GO:0061572 | actin filament bundle organization | 0.02425 |
| GO:0003407 | neural retina development | 0.00561 | GO:0048584 | positive regulation of response to stimu... | 0.02454 |
| GO:0031146 | SCF-dependent proteasomal ubiquitin-depe... | 0.00582 | GO:0098542 | defense response to other organism | 0.02465 |
| GO:0000387 | spliceosomal snRNP assembly | 0.00585 | GO:0009617 | response to bacterium | 0.02465 |
| GO:0044281 | small molecule metabolic process | 0.00692 | GO:0031663 | lipopolysaccharide-mediated signaling pa... | 0.02511 |
| GO:0043241 | protein complex disassembly | 0.00712 | GO:0032722 | positive regulation of chemokine product... | 0.02511 |
| GO:0044271 | cellular nitrogen compound biosynthetic ... | 0.00739 | GO:0003151 | outflow tract morphogenesis | 0.02511 |
| GO:0016043 | cellular component organization | 0.00763 | GO:0070265 | necrotic cell death | 0.02511 |
| GO:0050852 | T cell receptor signaling pathway | 0.00907 | GO:0010628 | positive regulation of gene expression | 0.02539 |
| GO:0033617 | mitochondrial respiratory chain complex ... | 0.00985 | GO:0002237 | response to molecule of bacterial origin | 0.02579 |
| GO:0008543 | fibroblast growth factor receptor signal... | 0.01046 | GO:0008154 | actin polymerization or depolymerization | 0.02589 |
| GO:0060041 | retina development in camera-type eye | 0.01127 | GO:0097190 | apoptotic signaling pathway | 0.0261 |
| GO:0043928 | exonucleolytic nuclear-transcribed mRNA ... | 0.01147 | GO:0009968 | negative regulation of signal transducti... | 0.02671 |
| GO:0042461 | photoreceptor cell development | 0.0121 | GO:0045860 | positive regulation of protein kinase ac... | 0.02707 |
| GO:0045471 | response to ethanol | 0.01219 | GO:0001843 | neural tube closure | 0.02709 |
| GO:0042787 | protein ubiquitination involved in ubiqu... | 0.01223 | GO:0060606 | tube closure | 0.02709 |
| GO:0044267 | cellular protein metabolic process | 0.01331 | GO:0014074 | response to purine-containing compound | 0.02709 |
| GO:0099132 | ATP hydrolysis coupled cation transmembr... | 0.01366 | GO:0014020 | primary neural tube formation | 0.02709 |
| GO:0031396 | regulation of protein ubiquitination | 0.01469 | GO:0008064 | regulation of actin polymerization or de... | 0.02714 |
| GO:0098662 | inorganic cation transmembrane transport | 0.01533 | GO:0030832 | regulation of actin filament length | 0.02714 |
| GO:0006370 | 7-methylguanosine mRNA capping | 0.01585 | GO:0043620 | regulation of DNA-templated transcriptio... | 0.02714 |
| GO:0006122 | mitochondrial electron transport, ubiqui... | 0.01585 | GO:0051480 | regulation of cytosolic calcium ion conc... | 0.02714 |
| GO:0036260 | RNA capping | 0.01585 | GO:0043434 | response to peptide hormone | 0.02743 |
| GO:0042073 | intraciliary transport | 0.01585 | GO:0051235 | maintenance of location | 0.02798 |
| GO:0009452 | 7-methylguanosine RNA capping | 0.01585 | GO:0032956 | regulation of actin cytoskeleton organiz... | 0.02933 |
| GO:0097034 | mitochondrial respiratory chain complex ... | 0.01585 | GO:0032869 | cellular response to insulin stimulus | 0.03009 |
| GO:0006749 | glutathione metabolic process | 0.01623 | GO:0046777 | protein autophosphorylation | 0.03009 |
| GO:0042440 | pigment metabolic process | 0.01623 | GO:0007167 | enzyme linked receptor protein signaling... | 0.03056 |
| GO:0000291 | nuclear-transcribed mRNA catabolic proce... | 0.01674 | GO:0097553 | calcium ion transmembrane import into cy... | 0.03136 |
| GO:2000116 | regulation of cysteine-type endopeptidas... | 0.01693 | GO:0051282 | regulation of sequestering of calcium io... | 0.03136 |
| GO:0099131 | ATP hydrolysis coupled ion transmembrane... | 0.01752 | GO:0051283 | negative regulation of sequestering of c... | 0.03136 |
| GO:1903363 | negative regulation of cellular protein ... | 0.01752 | GO:0051208 | sequestering of calcium ion | 0.03136 |
| GO:0051436 | negative regulation of ubiquitin-protein... | 0.01793 | GO:0051209 | release of sequestered calcium ion into ... | 0.03136 |
| GO:0051437 | positive regulation of ubiquitin-protein... | 0.01793 | GO:0009266 | response to temperature stimulus | 0.03203 |
| GO:0051439 | regulation of ubiquitin-protein ligase a... | 0.01793 | GO:0007184 | SMAD protein import into nucleus | 0.03274 |
| GO:0031145 | anaphase-promoting complex-dependent cat... | 0.01793 | GO:1900543 | negative regulation of purine nucleotide... | 0.03274 |
| GO:0051348 | negative regulation of transferase activ... | 0.01809 | GO:0002251 | organ or tissue specific immune response | 0.03274 |
| GO:1902749 | regulation of cell cycle G2/M phase tran... | 0.01872 | GO:0071549 | cellular response to dexamethasone stimu... | 0.03274 |

Supplementary Table S2

| GO Term Pathway: NesCreCsf1flfl_Cb_P8 | GO Term_ID | overUnder | pvalue |
|---|---|---|---|
| regulation of generation of precursor metabolites and energy | GO:0043467 | under | 0.00337 |
| regulation of tissue remodeling | GO:0034103 | under | 0.00412 |
| sensory perception of mechanical stimulus | GO:0050954 | under | 0.00445 |
| regulation of phosphatase activity | GO:0010921 | under | 0.00778 |
| ensheathment of neurons | GO:0007272 | under | 0.00983 |
| myelination | GO:0042552 | under | 0.00983 |
| axon ensheathment | GO:0008366 | under | 0.00983 |
| regulation of coenzyme metabolic process | GO:0051196 | under | 0.00987 |
| regulation of cofactor metabolic process | GO:0051193 | under | 0.00987 |
| regulation of carbohydrate catabolic process | GO:0043470 | under | 0.00987 |
| sensory perception of sound | GO:0007605 | under | 0.0127 |
| negative regulation of protein complex assembly | GO:0031333 | under | 0.0143 |
| regulation of STAT cascade | GO:1904892 | under | 0.0145 |
| regulation of JAK-STAT cascade | GO:0046425 | under | 0.0145 |
| multi-organism reproductive process | GO:0044703 | under | 0.016 |
| response to endogenous stimulus | GO:0009719 | under | 0.0166 |
| regulation of bone resorption | GO:0045124 | under | 0.0174 |
| regulation of bone remodeling | GO:0046850 | under | 0.0174 |
| regulation of ATP metabolic process | GO:1903578 | under | 0.018 |
| nephron epithelium morphogenesis | GO:0072088 | under | 0.0184 |
| nephron tubule development | GO:0072080 | under | 0.0184 |
| nephron tubule morphogenesis | GO:0072078 | under | 0.0184 |
| kidney epithelium development | GO:0072073 | under | 0.0184 |
| nephron morphogenesis | GO:0072028 | under | 0.0184 |
| nephron epithelium development | GO:0072009 | under | 0.0184 |
| renal tubule morphogenesis | GO:0061333 | under | 0.0184 |
| renal tubule development | GO:0061326 | under | 0.0184 |
| ribonucleotide metabolic process | GO:0009259 | under | 0.0205 |
| purine ribonucleotide metabolic process | GO:0009150 | under | 0.0205 |
| ribose phosphate metabolic process | GO:0019693 | under | 0.0205 |
| cellular response to endogenous stimulus | GO:0071495 | under | 0.0218 |
| response to peptide hormone | GO:0043434 | under | 0.0221 |
| positive regulation of bone remodeling | GO:0046852 | under | 0.0223 |
| positive regulation of bone resorption | GO:0045780 | under | 0.0223 |
| positive regulation of tissue remodeling | GO:0034105 | under | 0.0223 |
| cranial nerve development | GO:0021545 | under | 0.0223 |
| parasympathetic nervous system development | GO:0048486 | under | 0.0223 |
| preganglionic parasympathetic fiber development | GO:0021783 | under | 0.0223 |
| regulation of tyrosine phosphorylation of STAT protein | GO:0042509 | under | 0.0225 |
| carbohydrate transport | GO:0008643 | under | 0.0233 |
| vesicle cytoskeletal trafficking | GO:0099518 | under | 0.0243 |
| sexual reproduction | GO:0019953 | under | 0.025 |
| purine nucleotide metabolic process | GO:0006163 | under | 0.0251 |
| purine-containing compound metabolic process | GO:0072521 | under | 0.0251 |
| muscle fiber development | GO:0048747 | under | 0.0259 |
| response to insulin | GO:0032868 | under | 0.0261 |
| nucleobase-containing small molecule metabolic process | GO:0055086 | under | 0.0262 |
| negative regulation of hydrolase activity | GO:0051346 | under | 0.0263 |
| regulation of glycolytic process | GO:0006110 | under | 0.0297 |
| heparin metabolic process | GO:0030202 | under | 0.0305 |
| hexose transport | GO:0008645 | under | 0.0307 |
| glucose transport | GO:0015758 | under | 0.0307 |
| monosaccharide transport | GO:0015749 | under | 0.0307 |
| male sex differentiation | GO:0046661 | under | 0.0315 |
| positive regulation of ATP metabolic process | GO:1903580 | under | 0.0315 |
| nerve development | GO:0021675 | under | 0.033 |
| regulation of carbohydrate metabolic process | GO:0006109 | under | 0.0333 |
| negative regulation of cellular protein localization | GO:1903828 | under | 0.0339 |
| negative regulation of peptidyl-tyrosine phosphorylation | GO:0050732 | under | 0.0339 |

| | | | |
|---|---|---|---|
| digestive system development | GO:0055123 | under | 0.0345 |
| small molecule metabolic process | GO:0044281 | under | 0.0352 |
| regulation of cellular carbohydrate metabolic process | GO:0010675 | under | 0.0369 |
| positive regulation of mitochondrial translation | GO:0070131 | under | 0.0373 |
| regulation of mitochondrial translation | GO:0070129 | under | 0.0373 |
| cellular response to hormone stimulus | GO:0032870 | under | 0.0381 |
| monocarboxylic acid metabolic process | GO:0032787 | under | 0.0382 |
| response to hormone | GO:0009725 | under | 0.0387 |
| purine ribonucleoside triphosphate metabolic process | GO:0009205 | under | 0.0403 |
| ribonucleoside triphosphate metabolic process | GO:0009199 | under | 0.0403 |
| negative regulation of cellular component organization | GO:0051129 | under | 0.0407 |
| proteoglycan biosynthetic process | GO:0030166 | under | 0.0416 |
| positive regulation of interleukin-5 production | GO:0032754 | under | 0.0423 |
| positive regulation of interleukin-13 production | GO:0032736 | under | 0.0423 |
| regulation of interleukin-5 production | GO:0032674 | under | 0.0423 |
| regulation of interleukin-13 production | GO:0032656 | under | 0.0423 |
| organic acid metabolic process | GO:0006082 | under | 0.044 |
| oxoacid metabolic process | GO:0043436 | under | 0.044 |
| anatomical structure morphogenesis | GO:0009653 | under | 0.0451 |
| negative regulation of phosphatase activity | GO:0010923 | under | 0.0452 |
| negative regulation of dephosphorylation | GO:0035305 | under | 0.0452 |
| positive regulation of tyrosine phosphorylation of STAT protein | GO:0042531 | under | 0.0466 |
| positive regulation of STAT cascade | GO:1904894 | under | 0.0466 |
| positive regulation of JAK-STAT cascade | GO:0046427 | under | 0.0466 |
| establishment of vesicle localization | GO:0051650 | under | 0.0467 |
| vesicle localization | GO:0051648 | under | 0.0467 |
| nucleoside phosphate metabolic process | GO:0006753 | under | 0.0469 |
| nucleotide metabolic process | GO:0009117 | under | 0.0469 |
| positive regulation of lipase activity | GO:0060193 | under | 0.0475 |
| regulation of lipase activity | GO:0060191 | under | 0.0475 |
| purine nucleoside triphosphate metabolic process | GO:0009144 | under | 0.0497 |
| nucleoside triphosphate metabolic process | GO:0009141 | under | 0.0497 |
| rhythmic behavior | GO:0007622 | under | 0.0498 |
| regulation of protein acetylation | GO:1901983 | over | 0.00256 |
| adaptive immune response | GO:0002250 | over | 0.00264 |
| cell cycle checkpoint | GO:0000075 | over | 0.00336 |
| immune response | GO:0006955 | over | 0.00656 |
| lymphocyte activation | GO:0046649 | over | 0.00717 |
| DNA damage checkpoint | GO:0000077 | over | 0.0081 |
| DNA integrity checkpoint | GO:0031570 | over | 0.0081 |
| production of molecular mediator of immune response | GO:0002440 | over | 0.00825 |
| regulation of peptidyl-lysine acetylation | GO:2000756 | over | 0.00848 |
| leukocyte activation | GO:0045321 | over | 0.00945 |
| positive regulation of protein acetylation | GO:1901985 | over | 0.0102 |
| positive regulation of leukocyte chemotaxis | GO:0002690 | over | 0.0105 |
| mitotic cell cycle checkpoint | GO:0007093 | over | 0.0112 |
| immunoglobulin mediated immune response | GO:0016064 | over | 0.0128 |
| B cell mediated immunity | GO:0019724 | over | 0.0128 |
| positive regulation of neuron death | GO:1901216 | over | 0.0131 |
| defense response | GO:0006952 | over | 0.0136 |
| lymphocyte differentiation | GO:0030098 | over | 0.0137 |
| regulation of viral life cycle | GO:1903900 | over | 0.0164 |
| regulation of leukocyte chemotaxis | GO:0002688 | over | 0.0165 |
| membrane lipid biosynthetic process | GO:0046467 | over | 0.018 |
| regulation of symbiosis, encompassing mutualism through parasitism | GO:0043903 | over | 0.0186 |
| regulation of viral process | GO:0050792 | over | 0.0186 |
| regulation of small GTPase mediated signal transduction | GO:0051056 | over | 0.019 |
| humoral immune response | GO:0006959 | over | 0.0202 |
| regulation of cation channel activity | GO:2001257 | over | 0.0216 |
| regulation of multi-organism process | GO:0043900 | over | 0.0221 |

| | | | |
|---|---|---|---|
| cellular response to DNA damage stimulus | GO:0006974 | over | 0.0229 |
| regulation of protein export from nucleus | GO:0046825 | over | 0.023 |
| regulation of granulocyte chemotaxis | GO:0071622 | over | 0.0233 |
| mitotic DNA integrity checkpoint | GO:0044774 | over | 0.0237 |
| mitotic DNA damage checkpoint | GO:0044773 | over | 0.0237 |
| positive regulation of hydrogen peroxide-mediated programmed cell death | GO:1901300 | over | 0.0238 |
| regulation of hydrogen peroxide-mediated programmed cell death | GO:1901298 | over | 0.0238 |
| positive regulation of response to oxidative stress | GO:1902884 | over | 0.0238 |
| positive regulation of response to reactive oxygen species | GO:1901033 | over | 0.0238 |
| regulation of response to reactive oxygen species | GO:1901031 | over | 0.0238 |
| positive regulation of oxidative stress-induced cell death | GO:1903209 | over | 0.0238 |
| regulation of hydrogen peroxide-induced cell death | GO:1903205 | over | 0.0238 |
| positive regulation of cellular response to oxidative stress | GO:1900409 | over | 0.0238 |
| positive regulation of hydrogen peroxide-induced cell death | GO:1905206 | over | 0.0238 |
| positive regulation of cellular response to drug | GO:2001040 | over | 0.0238 |
| regulation of cellular response to drug | GO:2001038 | over | 0.0238 |
| positive regulation of response to drug | GO:2001025 | over | 0.0238 |
| positive regulation of chemotaxis | GO:0050921 | over | 0.0243 |
| DNA metabolic process | GO:0006259 | over | 0.0256 |
| negative regulation of chromosome organization | GO:2001251 | over | 0.0261 |
| neuronal stem cell population maintenance | GO:0097150 | over | 0.0267 |
| regulation of signal transduction by p53 class mediator | GO:1901796 | over | 0.0269 |
| negative regulation of cell cycle G2/M phase transition | GO:1902750 | over | 0.0277 |
| G2 DNA damage checkpoint | GO:0031572 | over | 0.0277 |
| amine metabolic process | GO:0009308 | over | 0.0285 |
| cellular biogenic amine metabolic process | GO:0006576 | over | 0.0285 |
| cellular amine metabolic process | GO:0044106 | over | 0.0285 |
| regulation of histone acetylation | GO:0035065 | over | 0.0296 |
| lymphoid progenitor cell differentiation | GO:0002320 | over | 0.0298 |
| primary alcohol metabolic process | GO:0034308 | over | 0.0298 |
| adaptive immune response based on somatic recombination of immune receptors built | GO:0002460 | over | 0.0305 |
| lymphocyte mediated immunity | GO:0002449 | over | 0.0308 |
| regulation of chemotaxis | GO:0050920 | over | 0.0317 |
| immune system development | GO:0002520 | over | 0.0319 |
| oligosaccharide biosynthetic process | GO:0009312 | over | 0.0322 |
| oligosaccharide metabolic process | GO:0009311 | over | 0.0322 |
| protein N-linked glycosylation via asparagine | GO:0018279 | over | 0.0322 |
| peptidyl-asparagine modification | GO:0018196 | over | 0.0322 |
| response to osmotic stress | GO:0006970 | over | 0.0335 |
| positive regulation of peptidyl-lysine acetylation | GO:2000758 | over | 0.0354 |
| stem cell population maintenance | GO:0019827 | over | 0.036 |
| humoral immune response mediated by circulating immunoglobulin | GO:0002455 | over | 0.0362 |
| immunoglobulin production involved in immunoglobulin mediated immune response | GO:0002381 | over | 0.0363 |
| immunoglobulin production | GO:0002377 | over | 0.0363 |
| somatic recombination of immunoglobulin gene segments | GO:0016447 | over | 0.0363 |
| somatic diversification of immunoglobulins | GO:0016445 | over | 0.0363 |
| somatic cell DNA recombination | GO:0016444 | over | 0.0363 |
| somatic diversification of immunoglobulins involved in immune response | GO:0002208 | over | 0.0363 |
| somatic recombination of immunoglobulin genes involved in immune response | GO:0002204 | over | 0.0363 |
| somatic diversification of immune receptors | GO:0002200 | over | 0.0363 |
| isotype switching | GO:0045190 | over | 0.0363 |
| somatic diversification of immune receptors via germline recombination within a single | GO:0002562 | over | 0.0363 |
| positive regulation of viral life cycle | GO:1903902 | over | 0.0373 |
| glycolipid biosynthetic process | GO:0009247 | over | 0.0378 |
| signal transduction involved in DNA damage checkpoint | GO:0072422 | over | 0.0385 |
| signal transduction involved in DNA integrity checkpoint | GO:0072401 | over | 0.0385 |
| signal transduction involved in cell cycle checkpoint | GO:0072395 | over | 0.0385 |
| smooth muscle tissue development | GO:0048745 | over | 0.0389 |
| positive regulation of granulocyte chemotaxis | GO:0071624 | over | 0.0391 |
| positive regulation of neutrophil migration | GO:1902624 | over | 0.0391 |

| | | | |
|---|---|---|---|
| regulation of neutrophil migration | GO:1902622 | over | 0.0391 |
| positive regulation of neutrophil chemotaxis | GO:0090023 | over | 0.0391 |
| regulation of neutrophil chemotaxis | GO:0090022 | over | 0.0391 |
| B cell activation involved in immune response | GO:0002312 | over | 0.0392 |
| ovulation cycle | GO:0042698 | over | 0.0392 |
| response to stress | GO:0006950 | over | 0.0404 |
| liposaccharide metabolic process | GO:1903509 | over | 0.0404 |
| cardiac atrium development | GO:0003230 | over | 0.0408 |
| cardiac atrium morphogenesis | GO:0003209 | over | 0.0408 |
| regulation of Ras protein signal transduction | GO:0046578 | over | 0.0413 |
| leukocyte mediated immunity | GO:0002443 | over | 0.0436 |
| mitotic G1/S transition checkpoint | GO:0044819 | over | 0.0441 |
| positive regulation of protein export from nucleus | GO:0046827 | over | 0.0466 |
| negative regulation of viral genome replication | GO:0045071 | over | 0.0472 |
| regulation of viral genome replication | GO:0045069 | over | 0.0472 |
| glycolipid metabolic process | GO:0006664 | over | 0.0479 |
| negative regulation of viral process | GO:0048525 | over | 0.0494 |
| positive regulation of response to external stimulus | GO:0032103 | over | 0.0496 |

Supplementary Table S2, cont.

| GO Term Pathway: NesCreCsf1flfl_Fb_P8 | GO Term_ID | overUnder | pvalue |
|---|---|---|---|
| protein oligomerization | GO:0051259 | under | 0.00269 |
| protein complex assembly | GO:0006461 | under | 0.00309 |
| protein complex biogenesis | GO:0070271 | under | 0.00309 |
| response to wounding | GO:0009611 | under | 0.00389 |
| antigen processing and presentation of peptide antigen via MHC class II | GO:0002495 | under | 0.00429 |
| antigen processing and presentation of exogenous peptide antigen via MHC class II | GO:0019886 | under | 0.00429 |
| antigen processing and presentation of peptide or polysaccharide antigen via MHC class II | GO:0002504 | under | 0.00429 |
| multicellular organismal signaling | GO:0035637 | under | 0.00504 |
| regeneration | GO:0031099 | under | 0.00533 |
| protein complex subunit organization | GO:0071822 | under | 0.00604 |
| wound healing | GO:0042060 | under | 0.00704 |
| lactation | GO:0007595 | under | 0.00761 |
| body fluid secretion | GO:0007589 | under | 0.00761 |
| animal organ morphogenesis | GO:0009887 | under | 0.00869 |
| monocarboxylic acid metabolic process | GO:0032787 | under | 0.00971 |
| regulation of lipid transport | GO:0032368 | over | 0.0106 |
| protein acetylation | GO:0006473 | over | 0.0112 |
| protein acylation | GO:0043543 | over | 0.0112 |
| sensory perception of pain | GO:0019233 | under | 0.0121 |
| cation transport | GO:0006812 | under | 0.014 |
| negative regulation of tumor necrosis factor production | GO:0032720 | over | 0.0145 |
| negative regulation of tumor necrosis factor superfamily cytokine production | GO:1903556 | over | 0.0145 |
| antigen processing and presentation of exogenous peptide antigen | GO:0002478 | under | 0.015 |
| antigen processing and presentation of exogenous antigen | GO:0019884 | under | 0.015 |
| tissue remodeling | GO:0048771 | under | 0.015 |
| positive regulation of cellular catabolic process | GO:0031331 | over | 0.0154 |
| mRNA metabolic process | GO:0016071 | over | 0.0166 |
| macromolecular complex assembly | GO:0065003 | under | 0.0186 |
| positive regulation of inflammatory response | GO:0050729 | under | 0.0202 |
| oligodendrocyte differentiation | GO:0048709 | under | 0.0205 |
| positive regulation of toll-like receptor signaling pathway | GO:0034123 | under | 0.0205 |
| macromolecule catabolic process | GO:0009057 | over | 0.0212 |
| female gamete generation | GO:0007292 | over | 0.0224 |
| system process | GO:0003008 | under | 0.023 |
| cardiac conduction | GO:0061337 | under | 0.0231 |
| anatomical structure formation involved in morphogenesis | GO:0048646 | under | 0.0244 |
| regulation of macroautophagy | GO:0016241 | over | 0.0251 |
| positive regulation of macroautophagy | GO:0016239 | over | 0.0251 |
| positive regulation of autophagy | GO:0010508 | over | 0.0251 |
| nervous system process | GO:0050877 | under | 0.0254 |
| substrate adhesion-dependent cell spreading | GO:0034446 | under | 0.0259 |
| metal ion homeostasis | GO:0055065 | under | 0.0261 |
| negative regulation of endothelial cell proliferation | GO:0001937 | under | 0.0262 |
| regulation of membrane repolarization | GO:0060306 | under | 0.0262 |
| homeostatic process | GO:0042592 | under | 0.0266 |
| divalent inorganic cation homeostasis | GO:0072507 | under | 0.0275 |
| endoderm formation | GO:0001706 | under | 0.0278 |
| negative regulation of Wnt signaling pathway | GO:0030178 | under | 0.0278 |
| negative regulation of canonical Wnt signaling pathway | GO:0090090 | under | 0.0278 |
| endodermal cell differentiation | GO:0035987 | under | 0.0278 |
| endoderm development | GO:0007492 | under | 0.0278 |
| antigen processing and presentation of peptide antigen | GO:0048002 | under | 0.0278 |
| protein tetramerization | GO:0051262 | under | 0.028 |
| macromolecular complex subunit organization | GO:0043933 | under | 0.0283 |
| peptide cross-linking | GO:0018149 | under | 0.0286 |
| cell projection assembly | GO:0030031 | over | 0.0289 |
| heterophilic cell-cell adhesion via plasma membrane cell adhesion molecules | GO:0007157 | over | 0.0294 |
| liver development | GO:0001889 | under | 0.0301 |
| hepaticobiliary system development | GO:0061008 | under | 0.0301 |
| double-strand break repair via nonhomologous end joining | GO:0006303 | over | 0.031 |
| skeletal muscle organ development | GO:0060538 | under | 0.0311 |
| cardiovascular system development | GO:0072358 | under | 0.0315 |
| nucleoside triphosphate metabolic process | GO:0009141 | under | 0.0316 |
| cochlea development | GO:0090102 | under | 0.0316 |
| cation homeostasis | GO:0055080 | under | 0.0316 |
| inorganic ion homeostasis | GO:0098771 | under | 0.0316 |
| animal organ regeneration | GO:0031100 | under | 0.032 |
| peptidyl-lysine acetylation | GO:0018394 | over | 0.0328 |
| negative regulation of NF-kappaB transcription factor activity | GO:0032088 | over | 0.0339 |
| negative regulation of DNA binding transcription factor activity | GO:0043433 | over | 0.0339 |
| response to mechanical stimulus | GO:0009612 | under | 0.0341 |
| positive regulation of amine transport | GO:0051954 | under | 0.0343 |
| myotube differentiation | GO:0014902 | under | 0.0347 |

| | | | |
|---|---|---|---|
| regulation of autophagy | GO:0010506 | over | 0.0349 |
| cellular response to estradiol stimulus | GO:0071392 | over | 0.0355 |
| mammary gland epithelium development | GO:0061180 | over | 0.0355 |
| anatomical structure morphogenesis | GO:0009653 | under | 0.0358 |
| cellular calcium ion homeostasis | GO:0006874 | under | 0.0359 |
| calcium ion homeostasis | GO:0055074 | under | 0.0359 |
| response to ischemia | GO:0002931 | under | 0.0368 |
| positive regulation of ossification | GO:0045778 | under | 0.0375 |
| positive regulation of osteoblast differentiation | GO:0045669 | under | 0.0375 |
| regulation of body fluid levels | GO:0050878 | under | 0.0375 |
| adenylate cyclase-activating G-protein coupled receptor signaling pathway | GO:0007189 | over | 0.0383 |
| adenylate cyclase-modulating G-protein coupled receptor signaling pathway | GO:0007188 | over | 0.0383 |
| G-protein coupled receptor signaling pathway, coupled to cyclic nucleotide second messenger | GO:0007187 | over | 0.0383 |
| non-canonical Wnt signaling pathway | GO:0035567 | under | 0.0383 |
| response to acid chemical | GO:0001101 | under | 0.0386 |
| positive regulation of lipid transport | GO:0032370 | over | 0.0388 |
| odontogenesis | GO:0042476 | under | 0.0394 |
| odontogenesis of dentin-containing tooth | GO:0042475 | under | 0.0394 |
| amelogenesis | GO:0097186 | under | 0.0394 |
| positive regulation of defense response | GO:0031349 | under | 0.0398 |
| animal organ development | GO:0048513 | under | 0.0411 |
| cellular divalent inorganic cation homeostasis | GO:0072503 | under | 0.0414 |
| somatic hypermutation of immunoglobulin genes | GO:0016446 | over | 0.0416 |
| somatic diversification of immune receptors via somatic mutation | GO:0002566 | over | 0.0416 |
| fatty acid metabolic process | GO:0006631 | under | 0.0417 |
| protein polymerization | GO:0051258 | under | 0.0433 |
| inositol lipid-mediated signaling | GO:0048017 | over | 0.0438 |
| phosphatidylinositol-mediated signaling | GO:0048015 | over | 0.0438 |
| oligopeptide transport | GO:0006857 | under | 0.0438 |
| peptidyl-lysine deacetylation | GO:0034983 | over | 0.0438 |
| negative regulation of peptidyl-lysine acetylation | GO:2000757 | over | 0.0438 |
| regulation of peptidyl-lysine acetylation | GO:2000756 | over | 0.0438 |
| negative regulation of protein acetylation | GO:1901984 | over | 0.0438 |
| regulation of muscle contraction | GO:0006937 | under | 0.0443 |
| vasculature development | GO:0001944 | under | 0.0447 |
| blood vessel development | GO:0001568 | under | 0.0447 |
| negative regulation of response to biotic stimulus | GO:0002832 | over | 0.045 |
| positive regulation of response to endoplasmic reticulum stress | GO:1905898 | over | 0.0462 |
| cellular response to light stimulus | GO:0071482 | over | 0.0464 |
| cellular response to UV | GO:0034644 | over | 0.0464 |
| drug transport | GO:0015893 | under | 0.047 |
| apoptotic cell clearance | GO:0043277 | under | 0.0478 |
| ion transport | GO:0006811 | under | 0.0478 |
| leukocyte homeostasis | GO:0001776 | under | 0.0478 |
| hexose transport | GO:0008645 | over | 0.0488 |
| glucose transport | GO:0015758 | over | 0.0488 |
| monosaccharide transport | GO:0015749 | over | 0.0488 |
| protein deacylation | GO:0035601 | over | 0.0491 |
| macromolecule deacylation | GO:0098732 | over | 0.0491 |
| I-kappaB kinase/NF-kappaB signaling | GO:0007249 | over | 0.0493 |
| regulation of membrane potential | GO:0042391 | under | 0.05 |

| GO Term Pathway: NesCreCsf1flfl_Cb_9w | GO Term_ID | overUnder | pvalue |
|---|---|---|---|
| response to external biotic stimulus | GO:0043207 | under | 0.000578 |
| response to other organism | GO:0051707 | under | 0.000578 |
| response to biotic stimulus | GO:0009607 | under | 0.000578 |
| regulation of heart contraction | GO:0008016 | under | 0.00233 |
| multicellular organismal signaling | GO:0035637 | under | 0.00662 |
| AV node cell to bundle of His cell communication | GO:0086067 | under | 0.00662 |
| cell communication involved in cardiac conduction | GO:0086065 | under | 0.00662 |
| cardiac conduction | GO:0061337 | under | 0.00662 |
| defense response to other organism | GO:0098542 | under | 0.0073 |
| defense response | GO:0006952 | under | 0.00761 |
| DNA replication | GO:0006260 | under | 0.00971 |
| response to virus | GO:0009615 | under | 0.00985 |
| innate immune response | GO:0045087 | under | 0.01 |
| in utero embryonic development | GO:0001701 | under | 0.0102 |
| regulation of transmembrane receptor protein serine/threonine kinase signaling pathway | GO:0090092 | under | 0.0102 |
| chordate embryonic development | GO:0043009 | under | 0.0107 |
| embryo development ending in birth or egg hatching | GO:0009792 | under | 0.0107 |
| multi-organism process | GO:0051704 | under | 0.0123 |
| positive regulation of reactive oxygen species metabolic process | GO:2000379 | under | 0.0123 |
| positive regulation of DNA binding transcription factor activity | GO:0051091 | under | 0.0136 |
| gap junction assembly | GO:0016264 | under | 0.0152 |
| cell communication by electrical coupling involved in cardiac conduction | GO:0086064 | under | 0.0152 |
| AV node cell to bundle of His cell communication by electrical coupling | GO:0086053 | under | 0.0152 |
| cell communication by electrical coupling | GO:0010644 | under | 0.0152 |
| response to bacterium | GO:0009617 | under | 0.0161 |
| regulation of phosphatidylinositol 3-kinase signaling | GO:0014066 | over | 0.0165 |
| proteoglycan biosynthetic process | GO:0030166 | over | 0.0194 |
| regulation of cellular response to growth factor stimulus | GO:0090287 | under | 0.0206 |
| cell development | GO:0048468 | under | 0.0216 |
| cellular response to lipid | GO:0071396 | under | 0.0217 |
| DNA duplex unwinding | GO:0032508 | under | 0.0228 |
| DNA geometric change | GO:0032392 | under | 0.0228 |
| regulation of phospholipid biosynthetic process | GO:0071071 | over | 0.0232 |
| movement of cell or subcellular component | GO:0006928 | under | 0.0239 |
| response to ethanol | GO:0045471 | over | 0.0249 |
| dicarboxylic acid metabolic process | GO:0043648 | over | 0.0255 |
| negative regulation of mitochondrion organization | GO:0010823 | over | 0.0259 |
| negative regulation of release of cytochrome c from mitochondria | GO:0090201 | over | 0.0259 |
| regulation of release of cytochrome c from mitochondria | GO:0090199 | over | 0.0259 |
| cell differentiation | GO:0030154 | under | 0.0263 |
| response to lipid | GO:0033993 | under | 0.0264 |
| double-strand break repair | GO:0006302 | under | 0.0273 |
| ribonucleoprotein complex subunit organization | GO:0071826 | over | 0.0275 |
| ribonucleoprotein complex assembly | GO:0022618 | over | 0.0275 |
| meiosis I | GO:0007127 | under | 0.0275 |
| meiotic nuclear division | GO:0140013 | under | 0.0275 |
| organonitrogen compound metabolic process | GO:1901564 | under | 0.0276 |
| regulation of blood circulation | GO:1903522 | under | 0.0276 |
| cellular response to drug | GO:0035690 | under | 0.0277 |
| peptidyl-amino acid modification | GO:0018193 | under | 0.0278 |
| response to stimulus | GO:0050896 | under | 0.0278 |
| sensory organ morphogenesis | GO:0090596 | under | 0.0287 |
| regulation of phospholipase activity | GO:0010517 | under | 0.0289 |
| regulation of intracellular steroid hormone receptor signaling pathway | GO:0033143 | over | 0.0293 |
| regulation of toll-like receptor signaling pathway | GO:0034121 | under | 0.0296 |
| regulation of DNA binding transcription factor activity | GO:0051090 | under | 0.0304 |
| positive regulation of gene expression | GO:0010628 | under | 0.0313 |
| positive regulation of transmembrane receptor protein serine/threonine kinase signaling pathway | GO:0090100 | under | 0.0316 |
| positive regulation of cell differentiation | GO:0045597 | under | 0.0321 |
| negative regulation of ERK1 and ERK2 cascade | GO:0070373 | over | 0.0321 |
| tRNA processing | GO:0008033 | over | 0.0326 |
| response to heat | GO:0009408 | over | 0.0327 |
| response to temperature stimulus | GO:0009266 | over | 0.0327 |
| positive regulation of protein complex assembly | GO:0031334 | under | 0.0328 |
| peptidyl-threonine modification | GO:0018210 | under | 0.033 |
| peptide cross-linking | GO:0018149 | under | 0.0331 |
| negative regulation of response to biotic stimulus | GO:0002832 | under | 0.0335 |
| RNA modification | GO:0009451 | over | 0.0337 |
| regulation of vesicle fusion | GO:0031338 | under | 0.0341 |
| positive regulation of reactive oxygen species biosynthetic process | GO:1903428 | under | 0.0346 |
| positive regulation of nitric oxide biosynthetic process | GO:0045429 | under | 0.0346 |
| positive regulation of nitric oxide metabolic process | GO:1904407 | under | 0.0346 |
| positive regulation of mitotic cell cycle phase transition | GO:1901992 | under | 0.035 |
| positive regulation of cell cycle phase transition | GO:1901989 | under | 0.035 |
| positive regulation of phospholipid biosynthetic process | GO:0071073 | over | 0.0352 |
| cellular response to steroid hormone stimulus | GO:0071383 | under | 0.0365 |
| response to steroid hormone | GO:0048545 | under | 0.0365 |
| RNA processing | GO:0006396 | over | 0.0366 |
| homeostasis of number of cells | GO:0048872 | under | 0.0374 |

| | | | |
|---|---|---|---|
| ncRNA processing | GO:0034470 | over | 0.0377 |
| activation of protein kinase activity | GO:0032147 | over | 0.0397 |
| mammary gland epithelium development | GO:0061180 | under | 0.0399 |
| positive regulation of cellular amide metabolic process | GO:0034250 | under | 0.0399 |
| positive regulation of translation | GO:0045727 | under | 0.0399 |
| cellular developmental process | GO:0048869 | under | 0.0403 |
| dephosphorylation | GO:0016311 | under | 0.0415 |
| positive regulation of NF-kappaB transcription factor activity | GO:0051092 | under | 0.0419 |
| regulation of BMP signaling pathway | GO:0030510 | under | 0.0426 |
| carbohydrate homeostasis | GO:0033500 | over | 0.0431 |
| glucose homeostasis | GO:0042593 | over | 0.0431 |
| activation of MAPK activity | GO:0000187 | over | 0.0432 |
| sensory perception of light stimulus | GO:0050953 | under | 0.044 |
| visual perception | GO:0007601 | under | 0.044 |
| recombinational repair | GO:0000725 | under | 0.0442 |
| double-strand break repair via homologous recombination | GO:0000724 | under | 0.0442 |
| peptidyl-serine modification | GO:0018209 | under | 0.0442 |
| protein modification process | GO:0036211 | under | 0.0453 |
| cellular protein modification process | GO:0006464 | under | 0.0453 |
| stem cell differentiation | GO:0048863 | under | 0.0458 |
| aminoglycan catabolic process | GO:0006026 | under | 0.0468 |
| animal organ development | GO:0048513 | under | 0.047 |
| regulation of ERBB signaling pathway | GO:1901184 | under | 0.0471 |
| regulation of epidermal growth factor receptor signaling pathway | GO:0042058 | under | 0.0471 |
| negative regulation of immune effector process | GO:0002698 | under | 0.0472 |
| cellular response to glucocorticoid stimulus | GO:0071385 | under | 0.0482 |
| cellular response to corticosteroid stimulus | GO:0071384 | under | 0.0482 |
| response to corticosteroid | GO:0031960 | under | 0.0482 |
| response to glucocorticoid | GO:0051384 | under | 0.0482 |
| negative regulation of MAPK cascade | GO:0043409 | over | 0.0482 |
| response to stress | GO:0006950 | under | 0.0494 |
| cellular response to organic substance | GO:0071310 | under | 0.0499 |
| membrane lipid metabolic process | GO:0006643 | under | 0.0499 |

Supplementary Table S2, cont.

| GO Term Pathway: NesCreCsf1flfl_Fb_9w | GO Term_ID | overUnder | pvalue |
|---|---|---|---|
| oxoacid metabolic process | GO:0043436 | over | 0.000899 |
| organic acid metabolic process | GO:0006082 | over | 0.000899 |
| regulation of reactive oxygen species metabolic process | GO:2000377 | under | 0.00179 |
| ATP metabolic process | GO:0046034 | over | 0.00287 |
| carboxylic acid metabolic process | GO:0019752 | over | 0.00445 |
| respiratory system development | GO:0060541 | under | 0.00468 |
| ribonucleoside triphosphate metabolic process | GO:0009199 | over | 0.00488 |
| purine ribonucleoside triphosphate metabolic process | GO:0009205 | over | 0.00488 |
| positive regulation of reactive oxygen species metabolic process | GO:2000379 | under | 0.00537 |
| regulation of neurotransmitter levels | GO:0001505 | under | 0.00608 |
| regulation of reactive oxygen species biosynthetic process | GO:1903426 | under | 0.00838 |
| lung development | GO:0030324 | under | 0.0095 |
| respiratory tube development | GO:0030323 | under | 0.0095 |
| response to heat | GO:0009408 | under | 0.011 |
| cellular amino acid metabolic process | GO:0006520 | over | 0.0113 |
| rRNA metabolic process | GO:0016072 | over | 0.0115 |
| mesenchymal cell development | GO:0014031 | under | 0.0133 |
| mesenchyme development | GO:0060485 | under | 0.0133 |
| mesenchymal cell differentiation | GO:0048762 | under | 0.0133 |
| ribosome biogenesis | GO:0042254 | over | 0.0145 |
| bone remodeling | GO:0046849 | over | 0.0162 |
| positive regulation of nitric oxide metabolic process | GO:1904407 | under | 0.0172 |
| positive regulation of nitric oxide biosynthetic process | GO:0045429 | under | 0.0172 |
| regulation of nitric oxide biosynthetic process | GO:0045428 | under | 0.0172 |
| positive regulation of reactive oxygen species biosynthetic process | GO:1903428 | under | 0.0172 |
| positive regulation of developmental process | GO:0051094 | under | 0.0182 |
| neural crest cell differentiation | GO:0014033 | under | 0.0191 |
| neural crest cell development | GO:0014032 | under | 0.0191 |
| stem cell development | GO:0048864 | under | 0.0191 |
| stem cell differentiation | GO:0048863 | under | 0.0191 |
| rRNA processing | GO:0006364 | over | 0.0191 |
| positive regulation of proteolysis | GO:0045862 | under | 0.0195 |
| positive regulation of cell differentiation | GO:0045597 | under | 0.0198 |
| positive regulation of leukocyte chemotaxis | GO:0002690 | over | 0.0201 |
| regulation of leukocyte chemotaxis | GO:0002688 | over | 0.0201 |
| regulation of mononuclear cell migration | GO:0071675 | over | 0.0201 |
| positive regulation of chemotaxis | GO:0050921 | over | 0.0201 |
| regulation of chemotaxis | GO:0050920 | over | 0.0201 |
| alpha-amino acid metabolic process | GO:1901605 | over | 0.0208 |
| aspartate family amino acid metabolic process | GO:0009066 | over | 0.0215 |
| protein prenylation | GO:0018342 | over | 0.0217 |
| prenylation | GO:0097354 | over | 0.0217 |
| corticospinal tract morphogenesis | GO:0021957 | under | 0.0227 |
| central nervous system neuron axonogenesis | GO:0021955 | under | 0.0227 |
| central nervous system neuron development | GO:0021954 | under | 0.0227 |
| central nervous system projection neuron axonogenesis | GO:0021952 | under | 0.0227 |
| regulation of T cell differentiation in thymus | GO:0033081 | under | 0.0229 |
| generation of precursor metabolites and energy | GO:0006091 | over | 0.0246 |
| monocarboxylic acid metabolic process | GO:0032787 | over | 0.0251 |
| odontogenesis | GO:0042476 | under | 0.0258 |
| programmed cell death | GO:0012501 | under | 0.0262 |
| regulation of cell differentiation | GO:0045595 | under | 0.0268 |
| aspartate metabolic process | GO:0006531 | over | 0.027 |
| nicotinamide nucleotide biosynthetic process from aspartate | GO:0019355 | over | 0.027 |
| 'de novo' NAD biosynthetic process from aspartate | GO:0034628 | over | 0.027 |
| 'de novo' NAD biosynthetic process | GO:0034627 | over | 0.027 |
| regulation of cysteine-type endopeptidase activity involved in apoptotic process | GO:0043281 | under | 0.027 |
| DNA replication | GO:0006260 | over | 0.0272 |
| regulation of endocytosis | GO:0030100 | under | 0.0282 |
| NAD biosynthetic process | GO:0009435 | over | 0.0288 |
| purine ribonucleoside monophosphate metabolic process | GO:0009167 | over | 0.0288 |
| ribonucleoside monophosphate metabolic process | GO:0009161 | over | 0.0288 |
| purine nucleoside monophosphate metabolic process | GO:0009126 | over | 0.0288 |
| nucleoside monophosphate metabolic process | GO:0009123 | over | 0.0288 |
| purine ribonucleotide metabolic process | GO:0009150 | over | 0.0288 |

| | | | |
|---|---|---|---|
| ribonucleotide metabolic process | GO:0009259 | over | 0.0288 |
| ribose phosphate metabolic process | GO:0019693 | over | 0.0288 |
| developmental growth involved in morphogenesis | GO:0060560 | under | 0.0306 |
| developmental cell growth | GO:0048588 | under | 0.0306 |
| cell death | GO:0008219 | under | 0.0306 |
| purine nucleoside triphosphate metabolic process | GO:0009144 | over | 0.0316 |
| nucleoside triphosphate metabolic process | GO:0009141 | over | 0.0316 |
| striated muscle tissue development | GO:0014706 | over | 0.0327 |
| muscle tissue development | GO:0060537 | over | 0.0327 |
| cellular component morphogenesis | GO:0032989 | under | 0.0332 |
| integrin-mediated signaling pathway | GO:0007229 | under | 0.0334 |
| cell growth | GO:0016049 | under | 0.0338 |
| humoral immune response | GO:0006959 | over | 0.0339 |
| positive regulation of cell cycle checkpoint | GO:1901978 | over | 0.0344 |
| regulation of cell cycle checkpoint | GO:1901976 | over | 0.0344 |
| apoptotic process | GO:0006915 | under | 0.035 |
| cell morphogenesis | GO:0000902 | under | 0.0369 |
| negative regulation of inflammatory response | GO:0050728 | under | 0.0379 |
| regulation of neuron death | GO:1901214 | under | 0.0382 |
| positive regulation of cell death | GO:0010942 | under | 0.0401 |
| cell differentiation | GO:0030154 | under | 0.0404 |
| positive regulation of programmed cell death | GO:0043068 | under | 0.0404 |
| positive regulation of apoptotic process | GO:0043065 | under | 0.0404 |
| regulation of cell morphogenesis involved in differentiation | GO:0010769 | under | 0.0411 |
| positive regulation of T cell differentiation in thymus | GO:0033089 | under | 0.0411 |
| neurotransmitter transport | GO:0006836 | under | 0.0413 |
| tissue remodeling | GO:0048771 | over | 0.0418 |
| regulation of multicellular organismal development | GO:2000026 | under | 0.0431 |
| cellular developmental process | GO:0048869 | under | 0.0435 |
| coenzyme metabolic process | GO:0006732 | over | 0.045 |
| B cell activation | GO:0042113 | under | 0.0451 |
| protein complex localization | GO:0031503 | over | 0.0455 |
| response to nutrient | GO:0007584 | under | 0.0465 |
| positive regulation of neuron death | GO:1901216 | under | 0.0469 |
| negative regulation of kinase activity | GO:0033673 | under | 0.0471 |
| placenta development | GO:0001890 | under | 0.0471 |
| cell part morphogenesis | GO:0032990 | under | 0.0472 |

Supplementary Table S2, cont.

| GO Term Pathway: Il34LaczLacz_Cb_9w | GO Term_ID | overUnder | pvalue |
|---|---|---|---|
| regulation of multicellular organismal process | GO:0051239 | over | 0.00274 |
| regulation of leukocyte cell-cell adhesion | GO:1903037 | over | 0.00303 |
| regulation of cell-cell adhesion | GO:0022407 | over | 0.00303 |
| regulation of T cell activation | GO:0050863 | over | 0.00303 |
| regulation of cell adhesion | GO:0030155 | over | 0.00458 |
| regulation of T cell proliferation | GO:0042129 | over | 0.0051 |
| regulation of actin polymerization or depolymerization | GO:0008064 | over | 0.00625 |
| regulation of cellular component size | GO:0032535 | over | 0.00625 |
| regulation of actin filament polymerization | GO:0030833 | over | 0.00625 |
| regulation of actin filament length | GO:0030832 | over | 0.00625 |
| regulation of anatomical structure size | GO:0090066 | over | 0.00625 |
| regulation of protein polymerization | GO:0032271 | over | 0.00625 |
| monovalent inorganic cation transport | GO:0015672 | under | 0.00627 |
| regulation of actin filament organization | GO:0110053 | over | 0.00708 |
| regulation of actin filament-based process | GO:0032970 | over | 0.00708 |
| regulation of actin cytoskeleton organization | GO:0032956 | over | 0.00708 |
| metal ion transport | GO:0030001 | under | 0.0073 |
| regulation of lymphocyte proliferation | GO:0050670 | over | 0.00847 |
| regulation of leukocyte proliferation | GO:0070663 | over | 0.00847 |
| regulation of mononuclear cell proliferation | GO:0032944 | over | 0.00847 |
| regulation of lymphocyte activation | GO:0051249 | over | 0.00906 |
| vesicle docking involved in exocytosis | GO:0006904 | under | 0.00923 |
| response to mechanical stimulus | GO:0009612 | under | 0.0103 |
| trans-synaptic signaling | GO:0099537 | under | 0.0104 |
| synaptic signaling | GO:0099536 | under | 0.0104 |
| chemical synaptic transmission | GO:0007268 | under | 0.0104 |
| anterograde trans-synaptic signaling | GO:0098916 | under | 0.0104 |
| regulation of cell morphogenesis | GO:0022604 | over | 0.0104 |
| regulation of cytoskeleton organization | GO:0051493 | over | 0.0109 |
| regulation of cell differentiation | GO:0045595 | over | 0.0125 |
| regulation of leukocyte activation | GO:0002694 | over | 0.0128 |
| regulation of cell activation | GO:0050865 | over | 0.0128 |
| multicellular organismal homeostasis | GO:0048871 | over | 0.0138 |
| tissue homeostasis | GO:0001894 | over | 0.0138 |
| regulation of cell morphogenesis involved in differentiation | GO:0010769 | over | 0.0161 |
| exocytic process | GO:0140029 | under | 0.0165 |
| positive regulation of ion transport | GO:0043270 | over | 0.0176 |
| microtubule-based process | GO:0007017 | under | 0.0183 |
| regulation of immune system process | GO:0002682 | over | 0.0206 |
| organophosphate ester transport | GO:0015748 | over | 0.021 |
| phospholipid transport | GO:0015914 | over | 0.021 |
| regulation of anion transport | GO:0044070 | over | 0.0216 |
| organic cation transport | GO:0015695 | over | 0.0216 |
| positive regulation of ion transmembrane transport | GO:0034767 | over | 0.0216 |
| positive regulation of transmembrane transport | GO:0034764 | over | 0.0216 |
| positive regulation of ion transmembrane transporter activity | GO:0032414 | over | 0.0216 |
| positive regulation of transporter activity | GO:0032411 | over | 0.0216 |
| positive regulation of actin filament polymerization | GO:0030838 | over | 0.0241 |
| positive regulation of cytoskeleton organization | GO:0051495 | over | 0.0241 |
| positive regulation of protein polymerization | GO:0032273 | over | 0.0241 |
| positive regulation of regulated secretory pathway | GO:1903307 | under | 0.0251 |
| cellular glucose homeostasis | GO:0001678 | under | 0.0251 |
| response to monosaccharide | GO:0034284 | under | 0.0251 |
| response to glucose | GO:0009749 | under | 0.0251 |
| response to hexose | GO:0009746 | under | 0.0251 |
| response to carbohydrate | GO:0009743 | under | 0.0251 |
| cellular response to glucose stimulus | GO:0071333 | under | 0.0251 |
| cellular response to hexose stimulus | GO:0071331 | under | 0.0251 |
| cellular response to monosaccharide stimulus | GO:0071326 | under | 0.0251 |
| cellular response to carbohydrate stimulus | GO:0071322 | under | 0.0251 |
| positive regulation of calcium ion-dependent exocytosis | GO:0045956 | under | 0.0251 |
| action potential | GO:0001508 | under | 0.0258 |
| organic anion transport | GO:0015711 | over | 0.026 |
| regulation of system process | GO:0044057 | over | 0.0271 |
| Ras protein signal transduction | GO:0007265 | over | 0.0273 |
| immune response | GO:0006955 | over | 0.0274 |
| positive regulation of cell morphogenesis involved in differentiation | GO:0010770 | over | 0.0285 |
| regulation of calcium ion-dependent exocytosis | GO:0017158 | under | 0.029 |
| regulation of cell cycle | GO:0051726 | under | 0.0297 |
| sodium ion transport | GO:0006814 | under | 0.0299 |
| cellular chemical homeostasis | GO:0055082 | under | 0.0329 |

| | | | |
|---|---|---|---|
| blood circulation | GO:0008015 | under | 0.033 |
| circulatory system process | GO:0003013 | under | 0.033 |
| exocytosis | GO:0006887 | under | 0.0338 |
| regulation of nuclease activity | GO:0032069 | under | 0.0338 |
| regulation of proteasomal protein catabolic process | GO:0061136 | under | 0.0339 |
| positive regulation of cell development | GO:0010720 | over | 0.0343 |
| regulation of cell development | GO:0060284 | over | 0.0343 |
| drug transport | GO:0015893 | over | 0.0347 |
| positive regulation of peptidyl-tyrosine phosphorylation | GO:0050731 | over | 0.0347 |
| drug transmembrane transport | GO:0006855 | over | 0.0347 |
| response to interferon-gamma | GO:0034341 | over | 0.0349 |
| synapse organization | GO:0050808 | under | 0.0357 |
| immune system process | GO:0002376 | over | 0.0364 |
| regulation of supramolecular fiber organization | GO:1902903 | over | 0.0365 |
| regulation of anatomical structure morphogenesis | GO:0022603 | over | 0.0372 |
| antigen processing and presentation of peptide or polysaccharide antigen via MHC class II | GO:0002504 | over | 0.0375 |
| antigen processing and presentation of peptide antigen via MHC class II | GO:0002495 | over | 0.0375 |
| antigen processing and presentation of exogenous peptide antigen | GO:0002478 | over | 0.0375 |
| antigen processing and presentation of exogenous peptide antigen via MHC class II | GO:0019886 | over | 0.0375 |
| antigen processing and presentation of exogenous antigen | GO:0019884 | over | 0.0375 |
| antigen processing and presentation of peptide antigen | GO:0048002 | over | 0.0375 |
| protein localization to cilium | GO:0061512 | under | 0.0375 |
| synaptic vesicle localization | GO:0097479 | under | 0.0382 |
| signal release from synapse | GO:0099643 | under | 0.0382 |
| vesicle localization | GO:0051648 | under | 0.0382 |
| presynaptic process involved in chemical synaptic transmission | GO:0099531 | under | 0.0382 |
| synaptic vesicle cycle | GO:0099504 | under | 0.0382 |
| neurotransmitter secretion | GO:0007269 | under | 0.0382 |
| cell cycle | GO:0007049 | under | 0.0386 |
| positive regulation of apoptotic signaling pathway | GO:2001235 | over | 0.0398 |
| positive regulation of cell adhesion | GO:0045785 | over | 0.0404 |
| regulation of developmental process | GO:0050793 | over | 0.042 |
| positive regulation of cellular component movement | GO:0051272 | over | 0.0434 |
| positive regulation of cell migration | GO:0030335 | over | 0.0434 |
| positive regulation of cell motility | GO:2000147 | over | 0.0434 |
| drug catabolic process | GO:0042737 | over | 0.044 |
| regulation of viral entry into host cell | GO:0046596 | over | 0.045 |
| limb morphogenesis | GO:0035108 | over | 0.0471 |
| appendage morphogenesis | GO:0035107 | over | 0.0471 |
| appendage development | GO:0048736 | over | 0.0471 |
| limb development | GO:0060173 | over | 0.0471 |
| microtubule cytoskeleton organization | GO:0000226 | under | 0.0477 |
| negative regulation of immune system process | GO:0002683 | over | 0.048 |
| regulation of cAMP metabolic process | GO:0030814 | under | 0.0482 |
| regulation of cyclic nucleotide metabolic process | GO:0030799 | under | 0.0482 |
| positive regulation of cell differentiation | GO:0045597 | over | 0.0494 |

| GO Term Pathway: Il34LaczLacz_Fb_9w | GO Term_ID | overUnder | pvalue |
|---|---|---|---|
| protein phosphorylation | GO:0006468 | under | 0.00354 |
| lymphocyte chemotaxis | GO:0048247 | over | 0.00501 |
| glycerolipid catabolic process | GO:0046503 | over | 0.00655 |
| negative regulation of supramolecular fiber organization | GO:1902904 | over | 0.00673 |
| negative regulation of cytoskeleton organization | GO:0051494 | over | 0.00673 |
| negative regulation of toll-like receptor signaling pathway | GO:0034122 | under | 0.00693 |
| negative regulation of protein complex disassembly | GO:0043242 | over | 0.0071 |
| negative regulation of protein depolymerization | GO:1901880 | over | 0.0071 |
| regulation of protein depolymerization | GO:1901879 | over | 0.0071 |
| macromolecule modification | GO:0043412 | under | 0.0075 |
| cellular protein metabolic process | GO:0044267 | under | 0.00754 |
| cellular lipid catabolic process | GO:0044242 | over | 0.00759 |
| cytokine-mediated signaling pathway | GO:0019221 | over | 0.00837 |
| mononuclear cell migration | GO:0071674 | over | 0.00867 |
| monocyte chemotaxis | GO:0002548 | over | 0.00867 |
| lymphocyte migration | GO:0072676 | over | 0.00879 |
| cyclic nucleotide metabolic process | GO:0009187 | under | 0.00881 |
| cAMP metabolic process | GO:0046058 | under | 0.00881 |
| positive regulation of cytokine-mediated signaling pathway | GO:0001961 | over | 0.00889 |
| positive regulation of response to cytokine stimulus | GO:0060760 | over | 0.00889 |
| potassium ion import | GO:0010107 | over | 0.00986 |
| ensheathment of neurons | GO:0007272 | over | 0.01 |
| myelination | GO:0042552 | over | 0.01 |
| axon ensheathment | GO:0008366 | over | 0.01 |
| regulation of peptide hormone secretion | GO:0090276 | under | 0.0113 |
| regulation of insulin secretion | GO:0050796 | under | 0.0113 |
| protein metabolic process | GO:0019538 | under | 0.0126 |
| negative regulation of intrinsic apoptotic signaling pathway | GO:2001243 | over | 0.0138 |
| organonitrogen compound metabolic process | GO:1901564 | under | 0.0144 |
| protein modification process | GO:0036211 | under | 0.0149 |
| cellular protein modification process | GO:0006464 | under | 0.0149 |
| phosphorylation | GO:0016310 | under | 0.0157 |
| ribonucleotide biosynthetic process | GO:0009260 | under | 0.0174 |
| ribose phosphate biosynthetic process | GO:0046390 | under | 0.0174 |
| double-strand break repair | GO:0006302 | under | 0.0185 |
| ribonucleotide metabolic process | GO:0009259 | under | 0.0197 |
| ribose phosphate metabolic process | GO:0019693 | under | 0.0197 |
| peptidyl-threonine modification | GO:0018210 | under | 0.0206 |
| negative regulation of immune response | GO:0050777 | under | 0.0206 |
| positive regulation of cytokine production | GO:0001819 | over | 0.0211 |
| developmental growth involved in morphogenesis | GO:0060560 | over | 0.0212 |
| mitochondrial gene expression | GO:0140053 | under | 0.0216 |
| recombinational repair | GO:0000725 | under | 0.0221 |
| double-strand break repair via homologous recombination | GO:0000724 | under | 0.0221 |
| positive regulation of membrane protein ectodomain proteolysis | GO:0051044 | over | 0.0227 |
| regulation of membrane protein ectodomain proteolysis | GO:0051043 | over | 0.0227 |
| protein complex disassembly | GO:0043241 | over | 0.0244 |
| tRNA processing | GO:0008033 | under | 0.0245 |
| 'de novo' protein folding | GO:0006458 | over | 0.0246 |
| 'de novo' posttranslational protein folding | GO:0051084 | over | 0.0246 |
| tRNA modification | GO:0006400 | under | 0.0248 |
| tRNA methylation | GO:0030488 | under | 0.0248 |
| double-strand break repair via nonhomologous end joining | GO:0006303 | under | 0.0256 |
| non-recombinational repair | GO:0000726 | under | 0.0256 |
| translational elongation | GO:0006414 | under | 0.0256 |
| negative regulation of intrinsic apoptotic signaling pathway by p53 class mediator | GO:1902254 | over | 0.0259 |
| regulation of intrinsic apoptotic signaling pathway by p53 class mediator | GO:1902253 | over | 0.0259 |
| negative regulation of signal transduction by p53 class mediator | GO:1901797 | over | 0.0259 |
| gastrulation | GO:0007369 | under | 0.0263 |
| regulation of plasma membrane bounded cell projection assembly | GO:0120032 | under | 0.0268 |
| regulation of cell projection assembly | GO:0060491 | under | 0.0268 |
| regulation of anion transmembrane transport | GO:1903959 | under | 0.0273 |
| regulation of anion channel activity | GO:0010359 | under | 0.0273 |
| negative regulation of amyloid precursor protein catabolic process | GO:1902992 | over | 0.0274 |
| regulation of amyloid precursor protein catabolic process | GO:1902991 | over | 0.0277 |

| | | | |
|---|---|---|---|
| peptidyl-serine phosphorylation | GO:0018105 | under | 0.0282 |
| peptidyl-threonine phosphorylation | GO:0018107 | under | 0.0288 |
| nucleotide-excision repair | GO:0006289 | under | 0.0293 |
| negative regulation of type I interferon production | GO:0032480 | under | 0.0294 |
| negative regulation of lipopolysaccharide-mediated signaling pathway | GO:0031665 | under | 0.0294 |
| cellular response to toxic substance | GO:0097237 | over | 0.0295 |
| peptidyl-amino acid modification | GO:0018193 | under | 0.0296 |
| negative regulation of organelle organization | GO:0010639 | over | 0.0297 |
| multi-multicellular organism process | GO:0044706 | under | 0.0302 |
| apoptotic nuclear changes | GO:0030262 | under | 0.0303 |
| cellular component disassembly involved in execution phase of apoptosis | GO:0006921 | under | 0.0303 |
| formation of primary germ layer | GO:0001704 | under | 0.0314 |
| receptor catabolic process | GO:0032801 | under | 0.0317 |
| histone H4 acetylation | GO:0043967 | under | 0.0319 |
| regulation of protein kinase A signaling | GO:0010738 | under | 0.033 |
| neural tube closure | GO:0001843 | over | 0.0331 |
| primary neural tube formation | GO:0014020 | over | 0.0331 |
| tube closure | GO:0060606 | over | 0.0331 |
| response to hydrogen peroxide | GO:0042542 | over | 0.0333 |
| cellular response to hydrogen peroxide | GO:0070301 | over | 0.0333 |
| histone H3 acetylation | GO:0043966 | under | 0.0334 |
| triglyceride metabolic process | GO:0006641 | over | 0.0336 |
| acylglycerol metabolic process | GO:0006639 | over | 0.0336 |
| neutral lipid metabolic process | GO:0006638 | over | 0.0336 |
| acylglycerol homeostasis | GO:0055090 | over | 0.0336 |
| acylglycerol catabolic process | GO:0046464 | over | 0.0336 |
| neutral lipid catabolic process | GO:0046461 | over | 0.0336 |
| very-low-density lipoprotein particle remodeling | GO:0034372 | over | 0.0336 |
| triglyceride-rich lipoprotein particle remodeling | GO:0034370 | over | 0.0336 |
| triglyceride homeostasis | GO:0070328 | over | 0.0336 |
| triglyceride catabolic process | GO:0019433 | over | 0.0336 |
| positive regulation of monocyte differentiation | GO:0045657 | over | 0.0343 |
| regulation of monocyte differentiation | GO:0045655 | over | 0.0343 |
| chaperone-mediated protein folding | GO:0061077 | over | 0.0343 |
| regulation of lipopolysaccharide-mediated signaling pathway | GO:0031664 | under | 0.0345 |
| macrophage migration | GO:1905517 | over | 0.0346 |
| astrocyte cell migration | GO:0043615 | over | 0.0346 |
| macrophage chemotaxis | GO:0048246 | over | 0.0346 |
| apoptotic process | GO:0006915 | under | 0.0348 |
| positive regulation of mitotic cell cycle | GO:0045931 | under | 0.0354 |
| regulation of toll-like receptor signaling pathway | GO:0034121 | under | 0.0355 |
| regulation of leukocyte proliferation | GO:0070663 | over | 0.0356 |
| regulation of microtubule depolymerization | GO:0031114 | over | 0.0356 |
| negative regulation of microtubule polymerization or depolymerization | GO:0031111 | over | 0.0356 |
| negative regulation of microtubule depolymerization | GO:0007026 | over | 0.0356 |
| regulation of supramolecular fiber organization | GO:1902903 | over | 0.0357 |
| cardiac muscle cell differentiation | GO:0055007 | under | 0.0359 |
| cardiocyte differentiation | GO:0035051 | under | 0.0359 |
| regulation of intrinsic apoptotic signaling pathway | GO:2001242 | over | 0.0363 |
| protein acylation | GO:0043543 | under | 0.0364 |
| histone acetylation | GO:0016573 | under | 0.0369 |
| peptidyl-lysine acetylation | GO:0018394 | under | 0.0369 |
| internal peptidyl-lysine acetylation | GO:0018393 | under | 0.0369 |
| negative regulation of interleukin-8 production | GO:0032717 | under | 0.037 |
| regulation of DNA-dependent DNA replication | GO:0090329 | under | 0.037 |
| Golgi vesicle budding | GO:0048194 | under | 0.0378 |
| vesicle budding from membrane | GO:0006900 | under | 0.0378 |
| regulation of GTPase activity | GO:0043087 | over | 0.0378 |
| regulation of natural killer cell chemotaxis | GO:2000501 | over | 0.0384 |
| positive regulation of macrophage derived foam cell differentiation | GO:0010744 | over | 0.0384 |
| programmed cell death | GO:0012501 | under | 0.0385 |
| renal system process | GO:0003014 | over | 0.0388 |
| regeneration | GO:0031099 | over | 0.0391 |
| protein lipidation | GO:0006497 | under | 0.0394 |
| positive regulation of gliogenesis | GO:0014015 | over | 0.0394 |
| smoothened signaling pathway | GO:0007224 | over | 0.0394 |

| | | | |
|---|---|---|---|
| macromolecule metabolic process | GO:0043170 | under | 0.0395 |
| adenylate cyclase-activating G-protein coupled receptor signaling pathway | GO:0007189 | under | 0.04 |
| cAMP-mediated signaling | GO:0019933 | under | 0.04 |
| regulation of cytoskeleton organization | GO:0051493 | over | 0.041 |
| cell death | GO:0008219 | under | 0.0412 |
| purine ribonucleotide metabolic process | GO:0009150 | under | 0.0417 |
| chloride transport | GO:0006821 | over | 0.0421 |
| inorganic anion transport | GO:0015698 | over | 0.0421 |
| embryo implantation | GO:0007566 | under | 0.0426 |
| cellular response to interferon-gamma | GO:0071346 | over | 0.0431 |
| response to interferon-gamma | GO:0034341 | over | 0.0431 |
| receptor-mediated endocytosis | GO:0006898 | over | 0.0432 |
| nitrogen compound metabolic process | GO:0006807 | under | 0.0437 |
| sterol transport | GO:0015918 | over | 0.0437 |
| double-strand break repair via classical nonhomologous end joining | GO:0097680 | under | 0.0437 |
| cholesterol transport | GO:0030301 | over | 0.0437 |
| phosphate-containing compound metabolic process | GO:0006796 | under | 0.0441 |
| phenol-containing compound biosynthetic process | GO:0046189 | over | 0.0441 |
| endothelial cell development | GO:0001885 | over | 0.045 |
| pyruvate metabolic process | GO:0006090 | under | 0.0451 |
| covalent chromatin modification | GO:0016569 | under | 0.0451 |
| single strand break repair | GO:0000012 | under | 0.0455 |
| regulation of actin filament-based movement | GO:1903115 | over | 0.0458 |
| regulation of cardiac muscle cell contraction | GO:0086004 | over | 0.0458 |
| plasma membrane organization | GO:0007009 | over | 0.0461 |
| positive regulation of protein localization to cell surface | GO:2000010 | over | 0.0466 |
| protein palmitoylation | GO:0018345 | under | 0.0476 |
| purine ribonucleotide biosynthetic process | GO:0009152 | under | 0.0481 |
| phosphorus metabolic process | GO:0006793 | under | 0.0484 |
| regulation of viral transcription | GO:0046782 | over | 0.0494 |
| regulation of peptidyl-serine phosphorylation | GO:0033135 | under | 0.0495 |
| COPII-coated vesicle budding | GO:0090114 | under | 0.0497 |

### Supplementary Table S3

| GO Term_ID | GO Term Pathway: Neonatal CSF-1 Stimulation | Fisher pvalue | GO Term_ID | GO Term Pathway: Neonatal IL-34 Stimulation | Fisher pvalue |
|---|---|---|---|---|---|
| GO:0044763 | single-organism cellular process | 1.50E-06 | GO:0006952 | defense response | 3.40E-08 |
| GO:0009888 | tissue development | 2.40E-06 | GO:0002376 | immune system process | 4.70E-07 |
| GO:0009653 | anatomical structure morphogenesis | 6.10E-06 | GO:0009605 | response to external stimulus | 1.60E-06 |
| GO:0044699 | single-organism process | 2.10E-05 | GO:0042221 | response to chemical | 3.40E-06 |
| GO:0048608 | reproductive structure development | 2.30E-05 | GO:0006955 | immune response | 3.70E-06 |
| GO:0061458 | reproductive system development | 2.30E-05 | GO:0050896 | response to stimulus | 3.70E-06 |
| GO:0060429 | epithelium development | 4.00E-05 | GO:0051716 | cellular response to stimulus | 7.80E-06 |
| GO:0007548 | sex differentiation | 4.90E-05 | GO:0051707 | response to other organism | 2.10E-05 |
| GO:0048513 | animal organ development | 8.10E-05 | GO:0043207 | response to external biotic stimulus | 2.10E-05 |
| GO:0009887 | animal organ morphogenesis | 0.00011 | GO:0001816 | cytokine production | 2.20E-05 |
| GO:0030855 | epithelial cell differentiation | 0.00019 | GO:0032101 | regulation of response to external stimu... | 2.50E-05 |
| GO:0008284 | positive regulation of cell proliferatio... | 0.00021 | GO:0009617 | response to bacterium | 2.50E-05 |
| GO:0050673 | epithelial cell proliferation | 0.00021 | GO:0009607 | response to biotic stimulus | 4.30E-05 |
| GO:0050679 | positive regulation of epithelial cell p... | 0.00023 | GO:0045321 | leukocyte activation | 4.30E-05 |
| GO:0032091 | negative regulation of protein binding | 0.00023 | GO:0070887 | cellular response to chemical stimulus | 4.70E-05 |
| GO:0044767 | single-organism developmental process | 0.00025 | GO:1904018 | positive regulation of vasculature devel... | 5.60E-05 |
| GO:0050678 | regulation of epithelial cell proliferat... | 0.0003 | GO:0045087 | innate immune response | 7.90E-05 |
| GO:0008406 | gonad development | 0.00035 | GO:0001775 | cell activation | 8.20E-05 |
| GO:0045137 | development of primary sexual characteri... | 0.00035 | GO:0030335 | positive regulation of cell migration | 9.20E-05 |
| GO:0032502 | developmental process | 0.00035 | GO:0002250 | adaptive immune response | 9.60E-05 |
| GO:0032963 | collagen metabolic process | 0.00039 | GO:0046649 | lymphocyte activation | 0.00013 |
| GO:0044259 | multicellular organismal macromolecule m... | 0.00039 | GO:0002237 | response to molecule of bacterial origin | 0.00013 |
| GO:0048856 | anatomical structure development | 0.00044 | GO:0010033 | response to organic substance | 0.00014 |
| GO:0010646 | regulation of cell communication | 0.00046 | GO:0007165 | signal transduction | 0.00016 |
| GO:0023051 | regulation of signaling | 0.00051 | GO:0035296 | regulation of tube diameter | 0.0002 |
| GO:0001568 | blood vessel development | 0.00055 | GO:0097746 | regulation of blood vessel diameter | 0.0002 |
| GO:0009966 | regulation of signal transduction | 0.00066 | GO:2000147 | positive regulation of cell motility | 0.00021 |
| GO:0048514 | blood vessel morphogenesis | 0.00071 | GO:0051240 | positive regulation of multicellular org... | 0.00024 |
| GO:0051716 | cellular response to stimulus | 0.00073 | GO:0040017 | positive regulation of locomotion | 0.00027 |
| GO:0003006 | developmental process involved in reprod... | 0.00079 | GO:0060326 | cell chemotaxis | 0.00031 |
| GO:0007049 | cell cycle | 0.0008 | GO:0045766 | positive regulation of angiogenesis | 0.00033 |
| GO:0001944 | vasculature development | 0.00083 | GO:0043269 | regulation of ion transport | 0.00041 |
| GO:0048732 | gland development | 0.00088 | GO:0051272 | positive regulation of cellular componen... | 0.00045 |
| GO:0043393 | regulation of protein binding | 0.00088 | GO:1901342 | regulation of vasculature development | 0.00047 |
| GO:0008283 | cell proliferation | 0.00091 | GO:0050880 | regulation of blood vessel size | 0.00049 |
| GO:0048729 | tissue morphogenesis | 0.00091 | GO:0035150 | regulation of tube size | 0.00049 |
| GO:0007155 | cell adhesion | 0.00098 | GO:0032496 | response to lipopolysaccharide | 0.00051 |
| GO:0022610 | biological adhesion | 0.00098 | GO:0097755 | positive regulation of blood vessel diam... | 0.00051 |
| GO:0072358 | cardiovascular system development | 0.00101 | GO:0048514 | blood vessel morphogenesis | 0.00053 |
| GO:0065008 | regulation of biological quality | 0.00103 | GO:0007154 | cell communication | 0.00056 |
| GO:0048583 | regulation of response to stimulus | 0.00107 | GO:0001525 | angiogenesis | 0.00068 |
| GO:0044707 | single-multicellular organism process | 0.00108 | GO:0044700 | single organism signaling | 0.00075 |
| GO:0046660 | female sex differentiation | 0.00109 | GO:0006935 | chemotaxis | 0.00079 |
| GO:0016477 | cell migration | 0.00111 | GO:0042330 | taxis | 0.00079 |
| GO:0007275 | multicellular organism development | 0.00116 | GO:0023052 | signaling | 0.00082 |
| GO:0009790 | embryo development | 0.00134 | GO:0034097 | response to cytokine | 0.00087 |
| GO:0006259 | DNA metabolic process | 0.00134 | GO:0033993 | response to lipid | 0.00088 |
| GO:0001890 | placenta development | 0.00135 | GO:0051239 | regulation of multicellular organismal p... | 0.00091 |
| GO:0007267 | cell-cell signaling | 0.00147 | GO:0050900 | leukocyte migration | 0.00099 |
| GO:0048646 | anatomical structure formation involved ... | 0.00156 | GO:0050920 | regulation of chemotaxis | 0.001 |
| GO:0051674 | localization of cell | 0.00168 | GO:0006950 | response to stress | 0.00113 |
| GO:0048870 | cell motility | 0.00168 | GO:0048518 | positive regulation of biological proces... | 0.00127 |
| GO:0048869 | cellular developmental process | 0.00173 | GO:0006954 | inflammatory response | 0.00129 |
| GO:0001763 | morphogenesis of a branching structure | 0.00178 | GO:0032103 | positive regulation of response to exter... | 0.00133 |
| GO:0009893 | positive regulation of metabolic process | 0.00188 | GO:0001568 | blood vessel development | 0.00133 |
| GO:0030334 | regulation of cell migration | 0.00196 | GO:0002694 | regulation of leukocyte activation | 0.00134 |
| GO:0048731 | system development | 0.00203 | GO:0048583 | regulation of response to stimulus | 0.00139 |
| GO:1902531 | regulation of intracellular signal trans... | 0.00214 | GO:2000378 | negative regulation of reactive oxygen s... | 0.00142 |
| GO:0044700 | single organism signaling | 0.00216 | GO:0032879 | regulation of localization | 0.00163 |
| GO:0023052 | signaling | 0.00216 | GO:0050865 | regulation of cell activation | 0.00165 |
| GO:0080134 | regulation of response to stress | 0.00232 | GO:0015850 | organic hydroxy compound transport | 0.00167 |
| GO:0035556 | intracellular signal transduction | 0.00234 | GO:0006936 | muscle contraction | 0.00167 |
| GO:0030324 | lung development | 0.00236 | GO:0030334 | regulation of cell migration | 0.0017 |
| GO:0023014 | signal transduction by protein phosphory... | 0.00236 | GO:0042108 | positive regulation of cytokine biosynth... | 0.00184 |
| GO:0000165 | MAPK cascade | 0.00236 | GO:0055088 | lipid homeostasis | 0.00184 |
| GO:2000145 | regulation of cell motility | 0.00236 | GO:0003018 | vascular process in circulatory system | 0.00184 |
| GO:0072676 | lymphocyte migration | 0.00236 | GO:0043270 | positive regulation of ion transport | 0.00192 |
| GO:0044236 | multicellular organism metabolic process | 0.00236 | GO:0051050 | positive regulation of transport | 0.00194 |
| GO:0030323 | respiratory tube development | 0.00236 | GO:0001817 | regulation of cytokine production | 0.00202 |
| GO:0050896 | response to stimulus | 0.00245 | GO:0031347 | regulation of defense response | 0.00203 |
| GO:0006811 | ion transport | 0.00249 | GO:0001944 | vasculature development | 0.00235 |
| GO:0051100 | negative regulation of binding | 0.00263 | GO:0044707 | single-multicellular organism process | 0.0024 |
| GO:0031325 | positive regulation of cellular metaboli... | 0.00297 | GO:0008283 | cell proliferation | 0.00251 |
| GO:0032501 | multicellular organismal process | 0.00314 | GO:0030595 | leukocyte chemotaxis | 0.00255 |
| GO:0051270 | regulation of cellular component movemen... | 0.00335 | GO:0048771 | tissue remodeling | 0.00255 |
| GO:0006260 | DNA replication | 0.00338 | GO:0072593 | reactive oxygen species metabolic proces... | 0.00256 |

| | | | | | |
|---|---|---|---|---|---|
| GO:0006928 | movement of cell or subcellular componen... | 0.00341 | GO:0051270 | regulation of cellular component movemen... | 0.00287 |
| GO:0072359 | circulatory system development | 0.00341 | GO:0006911 | phagocytosis | 0.00304 |
| GO:0007154 | cell communication | 0.0036 | GO:0032653 | regulation of interleukin-10 production | 0.00304 |
| GO:0006950 | response to stress | 0.00362 | GO:0050921 | positive regulation of chemotaxis | 0.00304 |
| GO:0051098 | regulation of binding | 0.00365 | GO:0042632 | cholesterol homeostasis | 0.00304 |
| GO:0007160 | cell-matrix adhesion | 0.00375 | GO:0030301 | cholesterol transport | 0.00304 |
| GO:0040012 | regulation of locomotion | 0.00396 | GO:0015918 | sterol transport | 0.00304 |
| GO:0040011 | locomotion | 0.00406 | GO:0010324 | membrane invagination | 0.00304 |
| GO:0045944 | positive regulation of transcription fro... | 0.00437 | GO:0055092 | sterol homeostasis | 0.00304 |
| GO:0043583 | ear development | 0.00437 | GO:0032613 | interleukin-10 production | 0.00304 |
| GO:1900371 | regulation of purine nucleotide biosynth... | 0.00437 | GO:0099024 | plasma membrane invagination | 0.00304 |
| GO:0030808 | regulation of nucleotide biosynthetic pr... | 0.00437 | GO:0001819 | positive regulation of cytokine producti... | 0.00309 |
| GO:0060541 | respiratory system development | 0.00437 | GO:1903409 | reactive oxygen species biosynthetic pro... | 0.0031 |
| GO:0030335 | positive regulation of cell migration | 0.00452 | GO:0042035 | regulation of cytokine biosynthetic proc... | 0.0031 |
| GO:2000147 | positive regulation of cell motility | 0.00452 | GO:0032501 | multicellular organismal process | 0.00315 |
| GO:0006974 | cellular response to DNA damage stimulus | 0.00465 | GO:0044699 | single-organism process | 0.00361 |
| GO:0007165 | signal transduction | 0.00503 | GO:1901700 | response to oxygen-containing compound | 0.00372 |
| GO:0006357 | regulation of transcription from RNA pol... | 0.00524 | GO:0030098 | lymphocyte differentiation | 0.00373 |
| GO:0045892 | negative regulation of transcription | 0.00532 | GO:0002696 | positive regulation of leukocyte activat... | 0.00373 |
| GO:0061138 | morphogenesis of a branching epithelium | 0.00542 | GO:0001818 | negative regulation of cytokine producti... | 0.00373 |
| GO:0010604 | positive regulation of macromolecule met... | 0.00547 | GO:2000377 | regulation of reactive oxygen species me... | 0.00373 |
| GO:0040017 | positive regulation of locomotion | 0.00554 | GO:2000145 | regulation of cell motility | 0.00374 |
| GO:0051272 | positive regulation of cellular componen... | 0.00554 | GO:0071345 | cellular response to cytokine stimulus | 0.00376 |
| GO:0051241 | negative regulation of multicellular org... | 0.00567 | GO:0072358 | cardiovascular system development | 0.00394 |
| GO:0031589 | cell-substrate adhesion | 0.00581 | GO:1901701 | cellular response to oxygen-containing c... | 0.00433 |
| GO:0035295 | tube development | 0.00585 | GO:0007166 | cell surface receptor signaling pathway | 0.00437 |
| GO:1903507 | negative regulation of nucleic acid-temp... | 0.00601 | GO:0007186 | G-protein coupled receptor signaling pat... | 0.00439 |
| GO:0000902 | cell morphogenesis | 0.00621 | GO:0001667 | ameboidal-type cell migration | 0.00439 |
| GO:0042127 | regulation of cell proliferation | 0.00711 | GO:0051049 | regulation of transport | 0.00464 |
| GO:1902532 | negative regulation of intracellular sig... | 0.00722 | GO:0050867 | positive regulation of cell activation | 0.00468 |
| GO:0043408 | regulation of MAPK cascade | 0.00722 | GO:0008203 | cholesterol metabolic process | 0.00487 |
| GO:1902679 | negative regulation of RNA biosynthetic ... | 0.0076 | GO:0071453 | cellular response to oxygen levels | 0.00487 |
| GO:0006270 | DNA replication initiation | 0.00776 | GO:0016125 | sterol metabolic process | 0.00487 |
| GO:0050806 | positive regulation of synaptic transmis... | 0.00776 | GO:0042107 | cytokine metabolic process | 0.00487 |
| GO:0008585 | female gonad development | 0.00776 | GO:0042089 | cytokine biosynthetic process | 0.00487 |
| GO:0008347 | glial cell migration | 0.00776 | GO:1902652 | secondary alcohol metabolic process | 0.00487 |
| GO:0048247 | lymphocyte chemotaxis | 0.00776 | GO:0003012 | muscle system process | 0.00498 |
| GO:0045737 | positive regulation of cyclin-dependent ... | 0.00776 | GO:0040012 | regulation of locomotion | 0.00499 |
| GO:0046545 | development of primary female sexual cha... | 0.00776 | GO:0006811 | ion transport | 0.00502 |
| GO:1904031 | positive regulation of cyclin-dependent ... | 0.00776 | GO:0071219 | cellular response to molecule of bacteri... | 0.00528 |
| GO:0001822 | kidney development | 0.00779 | GO:0048871 | multicellular organismal homeostasis | 0.00528 |
| GO:0050804 | modulation of synaptic transmission | 0.00779 | GO:0042110 | T cell activation | 0.00544 |
| GO:0030154 | cell differentiation | 0.00795 | GO:0002521 | leukocyte differentiation | 0.00544 |
| GO:0001525 | angiogenesis | 0.00812 | GO:1902041 | regulation of extrinsic apoptotic signal... | 0.0056 |
| GO:0051253 | negative regulation of RNA metabolic pro... | 0.00852 | GO:0042127 | regulation of cell proliferation | 0.00568 |
| GO:2000113 | negative regulation of cellular macromol... | 0.0087 | GO:0051249 | regulation of lymphocyte activation | 0.00572 |
| GO:0071900 | regulation of protein serine/threonine k... | 0.0089 | GO:0044765 | single-organism transport | 0.00573 |
| GO:0098916 | anterograde trans-synaptic signaling | 0.0092 | GO:0016477 | cell migration | 0.00619 |
| GO:0099536 | synaptic signaling | 0.0092 | GO:0048522 | positive regulation of cellular process | 0.00646 |
| GO:0099537 | trans-synaptic signaling | 0.0092 | GO:0051704 | multi-organism process | 0.0065 |
| GO:0007268 | chemical synaptic transmission | 0.0092 | GO:0002252 | immune effector process | 0.00701 |
| GO:0006366 | transcription from RNA polymerase II pro... | 0.00931 | GO:0051047 | positive regulation of secretion | 0.0071 |
| GO:0022603 | regulation of anatomical structure morph... | 0.00932 | GO:0045765 | regulation of angiogenesis | 0.00726 |
| GO:0010647 | positive regulation of cell communicatio... | 0.00952 | GO:0097529 | myeloid leukocyte migration | 0.00727 |
| GO:0023056 | positive regulation of signaling | 0.00952 | GO:0048584 | positive regulation of response to stimu... | 0.00742 |
| GO:0009719 | response to endogenous stimulus | 0.01036 | GO:0022610 | biological adhesion | 0.00779 |
| GO:0015711 | organic anion transport | 0.01079 | GO:0002682 | regulation of immune system process | 0.00871 |
| GO:0072001 | renal system development | 0.01079 | GO:0002687 | positive regulation of leukocyte migrati... | 0.00927 |
| GO:0007507 | heart development | 0.01082 | GO:0050672 | negative regulation of lymphocyte prolif... | 0.00927 |
| GO:0065009 | regulation of molecular function | 0.01084 | GO:0071456 | cellular response to hypoxia | 0.00927 |
| GO:0016043 | cellular component organization | 0.0109 | GO:0032945 | negative regulation of mononuclear cell ... | 0.00927 |
| GO:0000122 | negative regulation of transcription fro... | 0.01115 | GO:0070664 | negative regulation of leukocyte prolife... | 0.00927 |
| GO:0044702 | single organism reproductive process | 0.01115 | GO:0006937 | regulation of muscle contraction | 0.00939 |
| GO:2000027 | regulation of organ morphogenesis | 0.01124 | GO:0046849 | bone remodeling | 0.00939 |
| GO:0030856 | regulation of epithelial cell differenti... | 0.01124 | GO:0031638 | zymogen activation | 0.00939 |
| GO:0006261 | DNA-dependent DNA replication | 0.01124 | GO:0001893 | maternal placenta development | 0.00939 |
| GO:0051301 | cell division | 0.01172 | GO:0043266 | regulation of potassium ion transport | 0.00939 |
| GO:1901137 | carbohydrate derivative biosynthetic pro... | 0.01172 | GO:0042310 | vasoconstriction | 0.00939 |
| GO:0009260 | ribonucleotide biosynthetic process | 0.01188 | GO:0033344 | cholesterol efflux | 0.00939 |
| GO:0051179 | localization | 0.01195 | GO:1903427 | negative regulation of reactive oxygen s... | 0.00939 |
| GO:0022414 | reproductive process | 0.01271 | GO:0097756 | negative regulation of blood vessel diam... | 0.00939 |
| GO:0000003 | reproduction | 0.01271 | GO:0060135 | maternal process involved in female preg... | 0.00939 |
| GO:0009891 | positive regulation of biosynthetic proc... | 0.01281 | GO:0010038 | response to metal ion | 0.00972 |
| GO:0002009 | morphogenesis of an epithelium | 0.01303 | GO:0071216 | cellular response to biotic stimulus | 0.00972 |
| GO:0001667 | ameboidal-type cell migration | 0.01433 | GO:0001763 | morphogenesis of a branching structure | 0.01038 |
| GO:0060993 | kidney morphogenesis | 0.01443 | GO:0071706 | tumor necrosis factor superfamily cytoki... | 0.01038 |
| GO:0007492 | endoderm development | 0.01443 | GO:1903555 | regulation of tumor necrosis factor supe... | 0.01038 |

| | | | | | |
|---|---|---|---|---|---|
| GO:0006919 | activation of cysteine-type endopeptidas... | 0.01443 | GO:0031349 | positive regulation of defense response | 0.01077 |
| GO:0006949 | syncytium formation | 0.01443 | GO:0006869 | lipid transport | 0.01077 |
| GO:0014911 | positive regulation of smooth muscle cel... | 0.01443 | GO:0006629 | lipid metabolic process | 0.01134 |
| GO:0030857 | negative regulation of epithelial cell d... | 0.01443 | GO:0061061 | muscle structure development | 0.01233 |
| GO:0001892 | embryonic placenta development | 0.01443 | GO:0007155 | cell adhesion | 0.01263 |
| GO:0043407 | negative regulation of MAP kinase activi... | 0.01443 | GO:0051094 | positive regulation of developmental pro... | 0.01263 |
| GO:0000768 | syncytium formation by plasma membrane f... | 0.01443 | GO:0042113 | B cell activation | 0.01273 |
| GO:0048146 | positive regulation of fibroblast prolif... | 0.01443 | GO:0051674 | localization of cell | 0.01328 |
| GO:0001676 | long-chain fatty acid metabolic process | 0.01443 | GO:0048870 | cell motility | 0.01328 |
| GO:0010810 | regulation of cell-substrate adhesion | 0.01451 | GO:0072359 | circulatory system development | 0.01375 |
| GO:0001655 | urogenital system development | 0.01451 | GO:0046903 | secretion | 0.01417 |
| GO:0045934 | negative regulation of nucleobase-contai... | 0.01451 | GO:0006941 | striated muscle contraction | 0.01423 |
| GO:0006820 | anion transport | 0.01507 | GO:0031644 | regulation of neurological system proces... | 0.01423 |
| GO:0043405 | regulation of MAP kinase activity | 0.01507 | GO:0001894 | tissue homeostasis | 0.01423 |
| GO:0046390 | ribose phosphate biosynthetic process | 0.01507 | GO:0050680 | negative regulation of epithelial cell p... | 0.01423 |
| GO:0050727 | regulation of inflammatory response | 0.01507 | GO:0061138 | morphogenesis of a branching epithelium | 0.01423 |
| GO:0032989 | cellular component morphogenesis | 0.01522 | GO:0071222 | cellular response to lipopolysaccharide | 0.01429 |
| GO:0098602 | single organism cell adhesion | 0.01574 | GO:0019882 | antigen processing and presentation | 0.01429 |
| GO:0032101 | regulation of response to external stimu... | 0.01592 | GO:0071310 | cellular response to organic substance | 0.01587 |
| GO:0010558 | negative regulation of macromolecule bio... | 0.01592 | GO:0007159 | leukocyte cell-cell adhesion | 0.01607 |
| GO:0071840 | cellular component organization or bioge... | 0.01613 | GO:1903037 | regulation of leukocyte cell-cell adhesi... | 0.01607 |
| GO:0051093 | negative regulation of developmental pro... | 0.01632 | GO:0098542 | defense response to other organism | 0.01607 |
| GO:0002376 | immune system process | 0.01633 | GO:0050878 | regulation of body fluid levels | 0.01634 |
| GO:0015850 | organic hydroxy compound transport | 0.01636 | GO:0034220 | ion transmembrane transport | 0.01646 |
| GO:0051302 | regulation of cell division | 0.01636 | GO:0001937 | negative regulation of endothelial cell ... | 0.01736 |
| GO:0072006 | nephron development | 0.01636 | GO:0032733 | positive regulation of interleukin-10 pr... | 0.01736 |
| GO:0072009 | nephron epithelium development | 0.01636 | GO:0048469 | cell maturation | 0.01736 |
| GO:0048754 | branching morphogenesis of an epithelial... | 0.01636 | GO:0071804 | cellular potassium ion transport | 0.01736 |
| GO:0048519 | negative regulation of biological proces... | 0.01715 | GO:0071805 | potassium ion transmembrane transport | 0.01736 |
| GO:0033554 | cellular response to stress | 0.01767 | GO:0051930 | regulation of sensory perception of pain | 0.01736 |
| GO:0051254 | positive regulation of RNA metabolic pro... | 0.01768 | GO:0051931 | regulation of sensory perception | 0.01736 |
| GO:0032103 | positive regulation of response to exter... | 0.0188 | GO:0034138 | toll-like receptor 3 signaling pathway | 0.01736 |
| GO:0002064 | epithelial cell development | 0.01899 | GO:0007589 | body fluid secretion | 0.01736 |
| GO:0060485 | mesenchyme development | 0.01899 | GO:0061041 | regulation of wound healing | 0.01736 |
| GO:0048518 | positive regulation of biological proces... | 0.01905 | GO:0044710 | single-organism metabolic process | 0.01784 |
| GO:0051094 | positive regulation of developmental pro... | 0.01946 | GO:0008284 | positive regulation of cell proliferatio... | 0.0179 |
| GO:0051173 | positive regulation of nitrogen compound... | 0.01987 | GO:1902578 | single-organism localization | 0.01804 |
| GO:0031328 | positive regulation of cellular biosynth... | 0.02048 | GO:0009887 | animal organ morphogenesis | 0.01881 |
| GO:0045935 | positive regulation of nucleobase-contai... | 0.02048 | GO:0022407 | regulation of cell-cell adhesion | 0.01922 |
| GO:1902578 | single-organism localization | 0.02062 | GO:0010035 | response to inorganic substance | 0.01922 |
| GO:0044765 | single-organism transport | 0.02081 | GO:0007517 | muscle organ development | 0.02035 |
| GO:0030099 | myeloid cell differentiation | 0.02092 | GO:0071241 | cellular response to inorganic substance | 0.02059 |
| GO:0098609 | cell-cell adhesion | 0.02144 | GO:0000422 | mitophagy | 0.02059 |
| GO:0006281 | DNA repair | 0.02163 | GO:0010761 | fibroblast migration | 0.02059 |
| GO:0048522 | positive regulation of cellular process | 0.02211 | GO:0006813 | potassium ion transport | 0.02059 |
| GO:0050793 | regulation of developmental process | 0.02223 | GO:0002685 | regulation of leukocyte migration | 0.02059 |
| GO:0046942 | carboxylic acid transport | 0.02273 | GO:0036294 | cellular response to decreased oxygen le... | 0.02059 |
